# Nutritional vitamin E or plant extracts affect the immune response and mammary epithelium integrity during intramammary lipopolysaccharide challenge in early lactation

**DOI:** 10.1101/2025.09.18.677083

**Authors:** A. Corset, A. Remot, B. Graulet, J.F. Ricouleau, S. Philau, K. Reis-Santos, O. Dhumez, P. Germon, M. Boutinaud, A. Boudon

## Abstract

**Background:** Early lactation is a period at risk for mastitis. Our objective was to investigate the influence of nutritional vitamin E and plant extracts on redox and immune status at the systemic and local mammary levels during an inflammatory challenge in dairy cows. Thirty-six Holstein cows were placed in three groups: a ‘control’ group (n = 11) unsupplemented; a ‘vitamin E’ group (n = 13), supplemented with 3000 IU/d for 4 weeks before calving and 1000 IU/d for 4 weeks after calving; and a ‘plant extracts’ group (n = 14) supplemented with 10 g/d for 4 weeks before and after calving. Lipopolysaccharide (LPS) from *Escherichia coli* was infused into a one-udder quarter of the cows 5 weeks after calving. Blood and milk samples were collected before and 4, 7, 9, 28, and 76 h after the LPS infusion.

**Results:** In the plant extract group, antioxidant gene (*NQO1*), superoxide dismutase activity 4 hours after the LPS challenge and total antioxidant capacity were increased compared to the control group. In the vitamin E group, expression of the antioxidant gene (*SOD3*) and glutathione peroxidase activity were more elevated than in the control group. Systemic immunity appeared to be reduced by antioxidant supplementations, with a higher migration of classical neutrophils in the plant extract group and less ROS production of neutrophils in the vitamin E group. However, immune capacities were increased at the local level in classical neutrophils (ROS production and phagocytosis) in both the plant extract and vitamin E groups. Following the LPS infusion, supplementation did not reveal any differences in terms of rectal temperature, milk yield or milk cell counts, but it did have a positive effect on mammary epithelium integrity (lower milk Na^+^:K^+^ ratio and higher *OCLN* gene expression). In addition, the plant extracts stimulated the synthesis of milk constituents, with higher milk protein gene expressions (*LALBA*, *CSN3*, *CSN1S1*) and a tendency towards a higher lactose content in milk.

**Conclusion:** The LPS challenge showed that vitamin E or plant extracts enhanced the local inflammatory response while preserving systemic antioxidant capacities, integrity of the mammary gland and the composition of milk, and thus possibly contributing to preventing mammary infection in early lactating dairy cows.

## Background

Mastitis is defined as an inflammation of the udder, most frequently caused by bacteria; its incidence is highest at the start of lactation. Among other consequences, mastitis leads to a drop in milk yield and milk quality and an increase in somatic cell counts, resulting in higher financial costs, an impairment of animal welfare and significant antibiotic usage (1).

Nutritional strategies form part of the solutions available to prevent mastitis (2). Plasma vitamin E levels in healthy cows are higher than those in unhealthy cows (3). This is particularly true in cows with clinical mastitis, which tend to display lower plasma α-tocopherol levels than healthy cows (4). Hogan et al. (5) observed that 44% of cows with mastitis experienced a reduction in clinical mastitis symptoms when supplemented with vitamin E (5). Politis later showed a reduction in milk cell count and plasmin in milk, suggesting an effect on milk quality (6). Consistent with this, other studies have shown a regulation of blood neutrophils with vitamin E supplementation, lowering ROS production that could induce cellular and tissue damage (7, 8). Vitamin E has been shown to improve the migratory capacity of neutrophils by increasing urokinase synthesis (6), and to influence their function during the peripartum period (9). Vitamin E is also known to reduce oxidative stress when present in the blood, and to reduce MDA, but not to influence antioxidant capacities (FRAP) and reactive oxygen metabolites (ROM) (10).

Another nutritional strategy used to prevent mastitis may be to supplement the cows with plant extracts containing polyphenols which are known to have potential beneficial antioxidant properties. Some plant extracts have been shown to affect immune or redox status in different species (12–16). When bovine lymphocytes and monocytes were subjected to an *in vitro* phagocytosis test, *Silybum marianum* extracts were shown to increase phagocytosis (11). *Arctium lappa* administered to rats was able to reduce acetaminophen-induced liver hepatotoxicity and lower the MDA content (12). *White willow* has been found to have antioxidant functions in rabbits (13). *Sambucus nigra*, *Laurus nobilis*, and *Harpagophytum procumbens* extracts have been shown in the literature to have anti-inflammatory or antioxidant functions, but their extracts were more competent than their essential oils or isolated active ingredients (14–16). There is therefore a need to study the effect of a combination of plant extracts rather than a single molecule in dairy cows in order to prevent the damage associated with indexed mastitis.

The elimination of pathogens by immune cells can also trigger degradation of the blood-milk barrier, induce lesions in mammary epithelial tissue, and impact the immune, production and metabolic responses (17–19). Lipopolysaccharide (LPS) from Gram negative bacteria can be responsible for a severe inflammation (20). The injection of LPS thus represents a good model to induce sterile inflammation in the udder that mimics clinical mastitis, in order to evaluate the effects of vitamin E and plant extracts on the immune response and the prevention of mastitis. Our strategy was designed to evaluate how two months of nutritional supplementation with vitamin E or plant extracts could affect the ability of the cows to respond to an LPS-induced mammary inflammation, and then resolve the inflammation. To achieve this, we looked at redox and immune status indicators of the cows, both systemically and locally at the level of the mammary gland. The originality of our study also resided in our evaluation of mammary epithelium integrity during the LPS challenge, and the effects of dietary vitamin E or plant extract supplementations. Our hypothesis was that vitamin E and plant extracts could optimally modulate the LPS-induced inflammatory response at both the immune system and local levels, and affect antioxidant responses to prevent cellular and tissue damage.

## Results

### DMI, energy metabolism indicators, plasma and ingested vitamin contents during the LPS challenge and effects of nutritional strategy

Dry matter intake (DMI) was increased progressively between calving and the start of the LPS challenge, and then stabilized during the week of the challenge (*P* <0.001, Fig 1a). DMI was not affected by vitamin E and plant extracts (*P* = 0.45). Levels of ingested α-tocopherol were higher in the vitamin E group than in the control and plant extract groups (*P* <0.001, Fig 1b), but were not affected by the LPS challenge.

**Figure 1:**
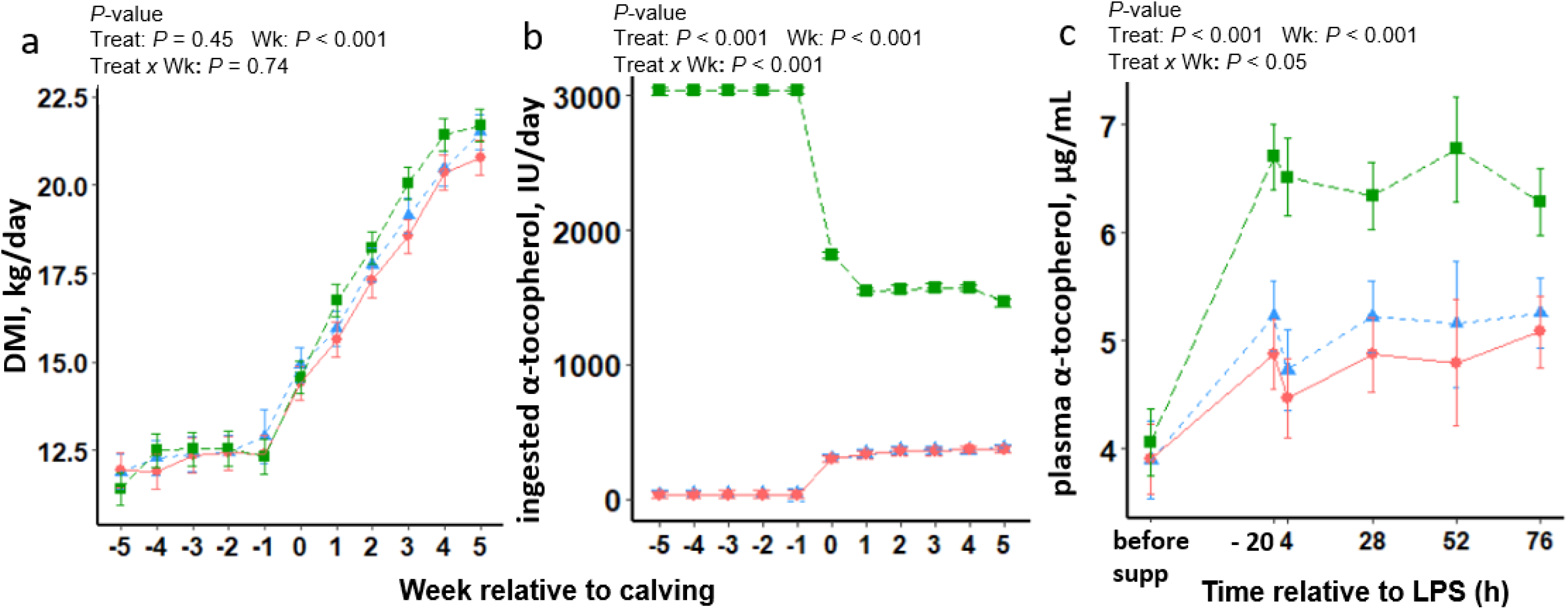
Dry matter intake (DMI), ingested α-tocopherol and plasma α-tocopherol concentrations in the control unsupplemented group (n = 11), vitamin E supplemented group (n = 13), and plant extract supplemented group (n = 12) of dairy cows before and during the LPS challenge in week 5 relative to calving. DMI in kg/d (**a**) and ingested α-tocopherol (α), in IU/d (**b**). Plasma α-tocopherol was measured before vitamin E supplementation and before the LPS challenge at -20 hours, and then during the LPS challenge in week 5 relative to calving, µg/mL (**c**). The cows were followed for between -5 and 5 weeks relative to calving. The control group is represented by a solid red line (□●□), the vitamin E group by a dotted green line (--▪--), and the plant extract group by a dotted blue line (- -▴- -). Adjusted means and SEM are represented and the data were analysed according to a mixed model. Significant differences between treatment (Treat) and time (Time) and their interaction (Treat x Time) are noted above the graphs.

As expected, plasma α-tocopherol levels were higher in the vitamin E group than in the control and plant extract groups from 20 days before calving until the end of the experiment (6.11 ± 0.23 *vs* 4.66 ± 0.25 and 4.92 ± 0.25 µg/ml) whereas they were not affected by the groups before supplementation was started (treatment x time interaction, *P* <0.03, Fig. 1c). Neither average levels nor the daily dynamics of plasma metabolites were affected by vitamin E or plant extract supplementation. Plasma non-esterified fatty acids (NEFA) and urea did not vary significantly during the LPS challenge (Supplementary Data Table S1, *P* >0.10), while glucose levels were higher at 4 h and 28 h after the LPS challenge than at any other time point (*P* <0.001). Plasma β-hydroxybutyrate (BHB) and calcium levels were lower at 4 h and 28 h after the LPS challenge than at any other time point (*P* <0.001). Inorganic phosphorus was lower 4h after the LPS challenge than at any other time point (*P* <0.001).

### Systemic and local redox status during the intramammary challenge and effects of nutritional strategy

Plasma reactive oxygen metabolite (ROM) concentrations were lower 4 h after the LPS challenge than at 76 h (*P* = 0.01, Fig. 2a). Plasma ferric reducing antioxidant power (FRAP) was not affected by the LPS challenge (time *P* >0.10) but tended to be higher in the plant extract group than among controls (63.44 ± 0.94 *vs* 60.44 ± 0.93 µg/mL) whatever the time point (*P* = 0.07, Fig. 2b). Plasma biological antioxidant potential (BAP) was reduced by LPS and gradually increased until 76 h after the challenge (*P* = 0.01, Fig. 2c), but was not affected by the vitamin E or plant extract treatments. GPx activity in plasma was higher at 28 h than at any other time point (*P* < 0.001, Fig. 2d), but was not affected by the vitamin E or plant extract treatments. At 76h after the LPS challenge, GPx activity in erythrocytes tented to be higher in the vitamin E group than in the plant extract group (3854 ± 303.4 *vs* 3182 ± 322.4 nmol/min/mL) (time *P* = 0.06, treatment × time, *P* = 0.07, Fig. 2e). Plasma SOD activity was lower at 28 h after than before and at 4 h after the LPS challenge (*P* = 0.01, Fig. 2f), but was not affected by the vitamin E or plant extract treatments. SOD activity in erythrocytes was lower at 28 h than at any other time point; it was higher in the plant extract group than in the control group and tended to be higher in the vitamin E group than in the control group (221.1 ± 11.27 *vs* 172.4 ± 11.18 and 186.9 ± 10.64 nmol/min/mL, respectively) (*P* = 0.01, Fig. 2g).

**Figure 2:**
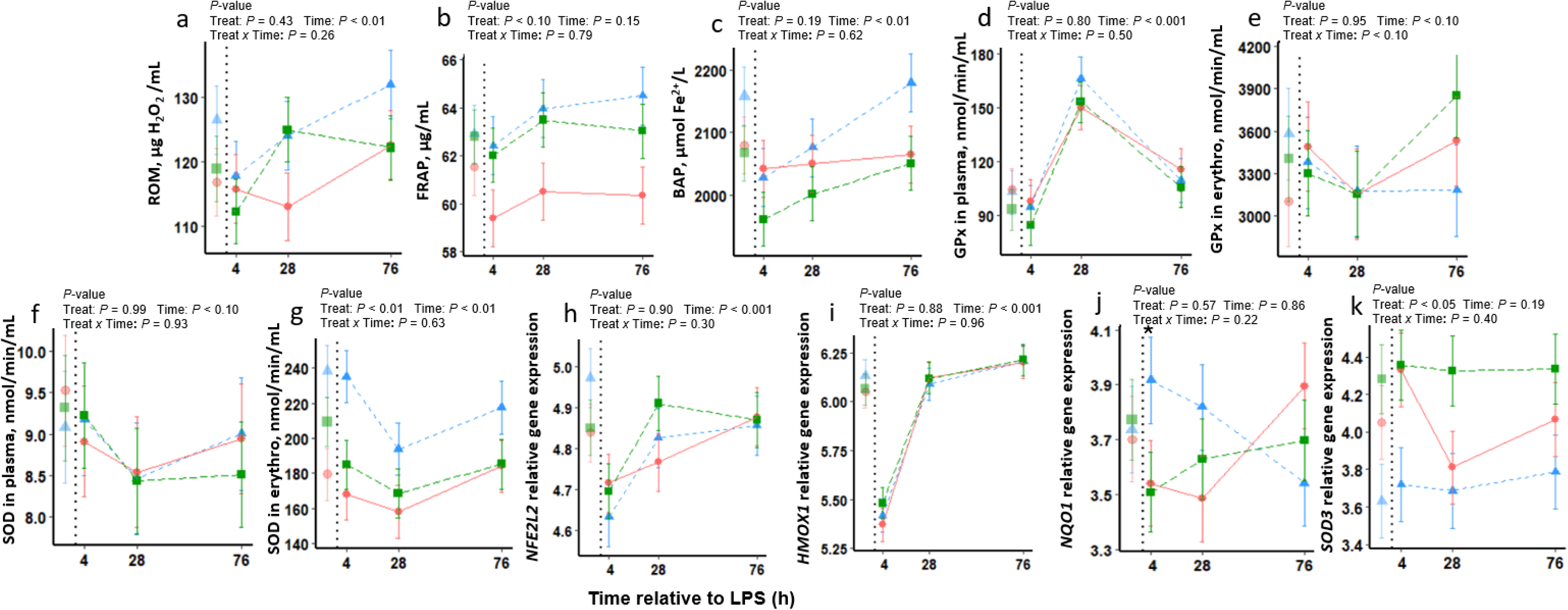
Biomarkers for the redox balance, enzymatic activity in plasma or erythrocytes and antioxidant gene expression in the control unsupplemented group (n = 11), vitamin E supplemented group (n = 13), and plant extract supplemented group (n = 12) of dairy cows before and during the LPS challenge (at 4 h, 28 h and 76h). Reactive oxygen metabolite (ROM), µg H_2_O_2_ /mL (**a**), ferric reducing antioxidant power (FRAP), µg/mL (**b**), biological antioxidant power (BAP), µmol Fe^2+^/L (**c**) in plasma. The antioxidant enzyme activity of glutathione peroxidase (GPx) was measured in plasma (**d**) and erythrocytes (**e**) in nmol/min/mL. The antioxidant enzyme activity of superoxide dismutase (SOD) was measured in plasma (**f**) and erythrocytes (**g**) in nmol/min/mL. Antioxidant gene expression in total blood was measured, *NFE2L2* (nuclear factor (erythroid-derived 2)-like 2) (**h**), *HMOX1* (heme oxygenase 1) (**i**), *NQO1* (NAD(P)H quinone dehydrogenase 1) (**j**), and SOD3 (superoxide dismutase 3) (**k**) in relative gene expression. The control group is represented as a solid red line (□●□), the vitamin E group as a dotted green line (--▪--), and the plant extract group as a dotted blue line (- -▴- -). Adjusted means and SEM are represented and the data were analysed according to a mixed model. Significant differences between treatment (Treat) and time (Time) and their interaction (Treat x Time) are noted above the graphs. Thresholds of significance were set at ∗ *P* <0.05

In the blood, the means of reference genes (*ACTB*, *YWHAZ*, and *RPLP0*) did not vary according to the treatment received (*P* = 0.70) (Supplementary Data Table S2). Expressions of the *NFE2L2* and *HMOX1* antioxidant genes were lower 4 h after the LPS challenge than at any other time point (*P* < 0.001, Fig. 2h and 2i) whereas that of *NQO1* was not affected by time relative to the LPS challenge (Fig 2j). Although *NQO1* antioxidant gene expression was not affected by the treatment x time interaction, Student’s t-tests showed that the level of *NQO1* expression was higher in the plant extract group than in the control group at 4 h and 28 h after the LPS challenge (*P* = 0.04 and *P* = 0.03) and tended to be higher in the plant extract group than in the vitamin E group at 4 h after the LPS challenge (Student’s t-test *P* = 0.06, Fig. 2j). The *SOD3* antioxidant gene was more strongly expressed in the vitamin E group than in the plant extract group during the LPS challenge (*P* = 0.04, Fig. 2k), whatever the time point.

In milk, the means of reference genes (*ACTB*, *YWHAZ*, and *PPIA*) did not vary according to treatment and parity (*P* >0.10) (Supplementary data Table S3). The expressions of some redox genes (*NFE2L2*, *TXNRD1, CAT, SOD1 and SOD3)* were lower at 4 h or 28 h after the LPS challenge than at any other time points (*P* < 0.001, Supplementary data Table S3) but they were not affected by the vitamin E or plant extract treatments.

### Systemic immune status during the intramammary challenge and nutritional strategy effects

Rectal temperature was higher at 4 h, 7 h and 9 h after the LPS challenge than before it, or at 28 h and 76 h after the challenge (*P* <0.001, Fig. 3a). Plasma cortisol levels were higher 4 h after the LPS challenge than at any other time point (*P* <0.001, Fig. 3b). Plasma haptoglobin levels were higher 28 h and 76 h than before, and 4 h after, the LPS challenge (*P* <0.001, Fig. 3c). The white blood cell and neutrophil counts were lower 4 h after the LPS challenge than at any other time point (*P* <0.001 and *P* <0.01, respectively) (Fig. 3d and 3e). The blood neutrophil count increased at 28 h after the challenge and then decreased (*P* <0.01, Fig. 3e). Blood monocyte and lymphocyte counts were lower at 4 h than at 28 h and 76 h after the challenge (*P* <0.001, Supplementary data Table S4). Treatment × time interactions were barely significant relative to those parameters. However, plasma cortisol was higher in the plant extract group than in the control group at 4 h and 28 h after the LPS challenge (Student’s t-test *P* = 0.04 for both time points). The white blood cell count was lower in the plant extract group than in the control group at 4 h after the LPS challenge (5.32 ± 0.77 *vs* 7.45 ± 0.77 x 10^9^/L, respectively) (Student’s t-test *P* = 0.04, Fig. 3d), while the blood neutrophil count was lower in the plant extract group than in the control group at 4 h after the LPS challenge (2.19 ± 0.44 *vs* 3.38 ± 0.44 x 10^9^/L) (Student’s t-test *P* = 0.04, Fig. 3e). Blood monocyte and lymphocyte counts were unaffected by the treatments (*P* = 0.70) (Supplementary data Table S4).

**Figure 3:**
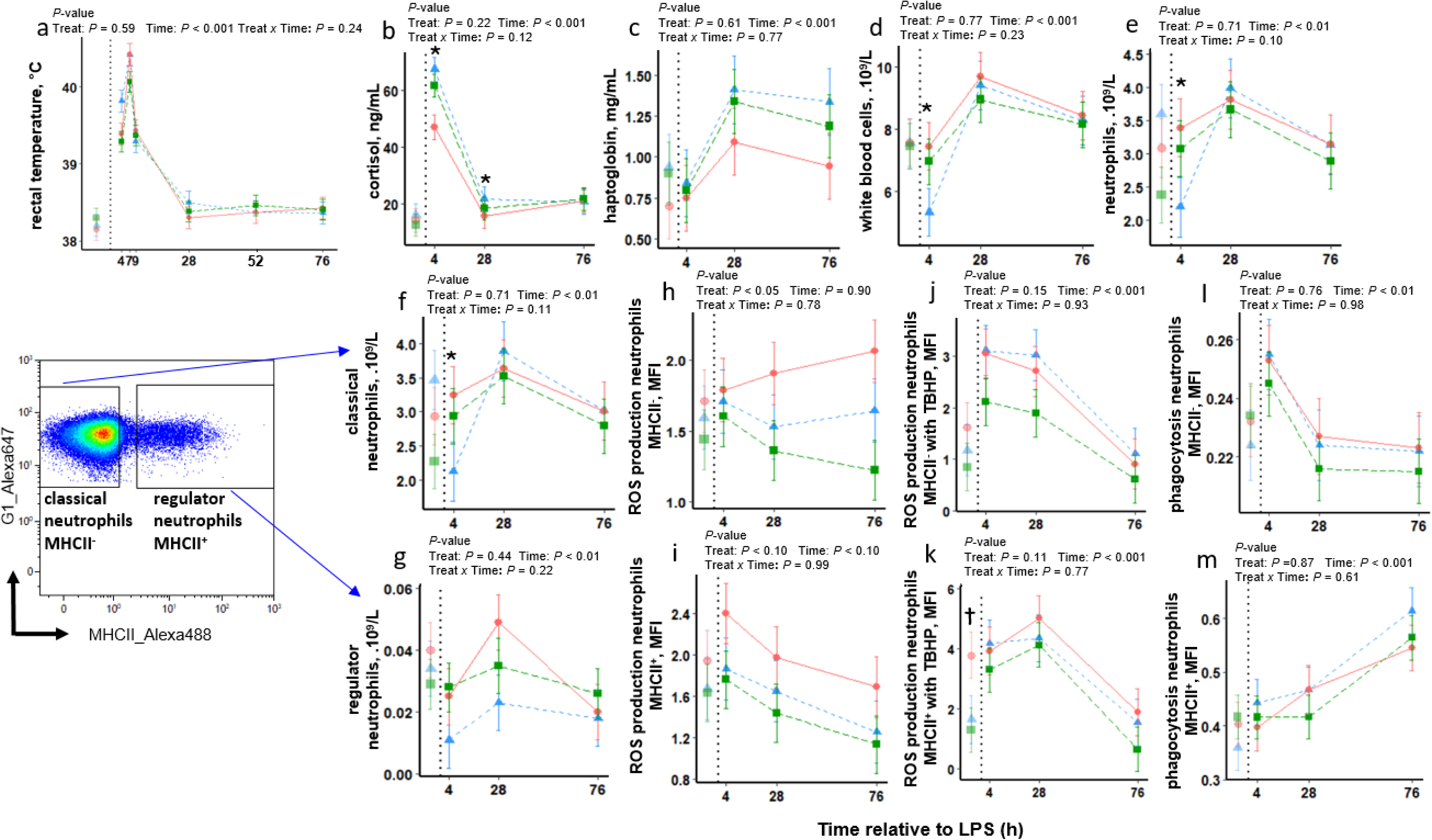
Rectal temperature, plasma haptoglobin, plasma cortisol, haematological profile and functional capacities of blood neutrophils in the control unsupplemented group (n = 11), vitamin E supplemented group (n = 13), and plant extract supplemented group (n = 12) of dairy cows before and during the LPS challenge (at 4 h, 7 h, 9 h, 28 h, 52 h and 76h). Rectal temperature in Celsius (**a**), cortisol, in ng/mL (**b**), haptoglobin in mg/mL (**c**), and haematological profile with white blood cells in 10^9^/L (**d**) and neutrophils in 10^9^/L (**e**). Neutrophil count measured by flow cytometry without class II major histocompatibility complex, MHCII**^-^** (**f**) and class II major histocompatibility complex neutrophils, MHCII**^+^** (**g**). *Ex vivo*, mean fluorescence intensity (MFI) measurement of intracellular ROS production in the two neutrophil subpopulations determined by flow cytometry without (**h**, **j**) or with (**k**, **l**) tert-butyl hydroperoxide (TBHP) stimulation. MFI measurement of *ex vivo* phagocytosis of the two neutrophil subpopulations (**m**, **n**). The control group is represented as a solid red line (□●□), the vitamin E group as a dotted green line (--▪--), and the plant extract group as a dotted blue line (- -▴- -). Adjusted means and SEM are represented and the data were analysed according to a mixed model. Significant differences between treatment (Treat) and time (Time) and their interaction (Treat x Time) are noted above the graphs. Thresholds of significance were set at ∗ *P* <0.05 and trends were noted at ^†^ *P* ≤0.10.

The classic neutrophil count (i.e. cells without the class II major histocompatibility complex (MHCII^-^) was higher 28 h after the LPS challenge than at any other time point (*P* = 0.01, Fig. 3f). Although classic MHCII^-^ neutrophils were not affected by the treatment × time interaction, Student’s t-tests showed that 4 h after the LPS challenge, their levels were lower in the plant extract group than in the control group (2.13 ± 0.43 *vs* 3.23 ± 0.42 x 10^9^/L, respectively) (*P* = 0.04, Fig. 3f). Like classic neutrophils, the numbers of regulatory neutrophils expressing the class II major histocompatibility complex (MHCII^+^) were lower at 4 h and 76 h than before, and 28 h after, the LPS challenge (*P* = 0.01) but without any treatment effect (*P* = 0.22) (Fig. 3g). Regarding the baseline level measured by flow cytometry without TBHP stimulation, the production of ROS by classic MHCII^-^ neutrophils did not vary as a function of time (*P* = 0.90, Fig. 3h) although ROS production by regulatory MHCII^+^ tended to increase at 4 h after the LPS challenge and then continuously decrease at 28 and 76 h thereafter (*P* = 0.05, Fig. 3i). Without TBHP stimulation, classic MHCII^-^ neutrophils and regulatory MHCII^+^neutrophils produced, or tended to produce, less ROS in the vitamin E group than in the control group, whatever the time point (*P* = 0.04 and *P* = 0.07 for MHCII^-^ and MHCII^+^, respectively, Fig 3h and 3i). With TBHP stimulation, ROS production by regulatory MHCII^+^ neutrophils tended to be lower in the vitamin E group than in the control group, but only before the LPS challenge (Student’s t-test *P* = 0.09, Fig. 3k). After the challenge, ROS production by classic MHCII^-^ neutrophils and regulatory MHCII^+^ neutrophils increased at 4 h, then started to gradually decline 28 h after the challenge (*P* <0.001, Fig. 3j and Fig. 3k). Classic MHCII^-^ neutrophils phagocytized more at 4 h after the LPS challenge than at any other time point (*P* <0.001, Fig. 3l) while regulatory MHCII^+^ neutrophils phagocytized more 76 h after the LPS challenge than at the other time points (*P* <0.001 Fig. 3m). Phagocytosis by neutrophils was not affected by the vitamin E or plant extract treatments. Blood monocyte capacities evaluated in terms of their percentage of ROS production and phagocytosis did not display any effects of treatment, time or time X treatment interaction (Supplementary data Table S5).

The transcriptomic profile of 79 amplified genes in blood samples showed that many antioxidant genes (*NFE2L2*, *HMOX1, GPX1* etc.) and immune stimulation genes (*TAP*, *CD14*, *NF*_κ_*B* etc.) are down-regulated at the start of inflammation (4 h) and up-regulated before and at the end of inflammation (-20 h and 76h, Fig. 4a). At the same time points, cytokine genes (*IL1A*, *ILB*, *CXCL8* etc.), complement factor (*CCR1*, *CCR7* etc.) or migration genes (*SELL*, *MMP9* etc.), displayed stronger expression at 4 h after the LPS challenge than at any other time point (Fig. 4a). The expression levels of genes involved in inflammation and not affected by the supplementations are shown in the Supplementary Data, at Table S2 and Fig. S1. Whatever the time relative to the LPS challenge, the *STAT5A* gene was up-regulated in the plant extract group compared to the control and vitamin E groups (fold increases of 0.18 and 0.02, respectively, *P* = 0.03, Fig. 4b). Before the LPS challenge, *CXCL2* was up-regulated in the plant extract group compared to the control group (1.73-fold increase, treatment x time interaction, *P* = 0.02, Fig. 4b) and *CCL20*, *CD80*, *TAP* genes tended to be up-regulated (fold increase of between 20.9 and 0.25, *P* <0.10). At the same time, the *FABP3* gene also tended to be up-regulated in the vitamin E and plant extract groups compared to the control group (+44% and +40% respectively, treatment x time interaction, *P* = 0.08) (Fig. 4b), and the *SOCS3* gene tended to be up-regulated in the plant extract group compared to the vitamin E group (0.44-fold increase, Student’s t-test, *P* = 0.08, Fig. 4b). At 4 h after the LPS challenge, the expression of a number of genes was disturbed: the *MMP14* gene was down-regulated in the plant extract group compared to the vitamin E and control groups (-73% and - 83%, respectively, *P* = 0.05); the *CCR7, IKZF1* and *CXCR4* genes were up-regulated in the plant extract group compared to the control group (fold increase of between 0.13 and 0.59, Student’s t-test, *P* <0.05); the *CCL5*, *ADGRE1*, *SOCS3* genes tended to be up-regulated in the plant extract group compared to the control group (fold increase of between 0.37 and 1.17, Student’s t-test, *P* <0.10, Fig. 4b); the *STAT3* gene tended to be down-regulated in the vitamin E group compared to the plant extract and control groups (-54% and -47%, respectively, *P* = 0.09, Fig. 4b). At 28 h after the LPS challenge, fewer changes were observed: *CCL20* and *ADGRE1* tended to be up-regulated in the plant extract group compared to the control group (fold increase of 1.43 and 0.29, respectively, Student’s t-test, *P* <0.10, Fig. 4b). At 76 h after the LPS challenge, only the *IKZF1* gene was up-regulated in the plant extract group compared to the vitamin E group (+14%, Student’s t-test, *P* = 0.02, Fig. 4b).

**Figure 4:**
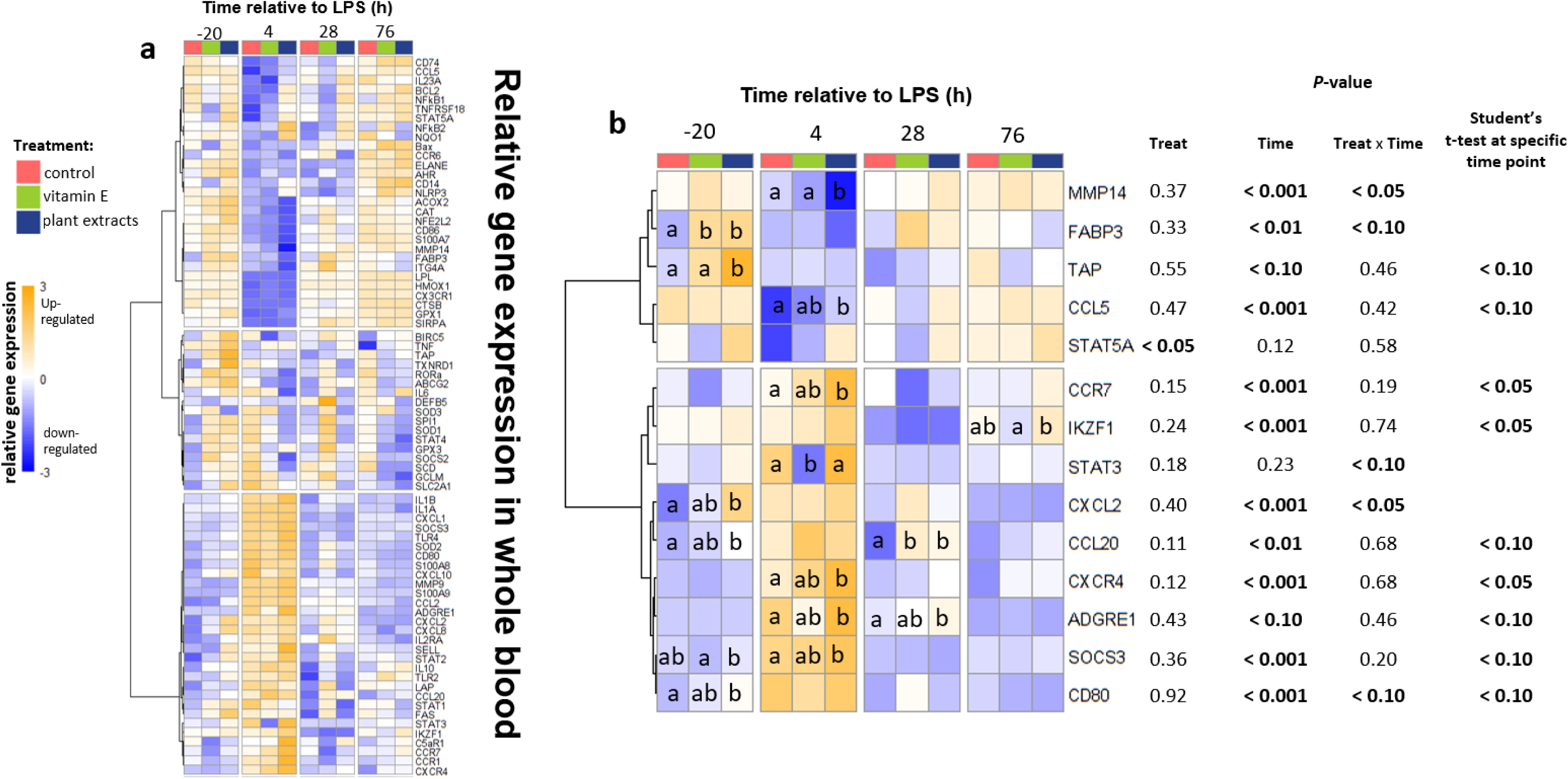
Blood transcriptional profile of genes related to redox and immune status according to the control unsupplemented group (n = 11), vitamin E supplemented group (n = 13), and plant extract supplemented group (n = 12) of dairy cows before (-20 h) and during the LPS challenge (at 4 h, 28 h, 76h). The transcriptional profiles of 79 genes are grouped according to their expression relative to the time of the LPS challenge (**a**), and the expression of inflammatory response genes according to time and treatments (**b**). Migration gene *MMP14* (matrix metallopeptidase 14), growth and proliferation *FABP3* (fatty acid binding protein 3), antimicrobial peptide *TAP* (Tracheal antimicrobial peptide) and *CD80* (CD80 antigen), *CCL5* (C-C motif chemokine ligand 5), *STAT5A* (signal transducer and activator of transcription 5A), *SOCS3* (suppressor of cytokine signalling 3), *CCR4* (C-C motif chemokine receptor 4), *CCR7* (C-C motif chemokine receptor 7), transcription factor *IKZF1* (family zinc finger 1), *STAT3* (signal transducer and activator of transcription 3), *CXCL2* (C-X-C motif chemokine ligand 2), *CCL20* (C-C motif chemokine ligand 20) and *ADGRE1* (adhesion G protein- coupled receptor E1). The control group is represented in red, the vitamin E group in green, and the plant extract group in blue. Significant differences between treatment (Treat) and time (Time) and their interaction (Treat x Time) are noted to the right of the graphs. The data were analysed according to a mixed model and Student’s t-test to differentiate time point differences between treatments. The relative expression of these genes was calculated according to the decimal logarithm of the geometric mean of three reference genes: ACTB (actin beta), YWHAZ (tyrosine 3-monooxygenase/tryptophan 5-monooxygenase activation protein zeta), and RPLP0 (Ribosomal Protein Lateral Stalk Subunit P0). **^a-b^** Mean values in the same row showing a difference regarding the interaction between treatment and time relative to the LPS challenge.

### Mammary epithelium integrity and milk composition during the intramammary challenge and nutritional strategy effects

Milk yield decreased on the day after the LPS infusion but increased thereafter to reach a level similar to that preceding the challenge between day+3 and day+5 (*P* = 0.04, Fig. 5a). The somatic cell counts of milk increased after the LPS challenge and then progressively decreased 28h after the LPS challenge (*P* <0.001, Fig. 5b). The milk protein content was higher before and at 4 h and 7 h after the LPS challenge than at 9 h, 28 h, 52 h and 76 h after the challenge (*P* <0.001, Fig. 5c). It was notably higher in the control group than in the plant extract group at 4 h after the LPS challenge (*P* <0.01, Fig. 5c). The milk lactose content fell immediately after the LPS challenge (lower from 4 h to 52 h after the LPS challenge than before and at 76 h after the challenge, *P* <0.001, Fig. 5d) and tended to be higher in the plant extract group than in the control group after the start of the LPS challenge (48.34 ± 0.73 *vs* 45.95 ± 0.71 g/kg) (treatment x time interaction, *P* <0.06, Fig 5d). The milk Na^+^:K^+^ ratio increased after the LPS challenge but this increase was smaller in supplemented cows than in the control group at 4 h after the LPS challenge (0.82 ± 0.18 and 0.83 ± 0.17 *vs* 1.49 ± 0.18, Student’s t-test *P* = 0.03 and *P* = 0.02, respectively, Fig. 5e). Milk pro-inflammatory cytokine IL8 levels rose with LPS and were higher at 4 h, 7 h, 9 h after the LPS challenge than at the other time points (*P* <0.001). This increase was more pronounced in the plant extract group than in the control and vitamin E groups 4 h after LPS (89 ± 6.14 *vs* 35 ± 6.10 and 51 ± 5.74 x 10^3^ ng/mL, respectively, treatment x time interaction, *P* <0.01, Fig. 5f). Milk cytokine IL1β levels also rose with LPS (higher at 4 h, 7 h and 9 h after the LPS challenge than at any other time points; *P* <0.001, Fig. 5g), whatever the treatments. Other components of milk (minerals) were not affected by either the treatments or the treatment x time interaction (Supplementary data Table S5).

**Figure 5:**
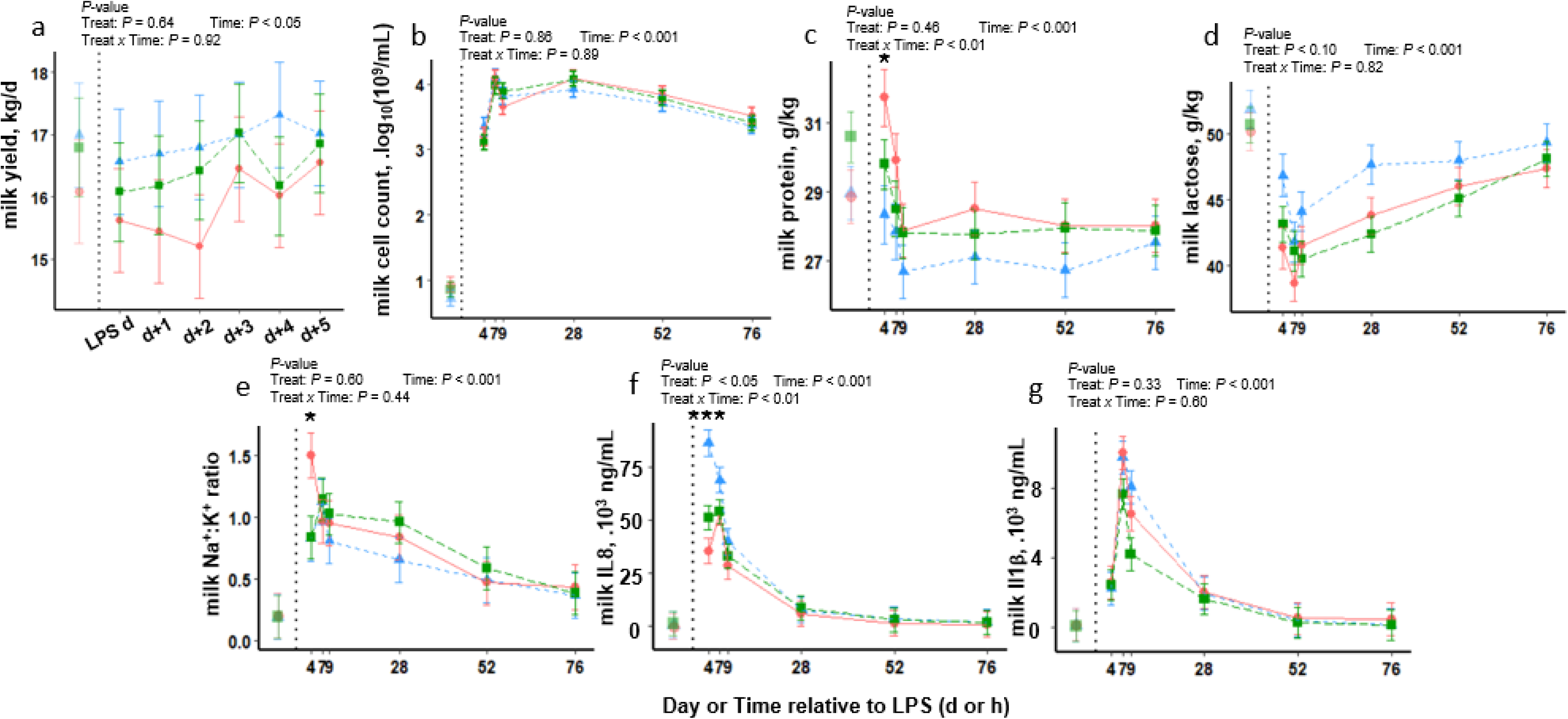
Milk yield and milk composition in the control unsupplemented group (n = 11), vitamin E supplemented group (n = 13), and plant extract supplemented group (n = 12) of dairy cows before and during the LPS challenge (at 4 h, 7 h, 9 h, 28 h, 52 h and 76h). Milk yield in kg/d (**a**), milk cells in log_10_ (10^9^/mL) (**b**), milk protein in g/kg (**c**), milk lactose in g/kg (**d**), milk Na^+^:K^+^ ratio (**e**), milk IL8 in ng/mL (**f**), milk IL1β in ng/mL (**g**). The control group is represented as a solid red line (□●□), the vitamin E group as a dotted green line (--▪--), and the plant extract group as a dotted blue line (- -▴- -). Adjusted means and SEM are represented and the data were analysed according to a mixed model. Significant differences between treatment (Treat) and time (Time) and their interaction (Treat x Time) are noted above the graphs. The thresholds of significance were set at ∗ *P* <0.05 and ∗∗∗ *P* <0.001. For the Mylab laboratory, the analytical limits for the various parameters (linked to our calibration ranges and linearity range) were: Fat: between 20g/L and 56g/L; Protein: between 22g/L and 42g/L; Lactose: between 46g/L and 60g/L; Cells: below 2000 cells/mL. Milk yield was evaluated on 4 quarters with twice-daily milking. Milk cell count, milk protein, milk lactose, milk Na^+^:K^+^, and milk IL8 and IL1b measurements were performed on the LPS-challenged one udder quarter only. The variations between 7 h and 9 h after the LPS challenge and other time points correspond to differentially enriched cisternal or alveolar milk depending on the time of milk accumulation between milkings.

Most of the genes expressed in milk that were under-expressed during inflammation, were linked to milk synthesis (*LALBA, CSN1S1, CSN3, SCD, FAS, LPL, FABP3*), and mammary epithelium integrity (*OCLN, TJP1, CDH1, CLDN1*) (*P* <0.01, Fig. 6a). However, the expression of genes related to cell death (*BAX, BCL2, CASP1, CASP8, CASP13, CTSB, IGFBP5 P* <0.01, *RIPK1*, *P* <0.05) were over-expressed after the LPS challenge (Fig. 6a). By contrast, the expression of genes involved in mammary tissue remodelling (*SPARC, MMP9 P* <0.01) increased in milk after the LPS challenge. The milk transcriptome established from 82 amplified genes indicated three distinct profiles according to treatment and time related to the LPS challenge (Fig. 6a).

**Figure 6:**
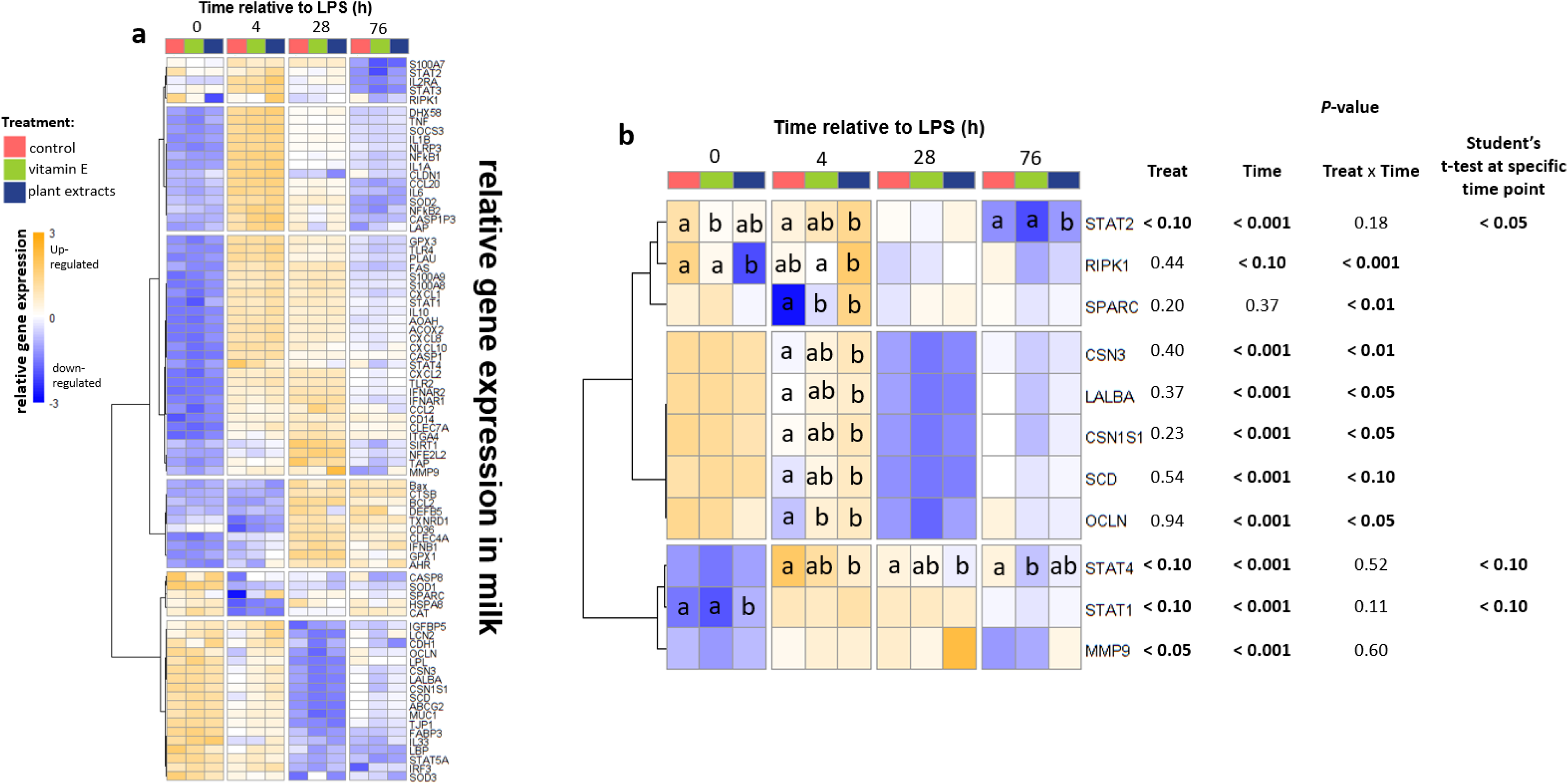
Milk transcriptional profiles of genes related to redox and immune status, milk synthesis according to the control unsupplemented group (n = 11), vitamin E supplemented group (n = 13), and plant extract supplemented group (n = 12) of dairy cows before (0 h) and during the LPS challenge (at 4 h, 28 h, 76h). The transcriptional profiles of 82 genes are grouped according to their expression relative to the time relative of the LPS challenge (**a**), and the expression of inflammatory response and milk synthesis genes according to time and treatments (**b**). Lactose synthesis *LALBA* (lactalbumin α), *CSN1S1* (casein alpha s1), *CSN3* (casein _κ_), cell junction *OCLN* (occluding), necroptosis *RIPK1* (receptor interacting serine/threonine kinase 1), extracellular synthesis *SPARC* (secreted protein acidic and cysteine rich), *MMP9* (matrix metallopeptidase 9), *STAT1* (signal transducer and activator of transcription 1), *STAT2* (signal transducer and activator of transcription 2), *STAT4* (signal transducer and activator of transcription 4) and *SCD* (stearoyl-CoA desaturase). The control group is represented in red, the vitamin E group in green, and the plant extract group in blue. Significant differences between treatment (Treat) and time (Time) and their interaction (Treat x Time) are noted to the right of the graphs. The data were analysed according to a mixed model and Student’s t-test to differentiate time point differences between treatments. The expression of these relative genes was calculated according to the decimal logarithm of the geometric mean of three reference genes: ACTB (actin beta), YWHAZ (tyrosine 3-monooxygenase/tryptophan 5-monooxygenase activation protein zeta), and PPIA (Cyclophilin A). **^a-b^** Mean values in the same row showing a difference regarding the interaction between treatment and times relative to the LPS challenge.

Genes whose expression varied in milk according to treatment or treatment x time interaction are shown in Fig. 6b and Fig. S2 (Supplementary Data). Before the LPS challenge, expression of the *RIPK1* gene (involved in the regulation of cell death) was lower in the plant extract group than in the control and vitamin E groups (-67% and -52%, respectively, Student’s t-test *P* <0.05). At 4 h after the LPS challenge, the expression of this gene was higher in the plant extract group than in the vitamin E group (+ 70%, Student’s t-test *P* <0.001, Fig. 6b), whereas the genes coding for milk proteins (*LALBA, CSN1S1, CSN3*) were upregulated in the plant extract group compared to the control group (fold increases of 0.91, 1.05 and 1.66, respectively, treatment x time interaction *P* <0.01, Fig. 6b). At the same time, *SCD* (involved in the synthesis of the lipids) tended to be more strongly expressed in the plant extract group than in the control group (1.59-fold increase, treatment x time interaction *P* = 0.10), *OCLN* was more strongly expressed in both the plant extract and vitamin E groups than in the control group (fold increases of 1.77 and 1.39, respectively, treatment x time interaction *P* <0.05, Fig. 6b) and *SPARC* (involved in synthesis of the extracellular matrix) was more strongly expressed in the plant extract and vitamin E groups than in the control group (fold increases of 7.31 and 2.35, respectively) (treatment x time interaction, *P* = 0.03, Fig. 6b). *MMP9* was more strongly expressed at 4 h and 28 h after than LPS challenge than before it, and at 76 h after the LPS challenge (*P* <0.001, Fig. 6b) and more in the plant extract group than in the vitamin E group (+ 88%, *P* = 0.05, Fig. 6b), particularly at 28 h and 76 h after the LPS challenge (fold increases of 2.82 and 1.42, respectively, *P* = 0.04 and *P* = 0.06 respectively, Fig. S2). The expression levels of genes related to milk synthesis that were not affected by either the treatment or the treatment x time interaction are shown in Supplementary Data Table S3.

### Local immune status during the intramammary challenge

Some immune genes were over-expressed in milk during inflammation and then decreased progressively, including cytokine genes (*IL1B*, *CXCL8* etc., *P* <0.01), immune stimulation genes (*TAP*, *CD14* etc.) and transcription factors (*STAT1, STAT2, STAT4, NF*_κ_*B1, NF*_κ_*B2 P* <0.01, Fig. 6a). The transcriptomic analysis of milk showed that the expression of inflammatory response genes varied according to supplementation. Before the LPS challenge, the expression level of *STAT1* was higher in the plant extract group than in the control and vitamin E groups (fold increases of 1.95 and 4.07, respectively, Student’s t-test, *P* = 0.06, Fig. 6b). At 4 h after the LPS challenge, the expression of *STAT2* was higher in the plant extract group than in the control group (0.50-fold increase, Student’s t-test, *P* = 0.01). Before and at 76 h after the LPS challenge, *STAT2* gene expression was lower in the vitamin E group than in the control group (+50% and +41%, respectively, Student’s t-test, *P* = 0.05). *STAT4* was lower in the plant extract group at 4 h and 28 h after then LPS challenge than in the control group, and also lower in the vitamin E group at 76 h after the LPS challenge than in the control group (-55%, -30% and -50%, respectively, *P* <0.05). In milk, other expression levels of genes involved in inflammation were measured but no links with supplementation were observed (Supplementary Data Table S3 and Supplementary Data Fig. S2).

The immune capacities of neutrophils measured in milk showed that the percentage of classic MHCII^-^ neutrophils tended to be lower at 4 h after the LPS challenge than at 76 h (*P* = 0.06, Fig. 7a), while the percentage of regulatory MHCII^+^ neutrophils was higher at 4 h than at 76 h after the LPS challenge (*P* <0.001, Fig. 7b). With and without TBHP stimulation, classic MHCII^-^ neutrophils produced more ROS at 4 h after the LPS challenge than at 76 h (*P* <0.01), and the same was seen for regulatory MHCII^+^ neutrophils (*P* <0.05). In the plant extract and vitamin E groups, classic MHCII^-^ neutrophils without TBHP produced and tended to produce more ROS than in the control group at 4 h after the LPS challenge (1.01 ± 0.14 and 0.85 ± 0.12 *vs* 0.71 ± 0.12 MFI, respectively, Student’s t-test, *P* = 0.02 and *P* = 0.06 for plant extract and vitamin E, respectively, Fig. 7c). The nutritional supplements did not affect ROS production by milk neutrophils after TBHP stimulation.

**Figure 7:**
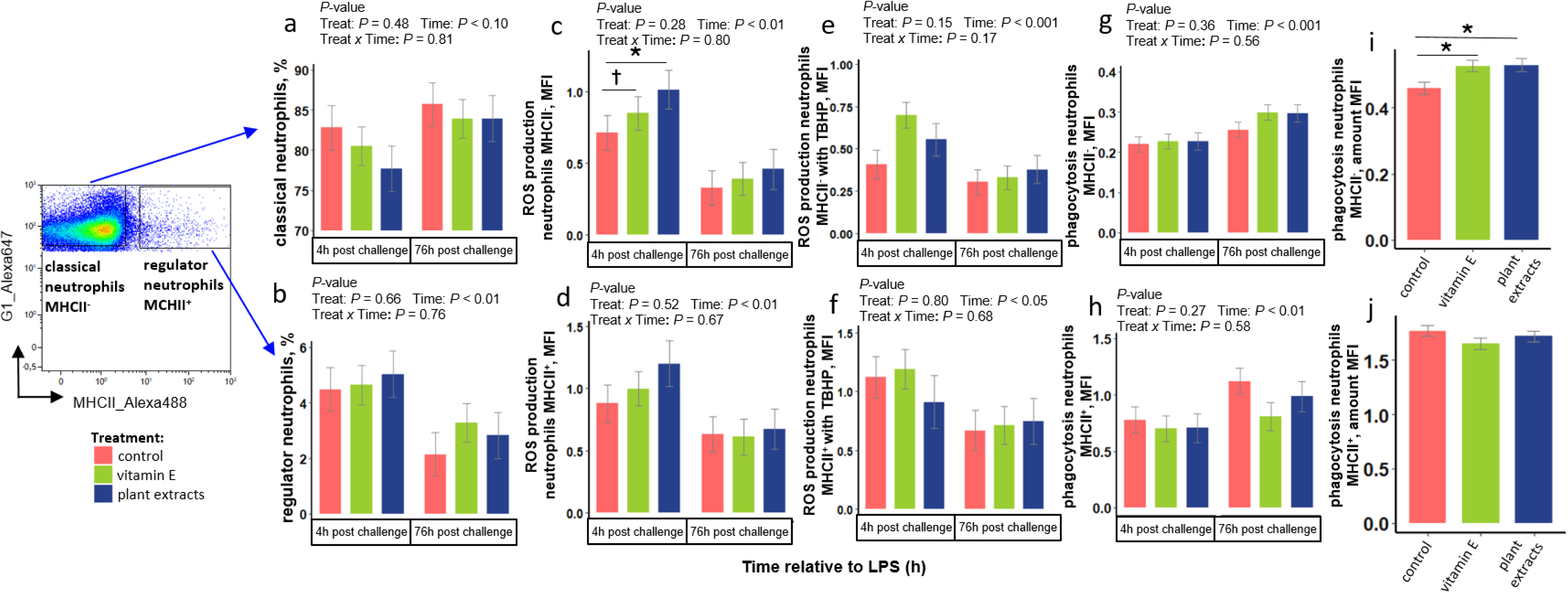
Flow cytometry identification of two types of neutrophils isolated from milk and measurement of their intracellular reactive oxygen species (ROS) production without or with stimulation *ex vivo* and phagocytosis in the control unsupplemented group (n = 11), vitamin E supplemented group (n = 13), and plant extract supplemented group (n = 12) of dairy cows before and during the LPS ch llenge (at 4 h, 28 h and 76h). Percentage of neutrophils without class II major histocompatibility complex, MHCII**^-^** (**a**) and class II major histocompatibility complex neutrophils, MHCII**^+^** (**b**). Mean fluorescence intensity (MFI) of intracellular ROS production after *ex vivo* measurement in the two neutrophil subpopulations by flow cytometry without (**c**, **d**) or with (**e**, **f**) tert-butyl hydroperoxide (TBHP) stimulation. MFI measurement of *ex vivo* phagocytosis with bead *E. coli* by flow cytometry (**g**, **h**). Addition of the phagocytosis activity of the two neutrophil subpopulations from the 4 h and 76 h kinetic points is represented (**i**, **j**).The bar charts shown 4 h after the LPS challenge on the left and 76 h post LPS challenge on the right, each in the same order (control, vitamin E, plant extracts). Thresholds of significance were set at ∗ *P* <0.05 and trends were noted at ^†^ *P* ≤0.10. For the measurement of phagocytic capacity, the time point data were summed to determine whether the treatments had an effect on phagocytic activity during the challenge, as the phagocytic activity of milk was too weak.

Both classic MHCII^-^ and regulatory MHCII^+^ neutrophils in milk phagocytosed at lower rates at 4 h than at 76 h after the LPS challenge (*P* <0.001, Fig. 7g for MHCII^-^ and *P* = 0.01, Fig. 7h for MHCII^+^). Because the phagocytosis signal was too low, we accumulated these signals for both days of cell preparation in order to verify a treatment effect. The amount of phagocytosis during the LPS week showed that classic MHCII^-^ neutrophils in the vitamin E and plant extract groups phagocytosed more than those in the control group (0.53 ± 0.02 and 0.52 ± 0.02 *vs* 0.46 ± 0.02 amount MFI, respectively, *P* <0.05, Fig. 7i). For regulatory MHCII^+^ neutrophils, the amount of phagocytosis did not differ between the treatments (*P* = 0.51). In milk, the macrophages capacities evaluated with their percentages of ROS production and phagocytosis did not display any effects of treatment or time (Supplementary Data Table S6).

## Discussion

Mastitis is an inflammatory disease of the mammary gland in dairy cows, which is most often caused by bacteria. The incidence of mastitis is high at the start of lactation, a period of oxidative stress and inflammation, leading to impaired health. To investigate the effects of vitamin E and plant extracts in preventing mastitis, we triggered an intramammary inflammation with LPS to determine whether these components given as dietary supplements could modulate the ability of cows to initiate an inflammatory response in the mammary gland. The effects of vitamin E or plant extract supplementations were measured on indicators of redox and immune status at the local or systemic levels during the induction of mammary inflammation.

As expected, and as previously observed with the same level of vitamin E supplementation (Corset et al. 2024b), plasma α-tocopherol concentrations were higher in cows supplemented with vitamin E, even during the LPS challenge.

During mammary inflammation, systemic redox status was impaired, with both a reduction in ROM (H_2_O_2_) and antioxidant capacities, as already seen in a previous study (21). However, both the vitamin E and plant extract groups seemed to display better antioxidant capacities in response to the possible oxidative stress occurring during mastitis, with higher glutathione peroxidase activity in erythrocytes and more *SOD3* expressed in the blood with vitamin E and higher FRAP, superoxide dismutase activity in erythrocytes and more *NQO1* expressed in the blood in the plant extract group.

There were no metabolic differences between the supplemented and control groups during the intramammary challenge, which induced higher plasma glucose and lower BHB and calcium in all cows. These metabolic changes during an LPS challenge have already been described in the literature for glucose and BHB (22) and calcium (23). In our study, the LPS infusion induced both local and systemic inflammation. The somatic cell count and levels of cytokines IL8 and IL1β in milk were higher at the first time points after the LPS challenge than before and 28 h after the challenge, thus confirming local inflammation (19, 24). In parallel, the rectal temperature rose and this was followed by the increase in plasma haptoglobin, a classic indicator of systemic inflammation. These results were in line with what could be expected from such an LPS dose being injected in a quarter of the udder (18, 19). The milk composition was modified by the LPS infusion, with more protein and less lactose, as previously observed (21). Unfortunately, we were not able to confirm fat lipolysis during mammary inflammation as in a previous study (21), because the time of the day we sampled milk before the LPS challenge (7 a.m.) was not the same at when we sampled milk during the LPS challenge (11 a.m.). It is known that milk fat content can vary considerably depending on the time of milking (25). We also observed a loss of epithelium integrity during the LPS challenge, with an exchange of minerals between blood and milk leading to an increase in the milk Na^+^:K^+^ ratio. This variation in the Na^+^ K^+^ ratio of milk reflected an opening of tight junctions between mammary epithelial cells, which was confirmed by a reduced expression of genes for tight junction proteins (*OCLN, TJP1, CDH1, CLDN1)* in milk. These observations were in line with a previous study that had shown damage to the mammary epithelium during mastitis (26).

Our results suggest that plant extract supplementation may allow the migration of more white blood cells to the mammary gland during an LPS challenge. IL8 is a pro-inflammatory cytokine that is released by local tissue to enable the migration of immune cells towards the mammary gland during an LPS challenge. A more pronounced rise in IL8 levels in milk in the plant extract group was associated with a fall in the white blood cell count, particularly in blood neutrophils, suggesting a higher migration of blood cells toward the mammary tissue (24). Immune cell migration was likely allowed by the stronger expression of the *CXCR4* cytokine receptor involved in cell migration, cell adhesion and leukocyte extravasation, and the *CCR7* chemokine receptor that controls the migration of memory T-cells to inflamed tissue (27). These two receptors are known to be more strongly expressed in the presence of *E. coli* and are thought to improve the immune response (20, 28, 29). *MMP14* was also less expressed in blood during the LPS challenge, particularly in the plant extract group. A previous study showed that *MMP14* was present in the cell membrane of monocytes for the transendothelial migration of monocytes and T-cell attraction, and that it was reduced during a LPS challenge in mice (30, 31). Thus the decrease in the expression of *MMP14* in the blood during our study could be an indication of cellmigration and therefore of the fall of white cell counts. Although the migration of white blood cells to the mammary gland appeared to be higher among cows in the plant extract group, it did not increase the milk somatic cell count. One reason might be improved mammary epithelium integrity in the plant extract group where the stronger expression of *IKZF1* suggested B-cell maturation and perhaps a modulation of the adaptive immune response (32). However, as far as we know, *IKZF1* has not yet been studied in the context of mastitis in dairy cows. It is also interesting to note that the immune system was probably better regulated in the plant extract group because of the stronger expression of *STAT5A*, a gene that induces pro-cytokine production in serum and has been qualified as a potentially effective marker of resistance to mastitis (33).

At the level of the local immune response, inflammatory capacities seemed to be greater in both the vitamin E and plant extract groups, with more ROS produced (oxidative burst) and more phagocytosis from classic neutrophils in milk, as observed in previous studies (4, 11). However, the ROS production of milk macrophages did not vary, thus confirming the findings of another study involving vitamin E supplementation (34). Local neutrophil capacities were probably increased in order to eliminate and detoxify LPS from the mammary tissue. At the local level in milk, cows in the plant extract group also displayed a stronger expression of *STAT2*, which has been shown to be inhibited by intracellular oxidative stress (35) which can lead to immunosuppression (36). This suggests that plant extracts may help to prevent inhibition of the immune response by enhancing the ability of neutrophils to manage high ROS production during mammary inflammation. At the local level also, *STAT4* expression was reduced by both plant extracts and vitamin E at the start and end of inflammation. *STAT4* allows the expression of IL12 for the maturation of T lymphocyte-helper 1 into cytotoxic T-lymphocytes during *E. coli* mastitis (37). We can therefore suppose that the supplementations with vitamin E and plant extracts may have decreased the inflammatory response of cytotoxic T-lymphocytes.

At the systemic level, in classic and regulatory neutrophils, vitamin E enabled a reduction in the oxidative burst during the LPS challenge but did not influence phagocytosis. This was consistent with the findings of a previous study in which vitamin E also reduced ROS production in both types of neutrophils(7). In the vitamin E group, *CCL20* was more strongly expressed in blood, which may have induced the recruitment of neutrophils, as seen in mammary epithelial cells stimulated by *E. coli* (20, 38). Other genes that tended to be upregulated in the blood may also have been involved in the immune cell migration observed in the plant extract group, such as that of the chemokine *CCL5*, *STAT3* and *ADGRE1*. *CCL5* and *STAT3* can recruit leukocytes as they have been linked to improvements in the immune response (20). *ADGRE1* is involved in adhesion between immune cells and feedback regulation by increasing *SOCS3* expression (38). It has been shown in several species that these three genes are expressed in monocytes/macrophages during an LPS challenge, and they are candidate genes for the regulation of immunity (38). In our study, cows in the vitamin E group tended to have a weaker expression of *STAT3* and *ADGRE1,* which may have been related to the non-migration of immune cells towards the mammary gland, this being consistent with the increase in white blood cells observed during a LPS challenge in another study (38).

Interestingly, at the onset of inflammation in our study, vitamin E and plant extracts seemed to enable better integrity of the mammary epithelium, with more *OCLN* expression in milk, a lower milk Na^+^:K^+^ ratio and more *SPARC* expression for extracellular matrix synthesis. At the same time, the plant extracts probably also improved the remodelling of mammary tissue during the LPS challenge, with an increased expression of the *RIPK1* gene that is associated with cell death and cytokine signalling (39). In conjunction with the expression of *RIPK1* and *SPARC*, the increase in *MMP9* expression in milk during the LPS challenge in the plant extract group may also have contributed to greater mammary tissue remodelling. The *MMP9* gene encodes for an enzyme responsible for mammary tissue remodelling during mammary involution (40), and is known to be more elevated at the end of inflammation (38). Improved mammary epithelium integrity associated with better mammary tissue remodelling under plant extract supplementation may also have been the reason for a lower milk protein content, suggesting less transfer of proteins from blood to milk. This greater integrity of the mammary epithelium was associated with higher blood cortisol levels, which could be indicative of a better adaptation to stress (as suggested by previous studies,41, 42), protecting mammary epithelium integrity and fighting the effects of the LPS challenge. Also, in the plant extract group, the higher milk lactose content during the LPS challenge may have been related to the reduced loss of lactose through a less disrupted epithelium and higher *LALBA* expression at 4 h after LPS. The effects of plant extract supplementation on milk composition were thus also associated with an upregulation of the genes involved in the synthesis of milk constituents (for *LALBA*, *CSN3*, *CSN1S1*) and a tendency towards more *SCD* being expressed in milk.

Ultimately, immunity was perhaps better regulated by the antioxidant capacities of the vitamin E and plant extracts, because immune cells are sensitive to oxidative stress (36). These beneficial local effects of vitamin E and plant extracts during the challenge may possibly have been due to their antioxidant function, as previously reported (43). Dairy cows supplemented with plant extracts and vitamin E may have a better immune response to an LPS challenge, thanks to, or inducing, better mammary epithelium integrity in both groups.

It should be noted that some nutritional supplement effects occurred before the LPS challenge, but these effects remained minor. At the systemic level, before the challenge, both supplements (vitamin E and plant extracts) tended to reduce *FABP3* expression in the blood, and vitamin E tended to reduce *STAT2* expression in milk. Since *STAT2* is known to be activated by type I IFN and to induce cytokine synthesis (44), it is possible that the tendency in the vitamin E group towards a reduction in *STAT2* expression may have inhibited cytokine production and possibly avoided hyper-inflammation, as previously observed in healthy cows supplemented with the same diets (Corset et al. 2024b). In line with these observations, a negative association has been described between genes related to lipid metabolism and those related to the immune system. (45). We also observed that *FABP3* expression tended to be lower in the supplemented groups. Before the LPS challenge, it seems that plant extracts induced an upregulation of some genes when compared to vitamin E. For instance, we observed a specific effect of plant extracts to increase *CXCL2* and a tendency to increase *CCL20*, *CD80* and *TAP* expressed in blood, which may suggest a potential for plant extracts to activate immune gene expression. *CXCL2* is important to controlling the early stage of neutrophil recruitment during tissue inflammation (46).

## Conclusion

We have demonstrated for the first time that nutritional strategies can modulate the immune responses of dairy cows during an LPS challenge. Regarding the systemic immune response, plant extracts increased the expression of chemoattractant and adhesion genes at the start of the inflammatory process for cell migration towards the mammary gland, while vitamin E limited the release of ROS by classic neutrophils. Both nutritional strategies helped to regulate the systemic immune response. This regulation probably enabled improved local immune capacities (oxidative burst and phagocytosis) in supplemented cows, possibly contributing to better integrity and remodelling of the mammary epithelium. Plant extracts may be more beneficial, with a tendency towards a higher milk lactose content and the stronger expression of genes involved in milk synthesis. These positive local effects during the challenge with vitamin E and plant extracts may have been due to their antioxidant function (as described in the literature) and might enable the prevention of mastitis. It is now necessary to further investigate whether these supplements act directly or indirectly on the genes we found to be affected in our study. Finally, it is still not clear whether supplementation induces better immunity, leading in turn to improved mammary tissue integrity and remodelling, or whether the opposite is true.

## Methods

### Animals, experimental design and animal housing

This experiment was conducted at the INRAE Experimental Farm (IE PL, INRAE, Dairy Nutrition and Physiology; https://doi.org/10.15454/yk9q-pf68, 35650 Le Rheu, Brittany, France accreditation for animal housing no. C-35–275-23”) between 8 September and 11 December 2022. The experiment was approved by the Rennes Ethics Committee on Animal Experimentation and the French Ministry for Higher Education, Research and Innovation (APAFIS project number # 37759-2022062117354678_v2) and performed in compliance with all applicable provisions established by European Directive 2010/63/UE.

Forty-five lactating Holstein dairy cows, clinically healthy and non-pregnant, were assigned to one of three groups. Among them, 36 were followed from 5 weeks before to 5 weeks after calving, with an average of 12 cows per group. These 36 cows were selected before their fifth week of lactation on the basis of their health status as specified in the Materials and Methods section related to the LPS challenge.

All cows were fed a total mixed ration (TMR) based on supplemented maize silage and formulated according to INRAE guidelines (Supplementary Table S7) (47). Three nutritional strategies were randomly assigned to each group. Due to discrepancies between the actual and predicted calving dates, the final number of cows per group differed slightly from the initial target of 12 per group. The “control” group received the standard TMR only and included 11 cows for the LPS challenge. The “vitamin E” (*all-rac-*α-tocopheryl acetate) group received the standard TMR supplemented with vitamin E and included 13 cows for the LPS challenge. The “plant extract” group received the standard TMR supplemented with plant extracts from Biodevas Laboratoires (Savigné-L’Evêque, France) and included 12 cows for the LPS challenge. In order to ensure sufficient statistical power for our design, we conducted a review of published studies that had examined the effects of oral vitamin E supplementation (7) or intramammary lipopolysaccharide (LPS) injections (18, 48) on gene expression or the immune, production, and metabolic responses of dairy cows. A total of between 9 to 12 cows per group was sufficient to observe significant effects on these responses in the three studies. The cows were allocated in a way that minimized any between-group variability that might have been linked to their pre-experimental characteristics. For multiparous cows, data on their milk yield during the first 60 days of the previous lactation, the average milk somatic cell count (SCC) during the last month of the previous lactation, and the last body condition score (BCS) before the start of the experiment were balanced between the groups. For primiparous cows, only the average of the last two body weight measurements before the experiment started was balanced. The control group comprised six multiparous and five primiparous cows, the vitamin E group eight multiparous and five primiparous cows, and the plant extract group eight multiparous and four primiparous cows.

The cows were housed in a free stall with free access to feed and water but a controlled individual intake, except for the 1 to 2 days around calving when they were placed in individual stalls. For the vitamin E group, vitamin E was added to the diet (Cooperl, Plestan, France) at a rate of 30,000 mg/kg (1 mg/eq IU). The cows received 3000 IU/day (100 g/day) for four weeks before calving and 1000 IU/day (35 g/day) for five weeks of lactation after calving (6, 49, 50). The plant extracts were designed using the following plants: *Sambucus nigra*, *Salix alba*, *Laurus nobilis, Haragophytum procumbens*, *Silybum marianum* and *Arctium lappa* (Biodevas Laboratoires, Savigné-L’Evêque, France). The plant extract preparation had received AFNOR certification, attesting to their compliance with ISO 22000 standards and thus ensuring their safety. The plant extracts had also been certified under the Feed Certification Scheme (GMP+ International) to guarantee worldwide food safety. The plant extracts were administered at a rate of 10 g/day from four weeks before calving until five weeks after calving. Both vitamin E and plant extracts were added in solid form top dressed on the TMR.. The amounts of feed offered and feed refused were measured daily in order to calculate the dry matter intake (DMI) (Supplementary Data Table S7). The cows were milked twice daily at 07:00 am and 4:30 p.m. prior to feeding. Milk yield was recorded individually at each milking. Health events were monitored daily and cows with health problems at calving (stillbirth, abortion) or lameness problems requiring isolation from the herd were excluded. At the same time points, rectal temperatures were measured with a digital thermometer to monitor inflammation and to be alerted if the temperature did not decrease, and an observation form was completed to monitor pain thresholds.

### Intramammary lipopolysaccharide challenge on one udder quarter

An intramammary LPS challenge was performed 5 weeks after calving. After the morning milking of all the cows involved in the 5^th^ week of lactation, one healthy rear udder quarter was infused with 10 µg Vaccigrade ultrapure LPS from *Escherichia coli* 0111:B4 (Invivogen, Toulouse, France) resuspended in 2 mL sterile PBS (CDM Lavoisier, Paris, France) containing 0.5% BSA (cell culture grade, endotoxin free, A9576, Sigma-Aldrich, Saint-Quentin-Fallavier, France). The LPS solution was prepared extemporaneously in a sterile environment and injected after aseptic preparation of the teat and orifice with 70% alcohol. The injection was performed into the teat cistern via the teat canal, using a sterile disposable syringe fitted with a teat cannula. The udder was then briefly massaged. The selection of cows and udder quarters to be challenged was based on health status according to the health records. A sample of milk was collected from each of the four quarters to check for the absence of mammary infection, which was assessed by SCC <20,000/mL during the week before the challenge and the day before the challenge. Bacteriological analysis of the milk was performed on the day before the challenge and a second time during the challenge by spreading a drop of quarter milk on a Petri dish of blood agar, incubating for 24 hours at 37°C and validating by the absence of bacterial colonies.

### Milk sampling

During the week of inflammation, 40 mL cisternal milk samples were collected from the infused teat at 0 h (just before the LPS infusion) and then 4, 7, 9, 28, 52 and 76 h after.. At some time points (0, 9), milk samples were collected before milking at 7:00 am and 4:00 pm, while at others, (4, 28, 52 and 76 h) the milk samples were collected 4h after milking at 11:00 a.m. For the 7h time point, milk was collected at 2:00 p.m. All sample collections were performed according to microbiological procedures (51). Briefly, the teats were carefully washed and dried, and then several streams of milk from the foremilk contained in the teat were discarded. Teat disinfectant was sprayed on the teats before milking and dried thoroughly with a single towel. The inflammatory quarter teat was rubbed with a moist gauze pad containing 70% alcohol and then 40 mL milk was collected for analysis.

The milk samples were analysed for milk protein, fat, lactose, cytokines and SCC (cell count in logarithm decimal) using an infrared method (Mylab, Châteaugiron, France). Integrity of the mammary epithelium was estimated by measuring the milk sodium and potassium Na^+^:K^+^ ratio. Milk minerals were measured in whole milk using the ICP-OES method (5110 Agilent Technology, Les Ulis, France) as previously reported (57). Plasma mineral concentrations were determined by ICP-OES (5110 Agilent Technology) after dilution with nitric acid (2% v/v) and ultrapure water, with the addition of Triton X-100 (0.01% v/v).

Quantitative real-time PCR analysis was employed to quantify RNA in milk samples collected at 0, 4, 28 and 76 h after the LPS challenge. The functional capacities of immune cells were evaluated in milk samples collected 4 and 76 h after the LPS challenge.

### Blood sampling

Blood samples were collected from the jugular vein at 20 h before the LPS infusion, and then at 4, 18, and 76 h post-infusion. Blood was collected in three tubes containing lithium heparin as an anticoagulant (Vacutest, Kima srl, Arzergrande, Italy) for the analysis of plasma metabolites, the redox balance and inflammatory markers, as well as calcium, as described in the Biochemical Analyses section, below. In addition, blood was collected in two tubes containing K_2_ EDTA as an anticoagulant (Vacutest) for the analysis of plasma vitamin E. Plasma was separated by centrifugation at 1,200 *g* for 10 min at 4°C and stored at −20°C, and stored at -80°C for RNA analysis samples. Plasma mineral levels were determined by ICP-OES (5110 Agilent Technology) after dilution with nitric acid (2% v/v) and ultrapure water, with the addition of Triton X-100 (0.01% v/v).

Blood was also collected in one lithium heparin tube to determine the haematological profile, antioxidant enzyme activity in plasma and erythrocytes and the functional capacities of blood immune cell function.

### Quantification of IL8 and IL1**β** milk cytokines

The quantification of IL8 in milk was measured using goat anti-bovine interleukin-8 Ab AHP2817, recombinant bovine interleukin-8 PBP039 and goat anti-bovine interleukin-8 Ab conjugated with biotin AHP2817B, using commercially available ELISA kits (BioRad, Hercules, CA), as in a previous study (52). IL1β in milk was quantified with rabbit anti-bovine interleukin-1β Ab AHP851B, recombinant bovine interleukin-1β PBP008 and mousse anti-sheep interleukin-1β Ab conjugated with biotin MCA1658, using commercially available ELISA kits (BioRad).

### Isolation of neutrophils and monocytes/macrophages in milk and blood

Cells prepared from 20 mL milk subsamples were resuspended in PBS (10 mM phosphate, 150 mM sodium chloride, pH 7.3 to 7.5) containing 50% (v/v). The preparation was gently stirred, allowed to stand for 15 min and then centrifuged (10 min, 1400 g). The cream was removed with a spatula and the supernatant with a pipette, leaving approximately 300 µL above the pellet; 500 µL PBS solution (2 mM EDTA, 1% horse serum) were added to resuspend the pellet. To filter the suspended pellet, 500 µL of the previous PBS solution was previously soaked in a 15 mL tube, and the sample was passed through a 200 µm filter. The cells were washed and suspended in 9 mL of the previous PBS solution. At this step, the number of cells recovered was counted with the TC20 Automated Cell Counter (Biorad) to verify that approximately 10 × 10^6^ cells were present in each pellet. The cell samples were secondly passed through a 70 µm filter (Miltenyi Biotec, Paris, France) and centrifuged at 700 g for 5 minutes at room temperature. The pellet was suspended in 700 µL of the previous PBS solution and plated at 2 × 10^6^ cells/well in a 96-well plate.

Cells prepared from the blood samples were purified after centrifugation at 1,000 g for 15 min at 20°C to obtain the red pellet fraction containing neutrophils and the buffy coat fraction containing monocytes. Each cell fraction was lysed by adding four volumes of ACK buffer (ammonium-chloride-potassium, Gibco) to one volume of blood. The neutrophils were then washed twice with PBS, counted with a TC20 automated cell counter, suspended in PBS containing 10% horse serum and 2 mM EDTA at a concentration of 10^7^ cells/mL, and plated at 2 x 10^6^ cells/well in a 96-well plate, as in a previous study (53).

In blood and milk, monocytes/macrophages, classic neutrophils and regulatory neutrophils were identified using the same protocol with fluorochrome-linked antibodies. Neutrophils were identified by flow cytometry, as previously described (53). Neutrophils were labelled with an anti-G1 IgM primary antibody (CH138A, 1/500 (v/v), Kingfisher, MN, USA) and then with an anti-IgM secondary antibody coupled to fluorochrome 647 (goat anti-mouse IgM Alexa Fluor 647, 1/200 (v/v) Invitrogen, MN, USA). Regulatory neutrophils were labelled with an anti-MHCII IgG1 primary antibody (CAT82A IgG1, 1/500 (v/v), Kingfisher) and then with an anti-IgG1 secondary antibody coupled to Alexa Fluor 488 (goat anti-mouse IgG1 Alexa Fluor 488, 1/200 (v/v) Invitrogen). Monocytes/macrophages were labelled with an anti-CD14 IgG2a monoclonal antibody coupled to RPE-Alexa Fluor 750 (mouse anti-human CD14:RPE-Alexa Fluor 750, 1/20 (v/v), Bio-Rad Laboratories Inc, Hercules CA, USA) and MHCII as previously for neutrophils. Isotype controls of mouse IgG1 and IgM (Invitrogen) were used to discriminate non-specific background during analysis (54).

For the identification of blood cells, monocytes and neutrophils, the wells were duplicated for ROS production and phagocytosis tests under stimulated or unstimulated conditions. For milk, the macrophages and neutrophils were found in the same well and only the ROS production assay was duplicated under stimulated or unstimulated conditions; the phagocytosis assay was performed under stimulated conditions only.

### Measurement of ROS production indicative of the functional capacities of neutrophils and monocytes/macrophages in blood and milk

ROS production by monocytes/macrophages and neutrophils was quantified using CellROX Orange Flow Cytometry Assay Kits (CD10493, Invitrogen) according to the manufacturer’s instructions. Briefly, to induce ROS production, 500,000 cells/well for blood cells and 700,000 cells for milk cells were analysed according to the method described in a previous study (53). A total of 250,000 events were acquired using a MACSQuant Analyzer 10 cytometer (Miltenyi Biotec), and after compensation, the results were analysed with Kaluza software 114 (analysis version 2.1, 2009-2021 Beckman Coulter, Inc.) (54) using a gating strategy (Supplementary Figs. S3 and S4). Cell numbers were expressed as a percentage and ROS production in mean fluorescence intensity for statistical analysis.

### Measurement of phagocytosis indicative of the functional capacities of neutrophils and monocytes/macrophages in blood and milk

Phagocytosis was measured in monocytes/macrophages and neutrophils using pHrodo *Escherichia coli* BioParticles conjugate for phagocytosis (Invitrogen) according to the manufacturer’s instructions and the method described in a previous study (55). Briefly, 500,000 cells/well for blood cells and 700,000 cells/well for milk cells were incubated in a 96-well black microplate for 45 min at 37°C with 20 µg/well pHrodo *E. coli* BioParticles. Because of time constraints, these phagocytosis tests were run through the cytometer the next day. Cell preservation was maintained at 4°C for a maximum of 3 days with 50 µL per well 4% paraformaldehyde (32%, Paraformaldehyde Electron Microscopy Sciences, Hatfield, Pennsylvania, United States) diluted in PBS (v/v). A total of 350,000 events linked to ingestion of the *E. coli* BioParticles were acquired using a MACSQuant Analyzer 10 cytometer (Miltenyi Biotec), and after compensation, the results were analysed with Kaluza software 114 (analysis version 2.1, 2009-2021 Beckman Coulter, Inc.) using a gating strategy (Supplementary Figs. S3 and S4). Cell percentages for cell numbers and mean fluorescence intensity for phagocytosis (only wells with beads enabled evaluation of the effects under study) were considered for statistical analysis.

### RNA extraction and gene expression analysis of blood and milk

A subsample of 200 µl milk was freshly transferred to a 2 mL Lysing Matrix D tube (Thermo Fisher Scientific, Strasbourg) and 800 µl Trizol (Invitrogen Life Technologies, Carlsbad, CA) was added. The samples were stored at -80°C until extraction. The milk RNA sample was crushed and homogenized for two cycles of 150 s at 6000 rpm for RNA extraction using a Precellys ball mill (DQ2368, Bertin, Montigny-le-Bretonneux, France) and the tube was frozen at -80°C until extraction. A subsample of 200 µl blood was freshly diluted in 200 µl DL lysis buffer (Macherey-Nagel, Düren, Germany) and the tube frozen at -80°C until extraction. Total RNA was extracted from cell-sorter purified cells using the NucleoSpin RNA Milk kit with a DNase treatment (Macherey Nagel) and reverse transcribed using iScript Reverse Transcriptase Mix (Bio-Rad) according to the manufacturer’s instructions.

The primers used for PCR amplifications of the milk and blood samples (Eurogentec, Seraing, Belgium) are listed in Supplementary Table S8. Primer validation was performed on a serially diluted pool of complementary DNA using a LightCycler 480 Real-Time PCR system (Roche, Boulogne-Billancourt, France). Pre-amplification was performed first with steps of 2 min at 95°C, 20 cycles consisting of 15s - 95°C and 4 min -60°C to 4°C to infinity, for 1.25 µL complementary DNA and 3.75 µL amplification solution. The pre-amplification solution for 96 genes consisted of a mixture containing all forward and reverse primers (v/v, 13:100). For the 96 genes to be amplified, we used 1 µL forward primer and 1 µL reverse primer of 10 µM stock solution. The volume of this mixture was adjusted to 200 µL with an RNase-DNase-free buffer (10 mM Tris, 8. 0 pH, 0.1 mM EDTA, Teknova, CA, USA), PreAmp Master Mix solution (v/v, 27:100) (Fluidigm 100-5581), and ultrapure water (v/v, 6:100). This pre-amplification was followed by exonuclease treatment for 30 min - 37°C, then 15 min - 80°C and then 4°C (Exonuclease I, New England Biolabs). After exonuclease treatment, the samples were diluted 5× with buffer (10 mM Tris-HCl, 1.0 mM EDTA, Teknova) and stored at -20°C until Fluidigm quantification, which was performed using the Biomark HD (Fluidigm) in a 48 x 48 well plate according to the manufacturer’s instructions. The melting temperature was 60°C. Data were analysed using Fluidigm Real-Time PCR software to determine the cycle threshold (Ct) values (54). Only genes expressed with a correct melting temperature and in 40% of the samples were retained in the analysis. So that the data could be analysed when the gene was very weakly expressed and undetectable by the PCR method (e.g. some genes before the LPS challenge), we attributed a value of Ctmax+1 when the data was missing for some samples. Ct values from milk samples were then expressed relative to the geometric mean of three reference genes (*ACTB*, *PPIA* and *YWHAZ*), and CT values from blood samples according to the geometric mean of three reference genes (*ACTB*, *YWHAZ*, *RPLP0*), using a semi-absolute method as previously reported (56). The transcriptomic profiles of blood and milk samples from all dairy cows were established with 79 and 82 genes, respectively.

### Blood haematological profiles

Haematological profiles were analysed by the Labocea Laboratory (Fougères, France) to determine the white blood cell count (10^9^/L) with a haematology counter and the percentage of monocytes and polynuclear neutrophils by microscopic readings after May-Grünwald Giemsa staining. The percentages of each cell type were multiplied by the total white blood cell concentration to obtain the count.

### Antioxidant enzyme activity of plasma and erythrocytes

The activities of GPx (glutathione peroxidase) and SOD (superoxide dismutase) enzymes were measured in plasma and erythrocytes, which were isolated and then stored at 80°C until analysis (58). Measurements of GPx activity in plasma and erythrocyte lysates were based on the amount of NADPH formed by an enzymatic reaction. A solution of 20 mM GSH (glutathione) and 2 mM NADPH formed a GSH-NADPH2 complex; the addition of 3 mM H_2_O_2_ then released NADPH at pH 7.0, measured at 37°C by absorbance at 340 nm after 300 seconds. These activities were measured on a multiparameter analyzer (Kone Instrument Corp., Espoo, Finland). This protocol was adapted from a GSH-Px assay protocol used in pigs (59).

### Determination of plasma metabolites, haptoglobin and redox status

Plasma glucose, urea, NEFA and BHB for energy metabolism, ROM for oxidative stress, BAP and FRAP for total antioxidant capacity and haptoglobin for inflammation were measured using a multiparameter analyzer (Kone Instrument Corp.) and appropriate kits as previously described (53). Tocopherols were quantified from plasma as previously described (60), in an Acquity UPLC system (Waters, Saint-Quentin-en-Yvelines, France) using a 150 × 2.1 mm, HSS T3, 1.8 µm column for separation, followed by detection with a photodiode array detector at 295 nm.

### Statistical analysis

All data were analysed using the lmerTest package for a linear mixed effects model procedure on R Studio (version 1.3.1093, 2009-2020) with the following model:

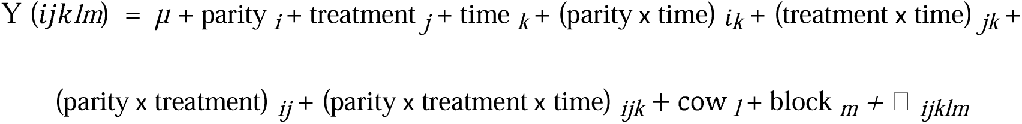

Y _*ijklm*_ was the variable analysed. Parity *_i_* was the fixed effect of the parity, i.e. primiparous or multiparous (1 degree of freedom). Treatment *_j_* was the fixed effect of the group, i.e. control, vitamin E or plant extracts (2 degrees of freedom). Time *_k_* was the fixed effect of time corresponding to time points before or after the LPS challenge, where the number of degrees of freedom was sampling dependent. Cow *_l_* was the cows considered as a random effect, block *_m_* was a fixed effect of the blocks (3 degrees of freedom) and □ ijklm was the residual error. Means and SEM (standard error of the mean) were used to plot the graph. Post-hoc two-way comparisons between modalities of parity, weeks or interactions were also calculated according to Satterthwaite’s method. Significance thresholds were set at ∗ *P* <0.05; ∗∗ *P* <0.01, and ∗∗∗ *P* <0.001, and trends were noted at ^†^ *P* ≤0.10. Student’s t-tests were used regarding time relative to LPS when the mixed model tests for the treatment or the treatment x time effect were not significant (*P* >0.10). Figures were plotted in R Studio using ggplot2 package (version 3.3.5, 2021), and based on the means and SEM resulting from the double interactions, specifically treatment by time before and after the LPS challenge. The transcriptomic profiles of blood and milk were plotted on Rstudio using the pheatmap package (version 1.0.12, 2018).

## List of abbreviations

IU: International unit
LPS: lipopolysaccharide
ROS: Reactive oxygen species
MDA: malondialdehyde
BAP: Biological Antioxidant Potential
FRAP: Ferric reducing antioxidant power
ROM: Reactive oxygen metabolite
DMI: Dry matter intake
NEFA: Non esterified fatty acid
BHB: β-hydroxybutyrate
GPx: Glutathione peroxidase
SOD: Superoxide dismutase
MHC: Major histocompatibility complex
TBHP: Hydroperoxyde de tert-butyle
MFI: Mean of fluorescence intensity
IL8: Interleukin-8
IL1b: Interleukin-1β
IFN: interferon
TMR: Total mixed ration
SCC: Somatic cell count
BCS: Body score condition
SEM: Standard error of the mean.

## Supplementary information

The online version contains supplementary material available at doi (link coming soon).

## Acknowledgements

Angelique Corset’s research received financial support from the Biodevas Laboratoires and Association Nationale de Recherche Technologique (ANRT) to cover her salary. Financial support for the animal study was provided by the Pays de Loire and Brittany regions, and BPI France in the context of the NEOLAC project. The authors would like to thank Dr Gaetan Vetea Plichart and all the technicians and animal staff at the IE PL experimental farm (le Rheu, France): Gaël Boullet, Françoise Pichot, laboratory technicians in the PEGASE Joint Research Unit (Saint-Gilles, France) Frédérique Mayeur, Perrine Poton, Jacques Portanguen for their assistance during the planning and execution of the experiment, Noémie Gaultier for blood and milk analyses and Maryline Lemarchand for diet analysis. Thanks also to the technicians, and particularly Christophe Gitton in the Bacterial Infections and Immunity of Ruminants (IBIR) team at the INRAE ISP unit (Nouzilly, France) and Agathe Degura for plasma vitamin E analysis at INRAE-VetagroSup Herbivores unit (Saint-Genès-Champanelle, France).

## Authors’ contributions

Drs A.C., A.B., M.B., P.G., B.G. and A.R. designed the protocol and all experiments in the dairy cows, performed the experiments, participated in the statistical analysis and interpretation of the results, and wrote the manuscript. A.C. performed the experiments in dairy cows, performed certain laboratory analyses, created the database, performed statistical tests, interpreted and represented the results and wrote the manuscript. Dr J-FR. provided funding, supported the design and implementation of the study, and reviewed the manuscript. O.D. participated in overall monitoring of the experimental farm and the collection of zootechnical data. K.R-S and S.P. performed and designed certain laboratory analyses by adapting the protocols for dairy cows. All authors read and approved the final manuscript.

## Availability of data and materials

Additional file 1: Table S1 at S8. The file contains additional results including **Table S1** entitled ‘Plasma energy metabolism indicators, vitamins, and minerals in the control unsupplemented group (n = 11), vitamin E supplemented group (n = 13), and plant extract supplemented group (n = 12) of dairy cows before supplementation and during the LPS challenge etc.’, **Table S2** entitled ‘Abundance of antioxidant function mRNA, immune function mRNA and gene cell death mRNA determined by real-time quantitative RT-PCR in blood etc.’, **Table S3** entitled ‘Abundance of antioxidant function mRNA, immune function mRNA, milk synthesis mRNA and gene cell death mRNA determined by real-time quantitative RT-PCR in milk etc.’, **Table S4** entitled ‘Plasma haptoglobin and cortisol, haematological profiles and functional capacities of immune cells etc.’, Table S5 entitled ‘Measurement of rectal temperature, milk content and biomarkers of mammary epithelium integrity etc.’, and **Table S6** entitled ‘Functional capacities of milk neutrophil and macrophage immune cells etc.’. This file also contains additional materials and methods including **Table S7** entitled ‘Composition of dry matter ingredients and chemical composition of the diet during close-up, first 15 DIM (days in milk) and after 15 DIM in dairy cows’, and **Table S8** entitled ‘Forward and reverse primers used for real-time quantitative PCR in milk and blood samples to measure the gene expression of antioxidant, immune or milk production functions’.

Additional file 2: Figure S1 at S4. The file contains additional results including, **Fig. S1** entitled ‘Abundance of blood mRNA determined by real-time quantitative PCR in the control unsupplemented group (n = 11), vitamin E supplemented group (n = 13), and plant extract supplemented group (n = 12) of dairy cows before and during the LPS challenge (at 4 h, 28 h, 76h) etc.’ and **Fig. S2** entitled ‘Abundance of milk mRNA determined by real-time quantitative PCR in the control unsupplemented group (n = 11), vitamin E supplemented group (n = 13), and plant extract supplemented group (n = 12) of dairy cows before and during the LPS challenge (at 4 h, 28 h, 76h) etc.’ This file also contains additional materials and methods including **Fig. S3** entitled ‘Flow cytometry gating strategy for the identification of monocytes (MCHII^+^ and CD14^+^) and neutrophils (G1^+^ MHCII^-^ or G1^+^ MHCII^+^) in the blood, and the measurement of reactive oxygen species (ROS) production and phagocytosis by these immune cells. etc.’, and **Fig. S4** entitled ‘Flow cytometry gating strategy for the identification of monocytes (MCHII^+^ and CD14^+^) and neutrophils (G1^+^ MHCII^-^ or G1^+^ MHCII^+^) in the milk and the measurement of reactive oxygen species (ROS) production and phagocytosis by these immune cells. etc.’.

The data that support the findings of this study are available from Biodevas Laboratoires and INRAE but restrictions apply to the availability of these data, which were used under license for the current study, and so are not publicly available. Data are however available from the authors upon reasonable request and with the permission of Biodevas Laboratoires and INRAE.

## Ethics approval and consent to participate

Blood and milk samples were collected before and after the LPS challenge infusion in one udder quarter at the Experimental Farm of IE PL, INRAE, Dairy Nutrition and Physiology; (Le Rheu, Brittany, France), and LPS injections were performed with the consent of the Rennes Ethics Committee on Animal Experimentation and the French Ministry for Higher Education, Research and Innovation and performed in compliance with all applicable provisions established by European Directive 2010/63/UE. The authorization number is # 37759-2022062117354678_v2. The study was built and reported in accordance with ARRIVE guidelines (https://arriveguidelines.org).

## Consent for publication

Not Applicable.

