## Supplementary tables for "Nutritional vitamin E or plant extracts affect the immune response and mammary epithelium integrity during intramammary lipopolysaccharide challenge in early lactation"

**Table S1: Plasma energy metabolism indicators, vitamins, and minerals in control unsupplemented group (n = 11), vitamin E supplemented group (n = 13), and plant extract supplemented group (n = 12) dairy cows before supplementation and during LPS challenge time (h). Energy metabolism indicators in plasma such as non-esterified fatty acids (NEFA), β-hydroxybutyrate (BHB), urea, and glucose, and plasma minerals (Ca, K, Mg, Na, P, Pi, S) were measured before LPS challenge (-20 h) and after LPS challenge times (4 h, 28 h and 76 h). Plasma vitamins were measured before supplementation (in week 5 after calving), before LPS challenge (-20 h) and after LPS challenge times (4 h, 28 h, 52 h and 76 h).**

| Item[1] | Parity | Control | | | | | | | Vitamin E | | | | | | | Plant extracts | | | | | | | *P*-value[2] | | | | | | |
|---|---|---|---|---|---|---|---|---|---|---|---|---|---|---|---|---|---|---|---|---|---|---|---|---|---|---|---|---|---|
| | | before supp | -20 | 4 | 28 | 52 | 76 | SEM | before supp | -20 | 4 | 28 | 52 | 76 | SEM | before supp | -20 | 4 | 28 | 52 | 76 | SEM | TREAT | TIMES | TREAT x TIMES | PAR | TREAT x PAR | PAR x TIMES | TREAT x PAR x TIMES |
| Indicators of plasma energy metabolism during LPS challenge | | | | | | | | | | | | | | | | | | | | | | | | | | | | | |
| NEFA (µmol/L) | multiparous | NE[3] | 91.9 | 78.6 | 87.8 | NE | 87.0 | 14.03 | NE | 110 | 98.2 | 115 | NE | 98.1 | 12.17 | NE | 101.8 | 85.18 | 89.75 | NE | 92.00 | 12.07 | | | | | | | |
| | primiparous | NE | 126 | 108 | 95.6 | NE | 79.8 | 15.95 | NE | 80.7 | 96.5 | 104 | NE | 92.5 | 15.66 | NE | 103.0 | 111.1 | 103.1 | NE | 95.43 | 17.45 | 0.91 | 0.25 | 0.42 | 0.62 | 0.43 | 0.31 | 0.42 |
| BHB (µmol/L) | multiparous | NE | 2135 | 1722 | 1943 | NE | 2570 | 445.23 | NE | 2260 | 1566 | 1685 | NE | 1860 | 386.48 | NE | 1640 | 1358 | 1908 | NE | 1930 | 381.88 | | | | | | | |
| | primiparous | NE | 1420 | 1379 | 1391 | NE | 1548 | 512.73 | NE | 1878 | 1783 | 1764 | NE | 1705.996 | 500.33 | NE | 1443 | 1021 | 1374 | NE | 1881 | 557.03 | 0.84 | **< 0.001** | 0.12 | 0.38 | 0.78 | 0.57 | 0.41 |
| Glucose (mg/L) | multiparous | NE | 50.6 | 57.2 | 56.1 | NE | 49.3 | 4.16 | NE | 49.38 | 58.33 | 56.56 | NE | 49.89 | 3.61 | NE | 58.69 | 59.56 | 55.39 | NE | 54.21 | 3.58 | | | | | | | |
| | primiparous | NE | 70.5 | 70.6 | 70.4 | NE | 69.3 | 4.75 | NE | 61.92 | 70.09 | 67.90 | NE | 63.65 | 4.65 | NE | 69.87 | 72.23 | 73.58 | NE | 64.15 | 5.18 | 0.58 | **< 0.001** | 0.52 | **< 0.001** | 0.80 | 0.90 | 0.58 |
| Urea (mg/dL) | multiparous | NE | 195 | 193 | 193 | NE | 190 | 14.91 | NE | 188 | 214 | 196 | NE | 201 | 12.94 | NE | 212 | 211 | 198 | NE | 200 | 12.83 | | | | | | | |
| | primiparous | NE | 191 | 177 | 164 | NE | 168 | 16.93 | NE | 144 | 141 | 143 | NE | 153 | 16.63 | NE | 149 | 139 | 117 | NE | 134 | 18.54 | 0.50 | 0.27 | 0.56 | **< 0.001** | 0.11 | 0.48 | 0.93 |
| Plasma vitamins during LPS challenge | | | | | | | | | | | | | | | | | | | | | | | | | | | | | |
| α-tocopherol (µg/mL) 4 | multiparous | 4.87 | 4.75 | 4.37 | 4.79 | 4.73 | 4.86 | 0.672 | 4.24 | 6.65 | 6.35 | 6.02 | 6.35 | 6.45 | 0.683 | 4.88 | 5.22 | 4.56 | 5.26 | 5.23 | 5.39 | 0.672 | | | | | | | |
| | primiparous | 2.93 | 5.01 | 4.56 | 4.96 | 4.86 | 5.31 | 0.964 | 3.88 | 6.77 | 6.69 | 6.67 | 7.21 | 6.13 | 0.957 | 2.92 | 5.24 | 4.90 | 5.18 | 5.08 | 5.13 | 0.696 | **< 0.001** | **< 0.001** | **0.03** | 0.77 | 0.69 | **< 0.001** | 0.67 |
| γ-tocopherol (µg/mL) | multiparous | 0.74 | 0.67 | 0.75 | 0.77 | 0.63 | 0.69 | 0.179 | 0.61 | 0.78 | 0.76 | 0.67 | 0.72 | 0.73 | 0.178 | 0.66 | 0.80 | 0.67 | 0.83 | 0.60 | 0.74 | 0.179 | | | | | | | |
| | primiparous | 0.53 | 0.79 | 0.54 | 0.68 | 0.80 | 0.66 | 0.250 | 0.67 | 0.62 | 0.67 | 0.77 | 0.64 | 0.56 | 0.250 | 0.47 | 0.69 | 0.83 | 0.91 | 0.66 | 0.73 | 0.178 | 0.81 | 0.19 | 0.94 | 0.46 | 0.87 | 0.88 | 0.59 |
| retinol (µg/mL) | multiparous | 0.58 | 0.62 | 0.58 | 0.65 | 0.58 | 0.63 | 0.062 | 0.57 | 0.63 | 0.60 | 0.59 | 0.62 | 0.66 | 0.062 | 0.63 | 0.64 | 0.59 | 0.63 | 0.59 | 0.64 | 0.062 | | | | | | | |
| | primiparous | 0.60 | 0.63 | 0.56 | 0.60 | 0.71 | 0.65 | 0.087 | 0.57 | 0.61 | 0.65 | 0.62 | 0.59 | 0.58 | 0.087 | 0.52 | 0.59 | 0.63 | 0.58 | 0.57 | 0.56 | 0.062 | 0.67 | 0.38 | 0.96 | 0.42 | 0.22 | 0.70 | 0.70 |
| Plasma minerals during LPS challenge | | | | | | | | | | | | | | | | | | | | | | | | | | | | | |
| Calcium (mg/L) | multiparous | NE | 97.1 | 89.8 | 95.0 | NE | 96.1 | 2.92 | NE | 103.7 | 94.9 | 94.1 | NE | 98.7 | 2.89 | NE | 100 | 101 | 98 | NE | 104 | 3.36 | | | | | | | |
| | primiparous | NE | 101 | 96.5 | 96.3 | NE | 105 | 3.74 | NE | 95.8 | 96.7 | 102.7 | NE | 103 | 4.17 | NE | 98 | 98 | 94 | NE | 101 | 3.80 | 0.71 | **< 0.001** | 0.65 | 0.60 | 0.29 | 0.34 | 0.16 |
| Potassium (mg/L) | multiparous | NE | 141 | 126 | 137 | NE | 129 | 5.84 | NE | 146 | 138 | 129 | NE | 133 | 5.81 | NE | 141 | 141 | 139 | NE | 136 | 6.74 | | | | | | | |
| | primiparous | NE | 135 | 128 | 141 | NE | 127 | 7.48 | NE | 117 | 123 | 128 | NE | 125 | 8.34 | NE | 129 | 148 | 124 | NE | 117 | 7.57 | 0.61 | 0.19 | 0.10 | **0.06** | 0.37 | 0.20 | 0.34 |
| Magnesium (mg/L) | multiparous | NE | 22.8 | 22.9 | 23.8 | NE | 24.0 | 0.90 | NE | 24.98 | 24.65 | 24.82 | NE | 23.6 | 0.89 | NE | 25.2 | 25.9 | 25.2 | NE | 24.3 | 1.04 | | | | | | | |
| | primiparous | NE | 25.7 | 25.6 | 26.7 | NE | 25.8 | 1.15 | NE | 22.26 | 23.12 | 24.73 | NE | 24.5 | 1.29 | NE | 24.3 | 24.4 | 23.6 | NE | 23.0 | 1.17 | 0.73 | 0.57 | 0.57 | 0.84 | **0.04** | 0.84 | 0.51 |
| Sodium (mg/L) | multiparous | NE | 3190 | 3098 | 3151 | NE | 3053 | 103.25 | NE | 3314 | 3211 | 3028 | NE | 3195 | 102.57 | NE | 3147 | 3334 | 3139 | NE | 3148 | 119.07 | | | | | | | |
| | primiparous | NE | 2945 | 3010 | 3121 | NE | 3014 | 132.33 | NE | 2989 | 2906 | 3174 | NE | 3099 | 147.59 | NE | 2976 | 3141 | 2830 | NE | 2828 | 134.21 | 0.83 | 0.75 | 0.18 | 0.04 | 0.70 | 0.46 | 0.35 |
| Phosphorus (mg/L) | multiparous | NE | 144 | 130 | 141 | NE | 137 | 8.81 | NE | 147.3 | 128.3 | 126.4 | NE | 144.7 | 8.71 | NE | 140 | 139 | 141 | NE | 150 | 10.15 | | | | | | | |
| | primiparous | NE | 137 | 131 | 141 | NE | 139 | 11.38 | NE | 141.0 | 129.2 | 149.9 | NE | 141.5 | 12.68 | NE | 150 | 145 | 140 | NE | 143 | 11.64 | 0.80 | **0.01** | 0.55 | 0.85 | 0.97 | 0.33 | 0.14 |
| Inorganic phosphorus (mg/L) | multiparous | NE | 45.1 | 38.2 | 41.5 | NE | 37.0 | 3.64 | NE | 40.9 | 27.9 | 29.8 | NE | 41.1 | 3.61 | NE | 37.4 | 30.4 | 35.5 | NE | 40.5 | 4.19 | | | | | | | |
| | primiparous | NE | 42.0 | 39.0 | 47.5 | NE | 40.7 | 4.68 | NE | 47.7 | 36.8 | 41.5 | NE | 38.8 | 5.21 | NE | 52.0 | 44.2 | 44.6 | NE | 45.0 | 4.76 | 0.54 | **< 0.001** | 0.22 | **0.04** | 0.47 | 0.25 | 0.35 |
| Suffer (mg/L) | multiparous | NE | 1205 | 1113 | 1210 | NE | 1213 | 43.35 | NE | 1269 | 1197 | 1181 | NE | 1205 | 43.06 | NE | 1222 | 1264 | 1209 | NE | 1265 | 49.99 | | | | | | | |
| | primiparous | NE | 1143 | 1100 | 1104 | NE | 1142 | 55.58 | NE | 1072 | 1061 | 1166 | NE | 1157 | 61.98 | NE | 1115 | 1122 | 1084 | NE | 1108 | 56.39 | 0.87 | 0.43 | 0.67 | **< 0.001** | 0.66 | 0.86 | 0.33 |

1 Plasma samples were collected before supplementation (in week 5 after calving), before LPS challenge (-20 h) and after LPS challenge times (4 h, 28 h, 52 h and 76 h).
2 Probability: TREAT = effect of treatment (control vs vitamin E vs plant extract group); TIMES = effect of times before supplementation (5 week before calving) vs before LPS challenge (-20 hours) vs after LPS challenge (vs 4 vs 28 vs 52 vs 76 hours); TREAT × TIMES = the interaction between treatment and times; PAR = effect of parity (primiparous vs multiparous); PAR × TIMES = interaction between parity and times; TREAT × PAR = interaction between treatment and parity; TREAT × PAR × TIMES = interaction between treatment and parity and times.
3 NE =Not estimated because no collection.
4 Plasma alphatocopherol was not vary during LPS challenge, even if we did a multiparametric test ANOVA without time point before supplementation (data not shown, time effect P = 0.53, Treat x time interaction  P= 0.71, Treat effect P < 0.001)

**Table S2: Abundance of antioxidant function mRNA, immune function mRNA and gene cell death mRNA determined by real-time quantitative RT-PCR in blood in control unsupplemented group (n = 11), vitamin E supplemented group (n = 13), and plant extracts supplemented group (n = 12) dairy cows before (- 20 h) and after LPS challenge times (4 h, 28 h and 76 h).**

| Item [1] | Parity | control | | | | | vitamin E | | | | | Plant extracts | | | | | *P*-value [2] | | | | | | |
|---|---|---|---|---|---|---|---|---|---|---|---|---|---|---|---|---|---|---|---|---|---|---|---|
| | | -20 | 4 | 28 | 76 | SEM | -20 | 4 | 28 | 76 | SEM | -20 | 4 | 28 | 76 | SEM | TREAT | TIMES | TREAT x TIMES | PAR | TREAT x PAR | PAR x TIMES | TREAT x PAR x TIMES |
| **Redox gene expression** | | | | | | | | | | | | | | | | | | | | | | | |
| NFE2L2 | multiparous | 4.73 | 4.72 | 4.72 | 4.84 | 0.09 | 4.86 | 4.75 | 4.81 | 4.87 | 0.08 | 5.05 | 4.62 | 4.71 | 4.86 | 0.08 | | | | | | | |
| | primiparous | 4.95 | 4.71 | 4.81 | 4.92 | 0.11 | 4.84 | 4.64 | 5.01 | 4.87 | 0.11 | 4.89 | 4.64 | 4.94 | 4.85 | 0.12 | 0.90 | 0.00 | 0.30 | 0.50 | 0.83 | 0.14 | 0.36 |
| GPX3 | multiparous | 4.79 | 5.09 | 4.87 | 4.99 | 0.12 | 5.05 | 5.00 | 5.02 | 4.87 | 0.10 | 5.10 | 4.93 | 4.91 | 4.83 | 0.10 | | | | | | | |
| | primiparous | 5.00 | 4.95 | 5.00 | 4.97 | 0.14 | 4.99 | 4.87 | 5.18 | 4.92 | 0.13 | 4.96 | 5.05 | 5.01 | 4.92 | 0.15 | 0.92 | 0.62 | 0.47 | 0.64 | 0.95 | 0.60 | 0.61 |
| SOD2 | multiparous | 6.90 | 7.24 | 6.87 | 7.06 | 0.09 | 7.09 | 7.39 | 7.14 | 7.07 | 0.08 | 7.06 | 7.23 | 7.07 | 7.08 | 0.08 | | | | | | | |
| | primiparous | 7.06 | 7.39 | 7.11 | 7.06 | 0.10 | 7.04 | 7.13 | 7.21 | 7.00 | 0.10 | 7.06 | 7.48 | 7.07 | 6.93 | 0.11 | 0.71 | 0.00 | 0.26 | 0.61 | 0.26 | 0.27 | 0.13 |
| HMOX1 | multiparous | 6.11 | 5.17 | 6.10 | 6.25 | 0.11 | 6.18 | 5.37 | 6.14 | 6.21 | 0.10 | 6.22 | 5.44 | 6.19 | 6.27 | 0.10 | | | | | | | |
| | primiparous | 6.00 | 5.58 | 6.14 | 6.15 | 0.12 | 5.94 | 5.59 | 6.10 | 6.21 | 0.12 | 6.04 | 5.39 | 5.99 | 6.14 | 0.14 | 0.88 | 0.00 | 0.96 | 0.61 | 0.36 | 0.03 | 0.80 |
| SOD1 | multiparous | 6.93 | 7.03 | 6.92 | 7.16 | 0.12 | 7.18 | 7.21 | 7.08 | 7.00 | 0.11 | 7.36 | 6.94 | 6.98 | 6.95 | 0.11 | | | | | | | |
| | primiparous | 7.03 | 7.30 | 7.13 | 7.04 | 0.14 | 7.08 | 6.96 | 7.34 | 6.99 | 0.14 | 6.97 | 6.99 | 6.97 | 6.97 | 0.15 | 0.46 | 0.76 | 0.22 | 0.98 | 0.41 | 0.28 | 0.27 |
| CAT | multiparous | 6.34 | 6.24 | 6.33 | 6.48 | 0.07 | 6.53 | 6.31 | 6.40 | 6.42 | 0.06 | 6.62 | 6.24 | 6.39 | 6.40 | 0.06 | | | | | | | |
| | primiparous | 6.49 | 6.45 | 6.46 | 6.47 | 0.08 | 6.44 | 6.27 | 6.53 | 6.45 | 0.08 | 6.44 | 6.26 | 6.38 | 6.42 | 0.09 | 0.84 | 0.00 | 0.31 | 0.42 | 0.18 | 0.39 | 0.43 |
| NQO1 | multiparous | 3.52 | 3.42 | 3.57 | 3.98 | 0.21 | 3.70 | 3.68 | 3.50 | 3.77 | 0.18 | 3.64 | 3.96 | 3.92 | 3.63 | 0.18 | | | | | | | |
| | primiparous | 3.89 | 3.66 | 3.40 | 3.81 | 0.23 | 3.85 | 3.34 | 3.76 | 3.62 | 0.23 | 3.83 | 3.87 | 3.71 | 3.45 | 0.26 | 0.57 | 0.86 | 0.22 | 0.93 | 0.81 | 0.42 | 0.77 |
| SOD3 | multiparous | 4.09 | 4.23 | 3.77 | 4.04 | 0.26 | 4.38 | 4.49 | 4.34 | 4.44 | 0.23 | 4.28 | 4.09 | 4.08 | 4.13 | 0.22 | | | | | | | |
| | primiparous | 4.01 | 4.43 | 3.85 | 4.10 | 0.30 | 4.19 | 4.22 | 4.32 | 4.23 | 0.29 | 2.99 | 3.35 | 3.30 | 3.44 | 0.33 | 0.04 | 0.19 | 0.40 | 0.12 | 0.13 | 0.43 | 0.82 |
| TXNRD1 | multiparous | 6.46 | 6.48 | 6.44 | 6.60 | 0.09 | 6.63 | 6.61 | 6.51 | 6.50 | 0.08 | 6.83 | 6.50 | 6.48 | 6.51 | 0.08 | | | | | | | |
| | primiparous | 6.47 | 6.63 | 6.54 | 6.51 | 0.10 | 6.45 | 6.44 | 6.57 | 6.48 | 0.10 | 6.47 | 6.47 | 6.44 | 6.49 | 0.11 | 0.98 | 0.79 | 0.42 | 0.26 | 0.28 | 0.21 | 0.61 |
| GPX1 | multiparous | 7.51 | 7.19 | 7.43 | 7.57 | 0.07 | 7.54 | 7.26 | 7.48 | 7.57 | 0.06 | 7.64 | 7.34 | 7.51 | 7.59 | 0.06 | | | | | | | |
| | primiparous | 7.50 | 7.40 | 7.53 | 7.55 | 0.08 | 7.47 | 7.34 | 7.54 | 7.58 | 0.08 | 7.55 | 7.36 | 7.53 | 7.57 | 0.08 | 0.41 | 0.00 | 0.98 | 0.47 | 0.55 | 0.18 | 0.98 |
| **Immune gene expression** | | | | | | | | | | | | | | | | | | | | | | | |
| TNF | multiparous | 5.21 | 5.12 | 5.10 | 5.24 | 0.09 | 5.24 | 5.15 | 5.13 | 5.20 | 0.08 | 5.35 | 5.17 | 5.19 | 5.20 | 0.08 | | | | | | | |
| | primiparous | 5.19 | 5.12 | 5.19 | 4.77 | 0.10 | 5.18 | 5.21 | 5.09 | 5.08 | 0.10 | 5.21 | 5.20 | 5.15 | 5.14 | 0.11 | 0.27 | 0.10 | 0.86 | 0.16 | 0.82 | 0.08 | 0.36 |
| TLR2 | multiparous | 6.07 | 6.14 | 6.00 | 6.05 | 0.07 | 6.04 | 6.22 | 6.08 | 6.23 | 0.06 | 6.13 | 6.16 | 6.08 | 6.18 | 0.06 | | | | | | | |
| | primiparous | 6.17 | 6.28 | 6.02 | 6.25 | 0.08 | 6.17 | 6.19 | 6.19 | 6.23 | 0.08 | 6.22 | 6.32 | 6.06 | 6.18 | 0.09 | 0.48 | 0.00 | 0.60 | 0.05 | 0.73 | 0.83 | 0.51 |
| STAT2 | multiparous | 4.89 | 5.00 | 4.84 | 5.05 | 0.08 | 5.02 | 5.11 | 5.01 | 5.05 | 0.07 | 5.15 | 5.07 | 4.97 | 5.02 | 0.07 | | | | | | | |
| | primiparous | 4.85 | 5.10 | 5.02 | 4.91 | 0.09 | 4.90 | 4.97 | 5.04 | 4.96 | 0.09 | 4.94 | 5.16 | 4.91 | 4.92 | 0.10 | 0.38 | 0.12 | 0.60 | 0.31 | 0.43 | 0.21 | 0.69 |
| SLC2A1 | multiparous | 3.54 | 3.87 | 3.69 | 3.81 | 0.18 | 3.79 | 4.09 | 3.80 | 3.69 | 0.15 | 4.29 | 3.79 | 3.69 | 3.54 | 0.15 | | | | | | | |
| | primiparous | 3.44 | 3.98 | 3.60 | 3.48 | 0.20 | 3.48 | 3.49 | 3.69 | 3.48 | 0.20 | 3.46 | 3.57 | 3.42 | 3.41 | 0.22 | 0.89 | 0.18 | 0.25 | 0.00 | 0.35 | 0.64 | 0.40 |
| SELL | multiparous | 7.42 | 7.47 | 7.38 | 7.47 | 0.06 | 7.46 | 7.56 | 7.49 | 7.46 | 0.05 | 7.56 | 7.54 | 7.47 | 7.50 | 0.05 | | | | | | | |
| | primiparous | 7.45 | 7.51 | 7.47 | 7.46 | 0.06 | 7.38 | 7.43 | 7.46 | 7.46 | 0.06 | 7.46 | 7.62 | 7.44 | 7.40 | 0.07 | 0.42 | 0.07 | 0.62 | 0.55 | 0.38 | 0.75 | 0.49 |
| IL23A | multiparous | 4.87 | 4.84 | 5.01 | 5.02 | 0.11 | 4.87 | 4.54 | 4.85 | 4.94 | 0.10 | 5.02 | 4.85 | 4.99 | 4.98 | 0.10 | | | | | | | |
| | primiparous | 5.07 | 4.78 | 4.91 | 5.07 | 0.12 | 5.02 | 4.96 | 4.91 | 4.98 | 0.12 | 4.97 | 4.97 | 5.06 | 4.90 | 0.14 | 0.30 | 0.06 | 0.88 | 0.18 | 0.34 | 0.58 | 0.59 |
| CD80 | multiparous | 6.55 | 6.80 | 6.54 | 6.62 | 0.06 | 6.69 | 6.91 | 6.61 | 6.59 | 0.05 | 6.76 | 6.80 | 6.59 | 6.56 | 0.05 | | | | | | | |
| | primiparous | 6.55 | 6.81 | 6.53 | 6.55 | 0.07 | 6.51 | 6.59 | 6.68 | 6.50 | 0.07 | 6.54 | 6.82 | 6.53 | 6.49 | 0.08 | 0.92 | 0.00 | 0.09 | 0.06 | 0.46 | 0.16 | 0.01 |
| MMP14 | multiparous | 3.62 | 3.20 | 3.50 | 3.86 | 0.22 | 4.05 | 3.07 | 3.75 | 3.94 | 0.19 | 3.54 | 2.90 | 3.77 | 3.64 | 0.19 | | | | | | | |
| | primiparous | 3.65 | 3.60 | 3.71 | 3.61 | 0.25 | 3.80 | 3.32 | 3.49 | 3.81 | 0.25 | 3.85 | 2.36 | 3.93 | 3.95 | 0.28 | 0.37 | 0.00 | 0.05 | 0.84 | 0.66 | 1.00 | 0.24 |
| CXCL8 | multiparous | 6.76 | 6.98 | 6.80 | 6.95 | 0.14 | 7.06 | 7.22 | 6.99 | 6.87 | 0.12 | 7.23 | 7.07 | 7.00 | 6.91 | 0.12 | | | | | | | |
| | primiparous | 6.82 | 7.18 | 6.90 | 6.83 | 0.16 | 6.79 | 6.82 | 6.99 | 6.72 | 0.15 | 6.94 | 7.07 | 6.92 | 6.86 | 0.17 | 0.69 | 0.01 | 0.22 | 0.40 | 0.50 | 0.49 | 0.30 |
| IL1A | multiparous | 4.71 | 5.52 | 4.52 | 4.70 | 0.13 | 4.89 | 5.55 | 4.62 | 4.57 | 0.11 | 4.84 | 5.39 | 4.47 | 4.55 | 0.11 | | | | | | | |
| | primiparous | 4.78 | 5.21 | 4.56 | 4.51 | 0.14 | 4.60 | 5.24 | 4.84 | 4.66 | 0.14 | 4.70 | 5.44 | 4.52 | 4.06 | 0.16 | 0.38 | 0.00 | 0.13 | 0.21 | 0.93 | 0.08 | 0.05 |
| S100A7 | multiparous | 5.43 | 4.74 | 5.22 | 5.79 | 0.18 | 5.72 | 5.03 | 5.42 | 5.73 | 0.15 | 5.79 | 4.60 | 5.39 | 5.56 | 0.15 | | | | | | | |
| | primiparous | 5.45 | 5.37 | 5.39 | 5.61 | 0.20 | 5.64 | 4.83 | 5.73 | 5.64 | 0.20 | 5.52 | 4.79 | 5.30 | 5.64 | 0.22 | 0.45 | 0.00 | 0.26 | 0.69 | 0.69 | 0.29 | 0.25 |
| TLR4 | multiparous | 5.91 | 6.24 | 5.78 | 5.97 | 0.07 | 5.99 | 6.23 | 5.91 | 5.98 | 0.06 | 6.00 | 6.19 | 5.84 | 5.93 | 0.06 | | | | | | | |
| | primiparous | 5.97 | 6.16 | 5.94 | 5.94 | 0.08 | 5.90 | 6.07 | 6.00 | 5.96 | 0.08 | 6.00 | 6.34 | 5.89 | 5.91 | 0.09 | 0.86 | 0.00 | 0.28 | 0.84 | 0.56 | 0.22 | 0.37 |
| STAT5A | multiparous | 5.94 | 5.78 | 5.89 | 5.93 | 0.06 | 5.89 | 5.78 | 5.92 | 5.93 | 0.05 | 6.02 | 5.95 | 5.94 | 6.03 | 0.05 | | | | | | | |
| | primiparous | 5.97 | 5.87 | 6.01 | 6.01 | 0.06 | 5.91 | 6.02 | 5.88 | 6.01 | 0.06 | 6.02 | 6.02 | 6.02 | 5.98 | 0.07 | 0.03 | 0.12 | 0.58 | 0.04 | 0.65 | 0.30 | 0.45 |
| CCR6 | multiparous | 5.37 | 5.26 | 5.35 | 5.51 | 0.07 | 5.45 | 5.41 | 5.46 | 5.45 | 0.06 | 5.54 | 5.31 | 5.37 | 5.49 | 0.06 | | | | | | | |
| | primiparous | 5.34 | 5.44 | 5.47 | 5.45 | 0.08 | 5.37 | 5.32 | 5.42 | 5.42 | 0.08 | 5.36 | 5.32 | 5.26 | 5.48 | 0.09 | 0.93 | 0.01 | 0.25 | 0.57 | 0.47 | 0.31 | 0.47 |
| CX3CR1 | multiparous | 6.56 | 5.94 | 6.47 | 6.67 | 0.08 | 6.66 | 6.07 | 6.48 | 6.62 | 0.07 | 6.61 | 6.08 | 6.52 | 6.63 | 0.07 | | | | | | | |
| | primiparous | 6.59 | 6.31 | 6.51 | 6.59 | 0.09 | 6.59 | 6.27 | 6.57 | 6.60 | 0.09 | 6.55 | 6.17 | 6.48 | 6.54 | 0.10 | 0.78 | 0.00 | 0.99 | 0.41 | 0.59 | 0.00 | 0.80 |
| CD86 | multiparous | 6.23 | 5.78 | 6.16 | 6.47 | 0.10 | 6.49 | 6.03 | 6.26 | 6.36 | 0.09 | 6.50 | 5.89 | 6.22 | 6.29 | 0.09 | | | | | | | |
| | primiparous | 6.30 | 6.23 | 6.30 | 6.33 | 0.12 | 6.34 | 5.91 | 6.41 | 6.35 | 0.11 | 6.31 | 5.81 | 6.10 | 6.31 | 0.13 | 0.43 | 0.00 | 0.32 | 0.99 | 0.28 | 0.32 | 0.10 |
| TNFRSF18 | multiparous | 4.58 | 3.88 | 4.36 | 4.48 | 0.17 | 4.11 | 4.03 | 4.33 | 4.43 | 0.15 | 4.52 | 4.50 | 4.42 | 4.60 | 0.15 | | | | | | | |
| | primiparous | 4.42 | 4.25 | 4.46 | 4.53 | 0.19 | 4.33 | 4.49 | 4.20 | 4.56 | 0.19 | 4.58 | 4.33 | 4.52 | 4.49 | 0.22 | 0.20 | 0.05 | 0.48 | 0.40 | 0.63 | 0.69 | 0.54 |
| IL1B | multiparous | 6.82 | 7.20 | 6.76 | 6.83 | 0.07 | 6.90 | 7.16 | 6.87 | 6.86 | 0.06 | 6.88 | 7.08 | 6.85 | 6.86 | 0.06 | | | | | | | |
| | primiparous | 6.97 | 7.01 | 6.81 | 6.87 | 0.08 | 6.90 | 7.01 | 7.01 | 6.90 | 0.08 | 7.02 | 7.31 | 6.97 | 6.83 | 0.09 | 0.51 | 0.00 | 0.30 | 0.37 | 0.57 | 0.14 | 0.05 |
| S100A8 | multiparous | 7.96 | 8.53 | 7.96 | 8.33 | 0.13 | 8.25 | 8.74 | 8.25 | 8.16 | 0.11 | 8.31 | 8.46 | 8.16 | 8.16 | 0.11 | | | | | | | |
| | primiparous | 8.08 | 8.69 | 8.27 | 8.13 | 0.15 | 8.08 | 8.23 | 8.49 | 8.08 | 0.14 | 8.07 | 8.52 | 8.11 | 8.05 | 0.16 | 0.77 | 0.00 | 0.18 | 0.58 | 0.34 | 0.16 | 0.14 |
| FAS | multiparous | 6.38 | 6.38 | 6.23 | 6.47 | 0.08 | 6.48 | 6.50 | 6.40 | 6.41 | 0.07 | 6.54 | 6.36 | 6.33 | 6.41 | 0.07 | | | | | | | |
| | primiparous | 6.34 | 6.45 | 6.36 | 6.34 | 0.09 | 6.33 | 6.31 | 6.44 | 6.35 | 0.09 | 6.40 | 6.46 | 6.30 | 6.30 | 0.10 | 0.86 | 0.34 | 0.48 | 0.41 | 0.69 | 0.23 | 0.54 |
| CCR7 | multiparous | 6.42 | 6.46 | 6.43 | 6.38 | 0.08 | 6.30 | 6.45 | 6.39 | 6.39 | 0.07 | 6.43 | 6.62 | 6.37 | 6.51 | 0.07 | | | | | | | |
| | primiparous | 6.46 | 6.49 | 6.49 | 6.51 | 0.09 | 6.35 | 6.66 | 6.19 | 6.49 | 0.09 | 6.44 | 6.73 | 6.44 | 6.49 | 0.10 | 0.15 | 0.00 | 0.19 | 0.29 | 0.97 | 0.44 | 0.30 |
| CXCR4 | multiparous | 6.37 | 6.59 | 6.45 | 6.23 | 0.09 | 6.37 | 6.72 | 6.44 | 6.45 | 0.07 | 6.41 | 6.74 | 6.47 | 6.53 | 0.07 | | | | | | | |
| | primiparous | 6.54 | 6.64 | 6.45 | 6.51 | 0.10 | 6.48 | 6.70 | 6.50 | 6.59 | 0.09 | 6.51 | 6.87 | 6.60 | 6.51 | 0.11 | 0.12 | 0.00 | 0.68 | 0.04 | 0.87 | 0.78 | 0.65 |
| CCL5 | multiparous | 5.45 | 4.72 | 5.27 | 5.24 | 0.15 | 5.21 | 4.63 | 5.07 | 5.29 | 0.13 | 5.34 | 5.13 | 5.36 | 5.45 | 0.13 | | | | | | | |
| | primiparous | 5.49 | 4.89 | 5.30 | 5.42 | 0.17 | 5.62 | 5.33 | 5.21 | 5.54 | 0.17 | 5.46 | 5.16 | 5.38 | 5.35 | 0.18 | 0.47 | 0.00 | 0.42 | 0.04 | 0.13 | 0.52 | 0.81 |
| S100A9 | multiparous | 7.93 | 8.53 | 8.03 | 8.08 | 0.07 | 8.03 | 8.57 | 8.15 | 8.07 | 0.06 | 8.03 | 8.52 | 8.10 | 8.12 | 0.06 | | | | | | | |
| | primiparous | 8.05 | 8.43 | 8.12 | 8.08 | 0.08 | 7.94 | 8.33 | 8.23 | 8.03 | 0.07 | 8.01 | 8.58 | 8.10 | 7.99 | 0.08 | 0.88 | 0.00 | 0.25 | 0.58 | 0.53 | 0.12 | 0.19 |
| SOCS3 | multiparous | 5.81 | 6.45 | 5.63 | 5.79 | 0.09 | 5.65 | 6.43 | 5.68 | 5.76 | 0.08 | 5.86 | 6.49 | 5.57 | 5.83 | 0.08 | | | | | | | |
| | primiparous | 5.60 | 6.31 | 5.64 | 5.66 | 0.10 | 5.56 | 6.35 | 5.47 | 5.66 | 0.10 | 5.68 | 6.68 | 5.50 | 5.73 | 0.12 | 0.36 | 0.00 | 0.20 | 0.12 | 0.79 | 0.45 | 0.42 |
| SPI1 | multiparous | 7.31 | 7.42 | 7.31 | 7.61 | 0.15 | 7.62 | 7.65 | 7.50 | 7.42 | 0.13 | 7.83 | 7.34 | 7.42 | 7.37 | 0.13 | | | | | | | |
| | primiparous | 7.38 | 7.73 | 7.52 | 7.37 | 0.17 | 7.49 | 7.23 | 7.84 | 7.37 | 0.17 | 7.38 | 7.38 | 7.43 | 7.36 | 0.18 | 0.62 | 0.72 | 0.20 | 0.71 | 0.53 | 0.20 | 0.17 |
| ABCG2 | multiparous | 5.04 | 5.07 | 5.03 | 5.10 | 0.08 | 5.17 | 5.06 | 5.03 | 5.04 | 0.07 | 5.32 | 5.02 | 5.12 | 5.05 | 0.07 | | | | | | | |
| | primiparous | 5.24 | 5.18 | 5.04 | 5.21 | 0.08 | 5.17 | 5.01 | 5.26 | 5.07 | 0.08 | 5.13 | 5.02 | 5.13 | 5.11 | 0.09 | 0.95 | 0.07 | 0.33 | 0.21 | 0.25 | 0.78 | 0.31 |
| CCL20 | multiparous | 4.38 | 4.88 | 4.33 | 4.22 | 0.15 | 4.48 | 4.89 | 4.78 | 4.51 | 0.13 | 4.59 | 4.57 | 4.54 | 4.48 | 0.13 | | | | | | | |
| | primiparous | 4.46 | 4.45 | 4.23 | 4.52 | 0.17 | 4.48 | 4.87 | 4.55 | 4.44 | 0.17 | 4.54 | 4.96 | 4.71 | 4.59 | 0.19 | 0.11 | 0.00 | 0.68 | 0.87 | 0.47 | 0.76 | 0.21 |
| STAT4 | multiparous | 5.96 | 6.06 | 5.98 | 6.02 | 0.05 | 6.13 | 6.09 | 6.08 | 6.03 | 0.05 | 6.14 | 5.99 | 6.04 | 5.99 | 0.04 | | | | | | | |
| | primiparous | 6.11 | 6.15 | 6.11 | 6.09 | 0.06 | 6.06 | 5.99 | 6.13 | 6.00 | 0.06 | 6.02 | 6.03 | 5.98 | 5.96 | 0.06 | 0.38 | 0.22 | 0.19 | 0.73 | 0.10 | 0.79 | 0.37 |
| C5AR1 | multiparous | 6.75 | 6.82 | 6.75 | 6.59 | 0.13 | 6.50 | 6.65 | 6.64 | 6.63 | 0.12 | 6.52 | 6.99 | 6.72 | 6.78 | 0.11 | | | | | | | |
| | primiparous | 6.71 | 6.59 | 6.65 | 6.62 | 0.15 | 6.56 | 6.79 | 6.46 | 6.75 | 0.15 | 6.77 | 7.00 | 6.84 | 6.69 | 0.16 | 0.11 | 0.11 | 0.41 | 0.90 | 0.62 | 0.80 | 0.69 |
| BAX | multiparous | 5.94 | 5.89 | 5.88 | 5.88 | 0.05 | 5.85 | 5.88 | 5.90 | 6.00 | 0.04 | 5.93 | 5.92 | 5.94 | 6.00 | 0.04 | | | | | | | |
| | primiparous | 5.96 | 5.87 | 5.95 | 6.03 | 0.05 | 5.93 | 5.94 | 6.01 | 6.02 | 0.05 | 6.01 | 5.89 | 5.92 | 6.00 | 0.06 | 0.63 | 0.02 | 0.57 | 0.10 | 0.59 | 0.69 | 0.58 |
| NLRP3 | multiparous | 6.11 | 6.09 | 6.00 | 6.07 | 0.07 | 6.12 | 5.98 | 6.06 | 6.20 | 0.06 | 6.09 | 6.05 | 6.02 | 6.16 | 0.06 | | | | | | | |
| | primiparous | 6.15 | 5.99 | 6.05 | 6.14 | 0.08 | 6.15 | 6.03 | 5.91 | 6.23 | 0.08 | 6.28 | 6.19 | 6.08 | 6.17 | 0.09 | 0.42 | 0.00 | 0.74 | 0.36 | 0.42 | 0.68 | 0.61 |
| AHR | multiparous | 6.07 | 6.00 | 5.97 | 6.17 | 0.07 | 6.24 | 6.16 | 6.11 | 6.21 | 0.06 | 6.30 | 6.07 | 6.10 | 6.19 | 0.06 | | | | | | | |
| | primiparous | 6.14 | 6.06 | 6.08 | 6.15 | 0.07 | 6.11 | 6.02 | 6.12 | 6.14 | 0.07 | 6.12 | 6.03 | 6.02 | 6.15 | 0.08 | 0.53 | 0.00 | 0.69 | 0.43 | 0.33 | 0.44 | 0.57 |
| CD74 | multiparous | 8.02 | 7.75 | 7.93 | 7.93 | 0.08 | 7.97 | 7.75 | 7.95 | 8.09 | 0.07 | 7.84 | 7.93 | 8.01 | 8.09 | 0.07 | | | | | | | |
| | primiparous | 8.07 | 7.84 | 7.95 | 8.09 | 0.09 | 8.06 | 7.91 | 7.83 | 8.10 | 0.09 | 8.11 | 7.90 | 7.96 | 8.11 | 0.10 | 0.53 | 0.00 | 0.58 | 0.14 | 0.87 | 0.28 | 0.56 |
| CCR1 | multiparous | 6.55 | 6.78 | 6.49 | 6.39 | 0.13 | 6.34 | 6.63 | 6.38 | 6.47 | 0.11 | 6.34 | 6.96 | 6.47 | 6.60 | 0.11 | | | | | | | |
| | primiparous | 6.51 | 6.59 | 6.40 | 6.44 | 0.15 | 6.41 | 6.79 | 6.25 | 6.51 | 0.15 | 6.54 | 7.00 | 6.49 | 6.44 | 0.16 | 0.26 | 0.00 | 0.42 | 0.98 | 0.80 | 0.81 | 0.75 |
| NFkB1 | multiparous | 6.09 | 5.87 | 6.02 | 5.88 | 0.10 | 5.94 | 5.78 | 6.00 | 6.10 | 0.09 | 6.02 | 6.06 | 6.09 | 6.13 | 0.09 | | | | | | | |
| | primiparous | 6.21 | 5.99 | 6.11 | 6.20 | 0.11 | 6.19 | 6.17 | 5.96 | 6.22 | 0.11 | 6.29 | 6.20 | 6.26 | 6.17 | 0.12 | 0.09 | 0.18 | 0.55 | 0.00 | 0.98 | 0.58 | 0.43 |
| BCL2 | multiparous | 5.64 | 5.39 | 5.55 | 5.47 | 0.13 | 5.48 | 5.19 | 5.60 | 5.60 | 0.11 | 5.55 | 5.64 | 5.63 | 5.71 | 0.11 | | | | | | | |
| | primiparous | 5.70 | 5.39 | 5.48 | 5.71 | 0.15 | 5.59 | 5.55 | 5.27 | 5.63 | 0.14 | 5.70 | 5.61 | 5.62 | 5.70 | 0.16 | 0.11 | 0.09 | 0.83 | 0.50 | 0.97 | 0.29 | 0.46 |
| STAT1 | multiparous | 6.65 | 6.89 | 6.49 | 6.88 | 0.12 | 6.86 | 7.04 | 6.74 | 6.82 | 0.10 | 7.02 | 6.78 | 6.58 | 6.67 | 0.10 | | | | | | | |
| | primiparous | 6.77 | 7.12 | 6.81 | 6.77 | 0.13 | 6.86 | 6.74 | 6.99 | 6.84 | 0.13 | 6.81 | 6.81 | 6.61 | 6.75 | 0.15 | 0.32 | 0.04 | 0.12 | 0.53 | 0.47 | 0.31 | 0.29 |
| CD14 | multiparous | 6.27 | 5.99 | 6.12 | 6.20 | 0.10 | 6.14 | 6.07 | 6.15 | 6.32 | 0.08 | 6.18 | 6.16 | 6.20 | 6.38 | 0.08 | | | | | | | |
| | primiparous | 6.19 | 6.30 | 6.25 | 6.33 | 0.11 | 6.15 | 6.24 | 6.08 | 6.39 | 0.11 | 6.39 | 6.32 | 6.16 | 6.39 | 0.12 | 0.35 | 0.01 | 0.85 | 0.12 | 0.78 | 0.25 | 0.62 |
| LAP | multiparous | 5.94 | 6.03 | 5.84 | 6.16 | 0.13 | 6.22 | 6.60 | 6.29 | 6.29 | 0.12 | 6.16 | 6.13 | 6.10 | 6.19 | 0.12 | | | | | | | |
| | primiparous | 6.21 | 6.66 | 6.31 | 6.34 | 0.15 | 6.14 | 6.06 | 6.38 | 6.16 | 0.15 | 6.17 | 6.40 | 6.05 | 6.19 | 0.17 | 0.52 | 0.07 | 0.57 | 0.25 | 0.02 | 0.72 | 0.07 |
| CXCL1 | multiparous | 4.37 | 5.20 | 4.23 | 4.49 | 0.18 | 4.57 | 5.45 | 4.18 | 4.50 | 0.16 | 4.45 | 5.20 | 4.12 | 4.36 | 0.16 | | | | | | | |
| | primiparous | 4.53 | 5.34 | 4.02 | 4.44 | 0.21 | 4.13 | 5.01 | 4.39 | 4.46 | 0.20 | 4.34 | 5.46 | 4.05 | 4.42 | 0.23 | 0.92 | 0.00 | 0.92 | 0.59 | 0.47 | 0.94 | 0.37 |
| IL2RA | multiparous | 5.68 | 5.82 | 5.76 | 5.79 | 0.08 | 5.83 | 6.02 | 5.96 | 5.80 | 0.07 | 5.88 | 5.89 | 5.85 | 5.75 | 0.07 | | | | | | | |
| | primiparous | 5.65 | 5.94 | 5.85 | 5.72 | 0.09 | 5.58 | 5.68 | 5.96 | 5.65 | 0.09 | 5.75 | 6.01 | 5.80 | 5.60 | 0.10 | 0.72 | 0.00 | 0.10 | 0.13 | 0.11 | 0.26 | 0.27 |
| GCLM | multiparous | 5.67 | 5.79 | 5.72 | 5.81 | 0.09 | 5.78 | 5.95 | 5.77 | 5.68 | 0.08 | 5.89 | 5.71 | 5.69 | 5.61 | 0.08 | | | | | | | |
| | primiparous | 5.67 | 5.87 | 5.79 | 5.60 | 0.10 | 5.67 | 5.64 | 5.92 | 5.59 | 0.10 | 5.63 | 5.67 | 5.71 | 5.54 | 0.11 | 0.33 | 0.03 | 0.45 | 0.13 | 0.65 | 0.14 | 0.26 |
| ELANE | multiparous | 5.01 | 4.80 | 4.82 | 5.11 | 0.11 | 5.11 | 4.83 | 4.80 | 5.11 | 0.10 | 5.23 | 4.76 | 4.78 | 5.16 | 0.10 | | | | | | | |
| | primiparous | 5.06 | 4.70 | 4.73 | 5.07 | 0.13 | 5.16 | 4.84 | 4.86 | 5.17 | 0.13 | 5.09 | 4.94 | 4.90 | 5.30 | 0.14 | 0.52 | 0.00 | 0.99 | 0.76 | 0.80 | 0.93 | 0.55 |
| CTSB | multiparous | 6.83 | 6.04 | 6.71 | 6.83 | 0.11 | 6.65 | 6.03 | 6.64 | 6.86 | 0.09 | 6.62 | 6.21 | 6.69 | 6.92 | 0.09 | | | | | | | |
| | primiparous | 6.76 | 6.32 | 6.68 | 6.88 | 0.12 | 6.68 | 6.37 | 6.43 | 6.92 | 0.12 | 6.79 | 6.17 | 6.58 | 6.82 | 0.14 | 0.68 | 0.00 | 0.82 | 0.60 | 0.81 | 0.08 | 0.43 |
| ITGA4 | multiparous | 6.99 | 6.92 | 7.08 | 7.14 | 0.09 | 7.25 | 7.10 | 7.18 | 7.11 | 0.08 | 7.29 | 6.87 | 7.12 | 7.04 | 0.08 | | | | | | | |
| | primiparous | 7.17 | 7.15 | 7.21 | 7.16 | 0.10 | 7.14 | 6.89 | 7.41 | 7.10 | 0.10 | 7.07 | 6.90 | 7.13 | 7.05 | 0.12 | 0.34 | 0.00 | 0.35 | 0.64 | 0.27 | 0.40 | 0.26 |
| ACOX2 | multiparous | 4.65 | 4.42 | 4.58 | 4.67 | 0.14 | 4.77 | 4.55 | 4.57 | 4.68 | 0.12 | 4.94 | 4.54 | 4.56 | 4.65 | 0.12 | | | | | | | |
| | primiparous | 4.65 | 4.49 | 4.63 | 4.61 | 0.15 | 4.66 | 4.31 | 4.62 | 4.66 | 0.15 | 4.66 | 4.11 | 4.55 | 4.51 | 0.17 | 0.91 | 0.00 | 0.81 | 0.24 | 0.47 | 0.45 | 0.91 |
| MMP9 | multiparous | 5.22 | 6.25 | 5.59 | 5.40 | 0.12 | 5.16 | 6.10 | 5.46 | 5.23 | 0.10 | 5.27 | 6.31 | 5.50 | 5.38 | 0.10 | | | | | | | |
| | primiparous | 5.31 | 5.95 | 5.48 | 5.39 | 0.13 | 5.11 | 6.06 | 5.52 | 5.21 | 0.13 | 5.16 | 6.23 | 5.48 | 5.13 | 0.14 | 0.47 | 0.00 | 0.49 | 0.32 | 0.81 | 0.67 | 0.65 |
| CXCL2 | multiparous | 4.62 | 4.93 | 4.78 | 4.79 | 0.13 | 4.85 | 5.10 | 4.95 | 4.82 | 0.11 | 5.13 | 4.99 | 4.91 | 4.74 | 0.11 | | | | | | | |
| | primiparous | 4.72 | 5.12 | 4.84 | 4.72 | 0.14 | 4.85 | 4.93 | 5.06 | 4.67 | 0.14 | 5.09 | 5.22 | 4.85 | 4.73 | 0.16 | 0.40 | 0.00 | 0.02 | 0.86 | 0.82 | 0.48 | 0.47 |
| IL10 | multiparous | 4.24 | 4.52 | 4.12 | 4.45 | 0.11 | 4.37 | 4.58 | 4.25 | 4.38 | 0.10 | 4.36 | 4.52 | 4.18 | 4.46 | 0.10 | | | | | | | |
| | primiparous | 4.18 | 4.51 | 4.10 | 4.11 | 0.13 | 4.40 | 4.64 | 4.44 | 4.45 | 0.13 | 4.41 | 4.60 | 4.31 | 4.47 | 0.14 | 0.13 | 0.00 | 0.88 | 0.82 | 0.44 | 0.43 | 0.91 |
| SIRPA | multiparous | 6.87 | 6.68 | 6.84 | 6.92 | 0.06 | 6.93 | 6.81 | 6.94 | 6.97 | 0.05 | 6.84 | 6.71 | 6.93 | 6.97 | 0.05 | | | | | | | |
| | primiparous | 6.87 | 6.79 | 6.92 | 6.90 | 0.07 | 6.86 | 6.65 | 6.99 | 6.95 | 0.07 | 6.93 | 6.79 | 6.92 | 6.91 | 0.07 | 0.69 | 0.00 | 0.93 | 0.89 | 0.54 | 0.73 | 0.23 |
| SOCS2 | multiparous | 5.44 | 5.47 | 5.38 | 5.57 | 0.09 | 5.67 | 5.67 | 5.51 | 5.47 | 0.08 | 5.64 | 5.43 | 5.53 | 5.51 | 0.08 | | | | | | | |
| | primiparous | 5.44 | 5.65 | 5.58 | 5.50 | 0.11 | 5.46 | 5.30 | 5.58 | 5.43 | 0.10 | 5.38 | 5.34 | 5.42 | 5.40 | 0.12 | 0.60 | 0.93 | 0.45 | 0.19 | 0.13 | 0.25 | 0.32 |
| IKZF1 | multiparous | 6.44 | 6.41 | 6.35 | 6.43 | 0.03 | 6.45 | 6.48 | 6.37 | 6.41 | 0.03 | 6.51 | 6.51 | 6.36 | 6.46 | 0.03 | | | | | | | |
| | primiparous | 6.43 | 6.49 | 6.39 | 6.45 | 0.04 | 6.43 | 6.44 | 6.32 | 6.41 | 0.04 | 6.40 | 6.50 | 6.34 | 6.47 | 0.04 | 0.24 | 0.00 | 0.74 | 0.54 | 0.14 | 0.43 | 0.71 |
| RORa | multiparous | 5.96 | 5.90 | 5.92 | 6.10 | 0.08 | 6.14 | 6.00 | 5.99 | 5.99 | 0.07 | 6.21 | 5.92 | 6.00 | 6.00 | 0.07 | | | | | | | |
| | primiparous | 6.00 | 6.04 | 6.02 | 6.00 | 0.09 | 6.01 | 5.91 | 6.05 | 5.94 | 0.09 | 5.91 | 5.90 | 5.89 | 5.94 | 0.10 | 0.77 | 0.22 | 0.64 | 0.32 | 0.23 | 0.32 | 0.60 |
| FABP3 | multiparous | 5.77 | 5.85 | 5.85 | 5.96 | 0.07 | 6.04 | 5.91 | 6.04 | 5.97 | 0.06 | 6.13 | 5.75 | 5.94 | 5.88 | 0.06 | | | | | | | |
| | primiparous | 6.10 | 6.04 | 6.08 | 6.07 | 0.07 | 6.13 | 6.01 | 6.17 | 6.04 | 0.07 | 6.07 | 6.00 | 6.14 | 6.07 | 0.08 | 0.33 | 0.00 | 0.08 | 0.00 | 0.44 | 0.61 | 0.12 |
| CCL2 | multiparous | 2.97 | 4.39 | 3.33 | 3.28 | 0.35 | 3.25 | 4.61 | 3.92 | 3.79 | 0.30 | 4.03 | 4.34 | 3.47 | 3.34 | 0.30 | | | | | | | |
| | primiparous | 3.14 | 4.36 | 4.11 | 3.90 | 0.39 | 3.00 | 3.95 | 3.74 | 3.21 | 0.38 | 3.37 | 4.48 | 4.00 | 3.25 | 0.43 | 0.82 | 0.00 | 0.63 | 0.92 | 0.09 | 0.42 | 0.86 |
| STAT3 | multiparous | 6.66 | 6.82 | 6.64 | 6.66 | 0.09 | 6.64 | 6.83 | 6.66 | 6.70 | 0.08 | 6.76 | 6.84 | 6.67 | 6.71 | 0.08 | | | | | | | |
| | primiparous | 6.70 | 6.83 | 6.66 | 6.69 | 0.10 | 6.66 | 6.27 | 6.62 | 6.69 | 0.10 | 6.69 | 6.93 | 6.61 | 6.64 | 0.12 | 0.18 | 0.23 | 0.09 | 0.29 | 0.27 | 0.46 | 0.07 |
| ADGRE1 | multiparous | 5.90 | 6.37 | 6.02 | 5.85 | 0.12 | 6.09 | 6.40 | 6.10 | 5.96 | 0.10 | 6.00 | 6.45 | 6.13 | 6.01 | 0.10 | | | | | | | |
| | primiparous | 6.02 | 6.37 | 6.03 | 5.90 | 0.13 | 5.84 | 5.85 | 6.04 | 5.88 | 0.13 | 5.92 | 6.56 | 6.14 | 5.76 | 0.15 | 0.43 | 0.00 | 0.13 | 0.25 | 0.21 | 0.76 | 0.21 |
| BIRC5 | multiparous | 5.24 | 5.39 | 5.33 | 5.20 | 0.11 | 5.41 | 5.45 | 5.40 | 5.34 | 0.09 | 5.60 | 5.29 | 5.35 | 5.28 | 0.09 | | | | | | | |
| | primiparous | 5.45 | 5.48 | 5.44 | 5.40 | 0.12 | 5.47 | 5.06 | 5.54 | 5.46 | 0.12 | 5.46 | 5.40 | 5.48 | 5.55 | 0.13 | 0.67 | 0.35 | 0.32 | 0.20 | 0.44 | 0.16 | 0.21 |
| DEFB5 | multiparous | 4.65 | 5.10 | 5.22 | 5.24 | 0.37 | 4.74 | 4.96 | 5.50 | 4.96 | 0.32 | 5.28 | 4.95 | 5.12 | 4.84 | 0.32 | | | | | | | |
| | primiparous | 4.92 | 4.62 | 4.75 | 4.75 | 0.42 | 4.59 | 4.54 | 5.44 | 4.90 | 0.41 | 4.40 | 4.57 | 4.36 | 4.45 | 0.46 | 0.78 | 0.19 | 0.22 | 0.20 | 0.78 | 0.92 | 0.59 |
| CXCL10 | multiparous | 5.97 | 6.44 | 5.82 | 6.15 | 0.13 | 6.12 | 6.43 | 6.01 | 6.03 | 0.12 | 6.28 | 6.34 | 5.99 | 6.01 | 0.11 | | | | | | | |
| | primiparous | 5.86 | 6.38 | 5.90 | 5.90 | 0.15 | 6.02 | 5.90 | 6.07 | 5.89 | 0.15 | 5.97 | 6.25 | 5.78 | 5.88 | 0.16 | 0.99 | 0.00 | 0.24 | 0.02 | 0.73 | 0.61 | 0.50 |
| NFkB2 | multiparous | 4.44 | 4.67 | 4.25 | 4.62 | 0.30 | 4.44 | 4.61 | 4.74 | 4.81 | 0.26 | 4.74 | 5.13 | 4.96 | 4.82 | 0.25 | | | | | | | |
| | primiparous | 5.14 | 4.61 | 4.77 | 4.75 | 0.33 | 5.11 | 4.80 | 4.47 | 5.18 | 0.33 | 5.13 | 5.17 | 5.00 | 4.47 | 0.37 | 0.40 | 0.81 | 0.45 | 0.25 | 0.75 | 0.26 | 0.76 |
| IL6 | multiparous | 3.75 | 3.53 | 3.66 | 3.91 | 0.22 | 3.84 | 4.12 | 3.88 | 3.38 | 0.19 | 4.18 | 3.32 | 3.59 | 3.79 | 0.19 | | | | | | | |
| | primiparous | 3.58 | 4.26 | 3.52 | 3.78 | 0.25 | 3.66 | 3.54 | 3.80 | 3.92 | 0.25 | 3.63 | 3.61 | 3.62 | 4.09 | 0.27 | 0.96 | 0.74 | 0.17 | 0.96 | 0.85 | 0.14 | 0.05 |
| TAP | multiparous | 2.23 | 1.86 | 1.32 | 2.98 | 0.57 | 3.15 | 2.40 | 2.23 | 2.20 | 0.50 | 3.18 | 1.77 | 2.15 | 2.06 | 0.49 | | | | | | | |
| | primiparous | 1.99 | 2.36 | 2.20 | 2.42 | 0.64 | 2.57 | 1.92 | 1.97 | 1.94 | 0.63 | 3.72 | 2.35 | 2.34 | 2.66 | 0.71 | 0.55 | 0.09 | 0.46 | 0.78 | 0.39 | 0.92 | 0.94 |
| **cell function gene expression** | | | | | | | | | | | | | | | | | | | | | | | |
| SCD | multiparous | 5.70 | 5.82 | 5.78 | 5.90 | 0.10 | 5.84 | 5.79 | 5.81 | 5.68 | 0.09 | 6.05 | 5.71 | 5.83 | 5.69 | 0.09 | | | | | | | |
| | primiparous | 5.64 | 5.88 | 5.81 | 5.75 | 0.12 | 5.69 | 5.67 | 5.74 | 5.62 | 0.12 | 5.51 | 5.57 | 5.58 | 5.61 | 0.13 | 0.36 | 0.87 | 0.34 | 0.03 | 0.22 | 0.42 | 0.62 |
| LPL | multiparous | 5.49 | 4.73 | 5.44 | 5.67 | 0.12 | 5.57 | 4.82 | 5.52 | 5.68 | 0.11 | 5.33 | 4.88 | 5.49 | 5.71 | 0.11 | | | | | | | |
| | primiparous | 5.56 | 5.09 | 5.53 | 5.67 | 0.14 | 5.57 | 4.98 | 5.55 | 5.70 | 0.14 | 5.66 | 4.86 | 5.48 | 5.60 | 0.15 | 0.83 | 0.00 | 1.00 | 0.29 | 0.86 | 0.43 | 0.58 |
| **Reference gene expression** | | | | | | | | | | | | | | | | | | | | | | | |
| GAPDH | multiparous | 7.10 | 6.94 | 7.14 | 6.94 | 0.12 | 6.88 | 6.84 | 7.08 | 7.11 | 0.11 | 6.90 | 7.15 | 7.22 | 7.20 | 0.11 | | | | | | | |
| | primiparous | 7.19 | 6.96 | 7.11 | 7.16 | 0.14 | 7.17 | 7.19 | 7.07 | 7.22 | 0.14 | 7.26 | 7.29 | 7.37 | 7.23 | 0.15 | 0.09 | 0.47 | 0.54 | 0.02 | 0.68 | 0.51 | 0.76 |
| PPIA | multiparous | 7.40 | 7.44 | 7.35 | 7.54 | 0.09 | 7.62 | 7.58 | 7.53 | 7.49 | 0.07 | 7.72 | 7.38 | 7.46 | 7.44 | 0.07 | | | | | | | |
| | primiparous | 7.56 | 7.64 | 7.57 | 7.53 | 0.10 | 7.61 | 7.45 | 7.68 | 7.50 | 0.09 | 7.52 | 7.47 | 7.48 | 7.51 | 0.11 | 0.47 | 0.30 | 0.22 | 0.34 | 0.39 | 0.45 | 0.32 |
| RPL19 | multiparous | 7.91 | 7.91 | 7.87 | 8.07 | 0.08 | 8.09 | 8.05 | 8.05 | 7.99 | 0.07 | 8.19 | 7.86 | 7.98 | 7.97 | 0.07 | | | | | | | |
| | primiparous | 8.01 | 8.10 | 8.04 | 8.01 | 0.09 | 8.05 | 7.86 | 8.17 | 7.96 | 0.09 | 7.97 | 7.93 | 7.95 | 7.97 | 0.10 | 0.63 | 0.27 | 0.18 | 0.90 | 0.35 | 0.42 | 0.22 |
| reference genes [3] | multiparous | 9.90 | 9.70 | 9.99 | 9.74 | 0.16 | 9.73 | 9.53 | 9.87 | 9.94 | 0.14 | 9.39 | 9.75 | 9.98 | 9.98 | 0.14 | | | | | | | |
| | primiparous | 10.10 | 9.53 | 9.92 | 10.09 | 0.18 | 10.04 | 9.96 | 9.84 | 10.09 | 0.18 | 10.19 | 9.93 | 10.19 | 10.06 | 0.20 | 0.70 | 0.04 | 0.55 | 0.01 | 0.40 | 0.22 | 0.40 |

1 Blood samples were collected before LPS challenge (-20 h) and after LPS challenge times (4 h, 28 h and 76 h).

2 Probability: TREAT = effect of treatment (control vs vitamin E vs plant extract group); TIMES = effect of times before (-20 hours) vs after LPS challenge (vs 4 vs 28 vs 76 hours); TREAT × TIMES = the interaction between treatment and times; PAR = effect of parity (primiparous vs multiparous); PAR × TIMES = interaction between parity and times; TREAT ×

3 Decimal logarithm of the geometric mean of ACTB (actin beta), YWHAZ (tyrosine 3-monooxygenase/tryptophan 5-monooxygenase activation protein zeta), and RPLP0 (Ribosomal Protein Lateral Stalk Subunit P0) genes.

Table S3: Abundance of antioxidant function mRNA, immune function mRNA, milk synthesis mRNA and gene cell death mRNA determined by real-time quantitative RT-PCR in milk in control unsupplemented group (n = 11), vitamin E supplemented group (n = 13), and plant extracts supplemented group (n = 12) dairy cows before (0 h)and after LPS challenge times (4 h, 28 h and 76 h).

| Item[1] | Parity | control | | | | | vitamin E | | | | | Plant extracts | | | | | *P*-value[2] | | | | | | |
|---|---|---|---|---|---|---|---|---|---|---|---|---|---|---|---|---|---|---|---|---|---|---|---|
| | | 0 | 4 | 28 | 76 | SEM | 0 | 4 | 28 | 76 | SEM | 0 | 4 | 28 | 76 | SEM | TREAT | TIMES | TREAT x TIMES | PAR | TREAT x PAR | PAR x TIMES | TREAT x PAR x TIMES |
| **Redox gene expression** | | | | | | | | | | | | | | | | | | | | | | | |
| CAT | multiparous | 6.97 | 6.52 | 6.75 | 6.87 | 0.07 | 7.02 | 6.33 | 6.79 | 6.89 | 0.06 | 6.95 | 6.40 | 6.70 | 6.89 | 0.06 | | | | | | | |
| | primiparous | 6.78 | 6.19 | 6.72 | 6.95 | 0.08 | 7.03 | 6.53 | 6.67 | 6.94 | 0.08 | 6.92 | 6.35 | 6.74 | 6.87 | 0.08 | 0.33 | 0.00 | 0.71 | 0.36 | 0.20 | 0.52 | 0.02 |
| GPX1 | multiparous | 6.77 | 6.94 | 7.55 | 7.16 | 0.08 | 6.84 | 7.09 | 7.58 | 7.14 | 0.07 | 6.91 | 7.31 | 7.52 | 7.23 | 0.07 | | | | | | | |
| | primiparous | 7.05 | 7.13 | 7.54 | 7.43 | 0.09 | 7.00 | 7.13 | 7.47 | 7.26 | 0.09 | 7.10 | 7.17 | 7.56 | 7.33 | 0.10 | 0.19 | 0.00 | 0.48 | 0.03 | 0.28 | 0.04 | 0.74 |
| GPX3 | multiparous | 4.24 | 6.10 | 5.95 | 5.29 | 0.14 | 4.33 | 6.27 | 5.79 | 5.12 | 0.12 | 4.34 | 6.36 | 5.80 | 5.06 | 0.12 | | | | | | | |
| | primiparous | 5.22 | 6.18 | 5.78 | 5.28 | 0.16 | 4.71 | 6.02 | 5.80 | 5.10 | 0.16 | 5.24 | 6.10 | 5.81 | 5.35 | 0.18 | 0.30 | 0.00 | 0.79 | 0.04 | 0.40 | 0.00 | 0.35 |
| LAP | multiparous | 4.51 | 6.06 | 5.79 | 5.25 | 0.67 | 4.75 | 6.59 | 5.35 | 5.46 | 0.58 | 4.93 | 6.68 | 4.45 | 4.44 | 0.58 | | | | | | | |
| | primiparous | 4.69 | 6.15 | 5.89 | 3.79 | 0.75 | 4.46 | 6.22 | 4.84 | 4.46 | 0.75 | 4.92 | 6.20 | 6.96 | 6.21 | 0.83 | 0.66 | 0.00 | 0.88 | 0.90 | 0.18 | 0.54 | 0.29 |
| NFE2L2 | multiparous | 4.49 | 4.99 | 5.77 | 4.93 | 0.13 | 4.64 | 4.86 | 5.71 | 4.86 | 0.11 | 4.48 | 4.95 | 5.65 | 4.96 | 0.11 | | | | | | | |
| | primiparous | 4.98 | 5.09 | 5.60 | 5.18 | 0.15 | 4.76 | 4.96 | 5.80 | 4.95 | 0.15 | 4.86 | 5.14 | 5.79 | 5.07 | 0.16 | 0.73 | 0.00 | 0.86 | 0.03 | 0.79 | 0.21 | 0.69 |
| SOD1 | multiparous | 7.84 | 7.29 | 7.31 | 7.27 | 0.09 | 7.73 | 7.08 | 7.27 | 7.16 | 0.08 | 7.71 | 7.32 | 7.16 | 7.28 | 0.08 | | | | | | | |
| | primiparous | 7.68 | 7.04 | 7.28 | 7.36 | 0.10 | 7.72 | 7.34 | 7.30 | 7.34 | 0.10 | 7.68 | 7.16 | 7.25 | 7.25 | 0.12 | 0.85 | 0.00 | 0.88 | 0.95 | 0.19 | 0.48 | 0.36 |
| SOD2 | multiparous | 7.12 | 8.09 | 7.89 | 6.99 | 0.09 | 7.06 | 8.23 | 7.86 | 6.85 | 0.08 | 7.05 | 8.27 | 7.80 | 7.03 | 0.08 | | | | | | | |
| | primiparous | 7.34 | 8.15 | 7.79 | 7.08 | 0.11 | 7.11 | 8.01 | 7.88 | 6.92 | 0.10 | 7.32 | 8.27 | 7.87 | 7.03 | 0.12 | 0.29 | 0.00 | 0.44 | 0.40 | 0.62 | 0.14 | 0.68 |
| SOD3 | multiparous | 3.67 | 3.47 | 2.66 | 3.08 | 0.29 | 3.44 | 3.09 | 2.56 | 2.81 | 0.25 | 3.31 | 3.13 | 2.99 | 3.09 | 0.25 | | | | | | | |
| | primiparous | 3.64 | 3.05 | 2.72 | 3.12 | 0.32 | 3.61 | 3.31 | 3.81 | 3.27 | 0.32 | 3.53 | 3.42 | 2.57 | 2.60 | 0.36 | 0.57 | 0.00 | 0.74 | 0.39 | 0.06 | 0.82 | 0.32 |
| TXNRD1 | multiparous | 6.76 | 6.70 | 7.17 | 7.03 | 0.07 | 6.89 | 6.75 | 7.17 | 7.09 | 0.06 | 6.82 | 6.83 | 7.09 | 7.12 | 0.06 | | | | | | | |
| | primiparous | 6.85 | 6.61 | 7.05 | 7.20 | 0.08 | 6.86 | 6.70 | 7.08 | 7.05 | 0.07 | 6.97 | 6.70 | 7.03 | 7.06 | 0.08 | 0.68 | 0.00 | 0.58 | 0.48 | 0.65 | 0.12 | 0.57 |
| AHR | multiparous | 5.55 | 5.61 | 5.79 | 5.63 | 0.09 | 5.60 | 5.63 | 5.80 | 5.68 | 0.08 | 5.52 | 5.63 | 5.74 | 5.74 | 0.08 | | | | | | | |
| | primiparous | 5.48 | 5.56 | 5.63 | 5.78 | 0.10 | 5.38 | 5.41 | 5.64 | 5.63 | 0.10 | 5.45 | 5.70 | 5.73 | 5.73 | 0.11 | 0.53 | 0.00 | 0.90 | 0.17 | 0.28 | 0.38 | 0.87 |
| ACOX2 | multiparous | 3.72 | 7.08 | 6.79 | 5.90 | 0.26 | 4.23 | 7.27 | 6.71 | 5.89 | 0.23 | 4.12 | 7.49 | 6.68 | 5.97 | 0.22 | | | | | | | |
| | primiparous | 5.14 | 7.11 | 6.48 | 5.94 | 0.29 | 4.51 | 7.05 | 6.67 | 5.86 | 0.29 | 5.04 | 7.04 | 6.49 | 5.94 | 0.32 | 0.85 | 0.00 | 0.99 | 0.38 | 0.58 | 0.00 | 0.58 |
| AOAH | multiparous | 3.57 | 6.56 | 6.42 | 5.07 | 0.21 | 3.83 | 6.67 | 6.33 | 5.45 | 0.18 | 3.79 | 6.75 | 6.27 | 5.53 | 0.18 | | | | | | | |
| | primiparous | 4.58 | 6.57 | 6.15 | 5.39 | 0.23 | 4.15 | 6.49 | 6.24 | 5.36 | 0.23 | 4.93 | 6.66 | 6.37 | 5.59 | 0.26 | 0.20 | 0.00 | 0.89 | 0.07 | 0.32 | 0.00 | 0.73 |
| CCL2 | multiparous | 5.00 | 6.69 | 6.61 | 6.41 | 0.59 | 4.64 | 6.14 | 7.52 | 6.44 | 0.51 | 5.25 | 6.55 | 6.77 | 6.56 | 0.51 | | | | | | | |
| | primiparous | 5.81 | 6.81 | 6.94 | 6.60 | 0.65 | 5.19 | 6.94 | 7.53 | 6.67 | 0.65 | 5.38 | 6.90 | 7.39 | 5.59 | 0.72 | 0.96 | 0.00 | 0.78 | 0.33 | 0.81 | 0.76 | 0.94 |
| CD36 | multiparous | 7.96 | 7.69 | 8.33 | 8.17 | 0.08 | 8.06 | 7.58 | 8.34 | 8.10 | 0.07 | 7.97 | 7.74 | 8.25 | 8.10 | 0.07 | | | | | | | |
| | primiparous | 8.03 | 7.52 | 8.22 | 8.23 | 0.09 | 8.08 | 7.93 | 8.23 | 8.23 | 0.09 | 8.19 | 7.82 | 8.27 | 8.16 | 0.10 | 0.49 | 0.00 | 0.44 | 0.21 | 0.29 | 0.22 | 0.13 |
| CXCL8 | multiparous | 6.13 | 8.84 | 8.57 | 7.86 | 0.21 | 6.40 | 9.00 | 8.51 | 7.74 | 0.18 | 6.55 | 9.21 | 8.53 | 7.93 | 0.18 | | | | | | | |
| | primiparous | 6.93 | 9.00 | 8.57 | 7.95 | 0.24 | 6.56 | 8.88 | 8.71 | 7.88 | 0.23 | 7.10 | 9.15 | 8.52 | 7.92 | 0.26 | 0.35 | 0.00 | 0.82 | 0.11 | 0.73 | 0.17 | 0.87 |
| CLEC4A | multiparous | 4.87 | 6.12 | 7.47 | 6.86 | 0.19 | 5.01 | 6.30 | 7.30 | 6.80 | 0.17 | 5.19 | 6.30 | 7.18 | 6.91 | 0.16 | | | | | | | |
| | primiparous | 5.79 | 6.22 | 7.14 | 6.99 | 0.21 | 5.40 | 6.13 | 7.36 | 6.89 | 0.21 | 5.84 | 6.37 | 7.33 | 7.00 | 0.24 | 0.50 | 0.00 | 0.89 | 0.05 | 0.74 | 0.01 | 0.74 |
| DHX58 | multiparous | 5.40 | 6.45 | 6.06 | 5.59 | 0.15 | 5.15 | 6.81 | 6.01 | 5.66 | 0.13 | 5.37 | 6.82 | 6.08 | 5.72 | 0.13 | | | | | | | |
| | primiparous | 5.31 | 6.82 | 6.02 | 5.75 | 0.17 | 5.21 | 6.69 | 6.02 | 5.59 | 0.16 | 5.40 | 6.91 | 6.20 | 5.81 | 0.18 | 0.25 | 0.00 | 0.88 | 0.52 | 0.73 | 0.90 | 0.75 |
| TNF | multiparous | 3.57 | 6.12 | 5.31 | 4.81 | 0.23 | 3.55 | 6.42 | 5.03 | 4.56 | 0.20 | 3.95 | 6.43 | 5.14 | 4.68 | 0.20 | | | | | | | |
| | primiparous | 3.98 | 6.29 | 4.94 | 4.16 | 0.25 | 3.67 | 6.20 | 4.95 | 4.57 | 0.25 | 4.35 | 6.28 | 5.02 | 4.85 | 0.28 | 0.14 | 0.00 | 0.71 | 0.80 | 0.76 | 0.23 | 0.60 |
| MUC1 | multiparous | 5.97 | 5.82 | 5.11 | 5.61 | 0.14 | 6.10 | 5.60 | 4.79 | 5.34 | 0.12 | 6.07 | 5.85 | 5.16 | 5.50 | 0.12 | | | | | | | |
| | primiparous | 6.15 | 5.82 | 5.40 | 5.72 | 0.16 | 6.26 | 6.15 | 5.10 | 5.72 | 0.15 | 6.16 | 5.92 | 4.95 | 5.66 | 0.17 | 0.72 | 0.00 | 0.39 | 0.02 | 0.13 | 0.93 | 0.58 |
| TLR2 | multiparous | 5.30 | 6.84 | 6.97 | 6.42 | 0.11 | 5.44 | 6.95 | 6.95 | 6.34 | 0.09 | 5.47 | 7.06 | 6.93 | 6.42 | 0.09 | | | | | | | |
| | primiparous | 5.71 | 6.88 | 6.85 | 6.47 | 0.12 | 5.57 | 6.81 | 6.90 | 6.38 | 0.12 | 5.66 | 6.94 | 6.92 | 6.48 | 0.13 | 0.46 | 0.00 | 0.96 | 0.42 | 0.64 | 0.05 | 0.91 |
| STAT2 | multiparous | 4.79 | 4.71 | 4.78 | 4.42 | 0.08 | 4.59 | 4.85 | 4.71 | 4.29 | 0.07 | 4.64 | 4.91 | 4.70 | 4.49 | 0.07 | | | | | | | |
| | primiparous | 4.89 | 4.72 | 4.56 | 4.43 | 0.09 | 4.74 | 4.73 | 4.55 | 4.25 | 0.09 | 4.73 | 4.87 | 4.70 | 4.38 | 0.10 | 0.09 | 0.00 | 0.18 | 0.48 | 0.93 | 0.13 | 0.84 |
| LCN2 | multiparous | 6.07 | 6.03 | 5.91 | 6.06 | 0.12 | 6.20 | 6.20 | 5.76 | 5.86 | 0.11 | 6.18 | 6.34 | 5.84 | 6.01 | 0.11 | | | | | | | |
| | primiparous | 6.47 | 6.35 | 6.13 | 6.40 | 0.14 | 6.50 | 6.42 | 6.12 | 6.33 | 0.14 | 6.61 | 6.58 | 6.09 | 6.53 | 0.15 | 0.47 | 0.00 | 0.34 | 0.00 | 0.97 | 0.41 | 0.93 |
| PLAU | multiparous | 5.16 | 8.34 | 7.46 | 6.78 | 0.20 | 5.37 | 8.37 | 7.33 | 6.51 | 0.17 | 5.70 | 8.50 | 7.41 | 6.69 | 0.17 | | | | | | | |
| | primiparous | 6.02 | 8.15 | 7.30 | 6.71 | 0.22 | 5.88 | 8.01 | 7.44 | 6.57 | 0.22 | 6.03 | 8.26 | 7.39 | 6.63 | 0.25 | 0.45 | 0.00 | 0.91 | 0.53 | 0.89 | 0.00 | 0.90 |
| IL1A | multiparous | 4.23 | 7.95 | 6.40 | 6.02 | 0.20 | 4.63 | 8.08 | 6.23 | 5.59 | 0.18 | 4.63 | 8.28 | 6.20 | 5.70 | 0.18 | | | | | | | |
| | primiparous | 5.30 | 8.09 | 6.09 | 5.64 | 0.23 | 4.83 | 7.88 | 6.26 | 5.48 | 0.23 | 5.23 | 8.18 | 6.21 | 5.84 | 0.25 | 0.37 | 0.00 | 0.85 | 0.36 | 0.68 | 0.01 | 0.36 |
| S100A7 | multiparous | 4.17 | 4.56 | 4.72 | 3.58 | 0.18 | 4.23 | 4.68 | 4.65 | 3.35 | 0.16 | 4.17 | 4.72 | 4.43 | 3.43 | 0.16 | | | | | | | |
| | primiparous | 4.58 | 4.96 | 4.52 | 3.83 | 0.21 | 4.29 | 4.44 | 4.55 | 3.09 | 0.20 | 4.31 | 4.55 | 4.73 | 3.23 | 0.23 | 0.14 | 0.00 | 0.70 | 0.72 | 0.28 | 0.59 | 0.53 |
| TLR4 | multiparous | 5.29 | 6.89 | 6.64 | 5.95 | 0.14 | 5.35 | 7.02 | 6.48 | 5.82 | 0.12 | 5.21 | 7.21 | 6.55 | 5.99 | 0.12 | | | | | | | |
| | primiparous | 5.77 | 7.18 | 6.54 | 6.13 | 0.16 | 5.45 | 6.98 | 6.54 | 5.89 | 0.16 | 5.63 | 7.13 | 6.68 | 6.00 | 0.17 | 0.33 | 0.00 | 0.92 | 0.09 | 0.61 | 0.22 | 0.71 |
| STAT1 | multiparous | 5.05 | 6.96 | 7.05 | 6.42 | 0.18 | 5.45 | 7.06 | 7.06 | 6.35 | 0.15 | 5.63 | 7.24 | 6.92 | 6.44 | 0.15 | | | | | | | |
| | primiparous | 5.85 | 7.19 | 6.95 | 6.51 | 0.20 | 4.99 | 6.94 | 6.97 | 6.36 | 0.20 | 6.22 | 7.13 | 7.00 | 6.48 | 0.22 | 0.06 | 0.00 | 0.11 | 0.34 | 0.08 | 0.34 | 0.13 |
| CD14 | multiparous | 5.10 | 7.18 | 7.41 | 7.19 | 0.15 | 5.80 | 7.31 | 7.36 | 7.16 | 0.13 | 5.94 | 7.21 | 7.44 | 7.21 | 0.13 | | | | | | | |
| | primiparous | 6.50 | 7.34 | 7.37 | 7.34 | 0.17 | 6.28 | 7.19 | 7.24 | 7.22 | 0.17 | 6.33 | 7.32 | 7.44 | 7.32 | 0.19 | 0.43 | 0.00 | 0.64 | 0.00 | 0.09 | 0.00 | 0.23 |
| CXCL1 | multiparous | 5.37 | 7.92 | 7.84 | 7.02 | 0.24 | 5.65 | 8.10 | 7.83 | 6.70 | 0.21 | 5.51 | 8.32 | 7.87 | 6.90 | 0.21 | | | | | | | |
| | primiparous | 6.66 | 8.09 | 7.73 | 6.92 | 0.27 | 5.62 | 7.88 | 7.96 | 6.68 | 0.27 | 6.53 | 8.00 | 7.86 | 6.86 | 0.30 | 0.34 | 0.00 | 0.80 | 0.20 | 0.39 | 0.01 | 0.28 |
| IL2RA | multiparous | 4.60 | 5.04 | 4.05 | 3.90 | 0.27 | 4.54 | 4.85 | 4.17 | 3.92 | 0.24 | 4.42 | 5.34 | 4.27 | 4.05 | 0.24 | | | | | | | |
| | primiparous | 4.63 | 5.59 | 5.14 | 4.15 | 0.30 | 4.38 | 5.74 | 5.48 | 3.89 | 0.30 | 4.47 | 5.82 | 5.50 | 4.22 | 0.34 | 0.62 | 0.00 | 0.86 | 0.00 | 1.00 | 0.00 | 0.97 |
| IFNB1 | multiparous | 4.10 | 4.28 | 5.21 | 4.82 | 0.16 | 4.32 | 4.62 | 5.15 | 4.95 | 0.14 | 4.01 | 4.84 | 5.06 | 5.09 | 0.14 | | | | | | | |
| | primiparous | 4.69 | 4.67 | 5.04 | 4.84 | 0.18 | 4.35 | 4.62 | 5.35 | 5.06 | 0.18 | 4.72 | 4.65 | 5.24 | 5.19 | 0.20 | 0.25 | 0.00 | 0.73 | 0.03 | 0.70 | 0.15 | 0.20 |
| IL1B | multiparous | 4.30 | 8.67 | 7.01 | 6.17 | 0.30 | 4.63 | 8.80 | 6.96 | 5.95 | 0.26 | 4.47 | 8.94 | 6.97 | 6.15 | 0.26 | | | | | | | |
| | primiparous | 4.83 | 8.80 | 6.82 | 6.08 | 0.33 | 4.64 | 8.71 | 6.99 | 5.98 | 0.33 | 5.51 | 8.93 | 7.03 | 6.18 | 0.37 | 0.39 | 0.00 | 0.97 | 0.37 | 0.65 | 0.34 | 0.89 |
| S100A8 | multiparous | 5.17 | 8.18 | 8.27 | 6.95 | 0.28 | 5.85 | 8.33 | 8.36 | 7.09 | 0.24 | 5.78 | 8.64 | 8.21 | 7.13 | 0.24 | | | | | | | |
| | primiparous | 5.83 | 8.30 | 8.24 | 7.11 | 0.31 | 5.21 | 8.22 | 8.47 | 7.11 | 0.31 | 5.89 | 8.44 | 8.38 | 7.16 | 0.34 | 0.41 | 0.00 | 0.96 | 0.80 | 0.42 | 0.97 | 0.67 |
| STAT3 | multiparous | 7.11 | 7.34 | 7.09 | 6.74 | 0.07 | 7.16 | 7.48 | 7.03 | 6.55 | 0.06 | 7.13 | 7.47 | 7.04 | 6.69 | 0.06 | | | | | | | |
| | primiparous | 6.93 | 7.35 | 6.97 | 6.75 | 0.08 | 7.10 | 7.34 | 6.98 | 6.68 | 0.08 | 7.06 | 7.38 | 7.00 | 6.70 | 0.09 | 0.86 | 0.00 | 0.35 | 0.21 | 0.89 | 0.26 | 0.82 |
| IL6 | multiparous | 1.02 | 6.74 | 3.81 | 2.25 | 0.43 | 0.98 | 6.95 | 4.25 | 1.80 | 0.37 | 1.68 | 7.02 | 4.16 | 1.37 | 0.37 | | | | | | | |
| | primiparous | 1.87 | 6.86 | 4.19 | 1.67 | 0.48 | 1.02 | 6.52 | 4.75 | 1.69 | 0.48 | 2.03 | 6.86 | 4.19 | 0.97 | 0.53 | 0.97 | 0.00 | 0.21 | 0.80 | 0.86 | 0.39 | 0.96 |
| S100A9 | multiparous | 5.11 | 8.30 | 8.25 | 6.84 | 0.28 | 5.71 | 8.41 | 8.34 | 6.95 | 0.24 | 5.49 | 8.64 | 8.20 | 7.03 | 0.24 | | | | | | | |
| | primiparous | 5.64 | 8.40 | 8.17 | 7.03 | 0.31 | 5.61 | 8.32 | 8.41 | 6.98 | 0.31 | 6.37 | 8.48 | 8.34 | 7.10 | 0.35 | 0.33 | 0.00 | 0.90 | 0.33 | 0.67 | 0.48 | 0.85 |
| SOCS3 | multiparous | 4.56 | 7.54 | 6.32 | 5.79 | 0.20 | 4.51 | 7.83 | 6.32 | 5.74 | 0.17 | 4.81 | 7.90 | 6.42 | 5.78 | 0.17 | | | | | | | |
| | primiparous | 5.36 | 7.67 | 6.27 | 5.84 | 0.22 | 4.86 | 7.64 | 6.27 | 5.64 | 0.22 | 5.12 | 7.95 | 6.29 | 5.89 | 0.24 | 0.32 | 0.00 | 0.79 | 0.28 | 0.60 | 0.06 | 0.95 |
| SCD | multiparous | 8.06 | 7.53 | 7.06 | 7.66 | 0.13 | 8.11 | 7.68 | 6.96 | 7.39 | 0.11 | 8.09 | 7.98 | 7.10 | 7.60 | 0.11 | | | | | | | |
| | primiparous | 8.04 | 7.52 | 7.19 | 7.59 | 0.14 | 8.19 | 7.88 | 7.05 | 7.63 | 0.14 | 8.11 | 7.86 | 6.92 | 7.63 | 0.15 | 0.54 | 0.00 | 0.10 | 0.57 | 0.29 | 0.99 | 0.89 |
| CASP1 | multiparous | 4.45 | 6.31 | 6.04 | 5.80 | 0.13 | 4.50 | 6.47 | 6.05 | 5.90 | 0.11 | 4.40 | 6.55 | 5.94 | 5.91 | 0.11 | | | | | | | |
| | primiparous | 4.97 | 6.52 | 5.81 | 5.90 | 0.15 | 4.66 | 6.35 | 5.86 | 5.78 | 0.15 | 4.94 | 6.51 | 5.93 | 5.86 | 0.16 | 0.75 | 0.00 | 0.95 | 0.35 | 0.33 | 0.00 | 0.83 |
| CLEC7A | multiparous | 4.83 | 6.01 | 6.50 | 6.13 | 0.15 | 4.94 | 6.32 | 6.55 | 6.19 | 0.13 | 4.76 | 6.53 | 6.31 | 6.21 | 0.13 | | | | | | | |
| | primiparous | 5.54 | 6.44 | 6.36 | 6.26 | 0.17 | 5.21 | 6.28 | 6.52 | 6.15 | 0.17 | 5.62 | 6.52 | 6.48 | 6.26 | 0.19 | 0.69 | 0.00 | 0.56 | 0.02 | 0.33 | 0.00 | 0.52 |
| CCL20 | multiparous | 3.87 | 7.05 | 5.66 | 4.42 | 0.25 | 3.95 | 7.30 | 5.71 | 3.75 | 0.22 | 4.21 | 7.48 | 5.62 | 4.05 | 0.22 | | | | | | | |
| | primiparous | 4.69 | 7.11 | 5.65 | 4.05 | 0.28 | 4.23 | 6.99 | 5.74 | 3.95 | 0.28 | 5.01 | 7.19 | 5.65 | 4.36 | 0.31 | 0.26 | 0.00 | 0.55 | 0.32 | 0.85 | 0.04 | 0.68 |
| STAT4 | multiparous | 4.24 | 5.39 | 4.91 | 4.94 | 0.15 | 4.25 | 5.23 | 4.78 | 4.66 | 0.13 | 4.19 | 4.85 | 4.75 | 4.60 | 0.13 | | | | | | | |
| | primiparous | 4.73 | 5.26 | 4.93 | 4.89 | 0.16 | 4.46 | 5.04 | 4.95 | 4.59 | 0.16 | 4.79 | 5.10 | 4.78 | 4.90 | 0.18 | 0.09 | 0.00 | 0.52 | 0.07 | 0.25 | 0.04 | 0.79 |
| IFNAR2 | multiparous | 4.84 | 6.72 | 7.11 | 6.30 | 0.17 | 5.27 | 6.82 | 7.03 | 6.19 | 0.14 | 5.04 | 6.97 | 7.01 | 6.29 | 0.14 | | | | | | | |
| | primiparous | 5.40 | 6.74 | 6.81 | 6.31 | 0.19 | 5.20 | 6.62 | 6.98 | 6.22 | 0.18 | 5.58 | 6.73 | 7.01 | 6.30 | 0.21 | 0.59 | 0.00 | 0.97 | 0.75 | 0.63 | 0.05 | 0.54 |
| IFNAR1 | multiparous | 5.96 | 7.04 | 7.31 | 6.64 | 0.10 | 6.07 | 7.08 | 7.27 | 6.59 | 0.09 | 6.08 | 7.09 | 7.23 | 6.70 | 0.08 | | | | | | | |
| | primiparous | 6.39 | 6.99 | 7.09 | 6.76 | 0.11 | 6.18 | 6.84 | 7.17 | 6.57 | 0.11 | 6.41 | 7.02 | 7.22 | 6.66 | 0.12 | 0.40 | 0.00 | 0.95 | 0.72 | 0.52 | 0.00 | 0.67 |
| ITGA4 | multiparous | 4.11 | 5.91 | 6.51 | 5.93 | 0.25 | 4.32 | 5.85 | 6.59 | 5.99 | 0.22 | 4.60 | 6.02 | 6.37 | 5.91 | 0.22 | | | | | | | |
| | primiparous | 4.49 | 6.00 | 6.52 | 6.19 | 0.28 | 4.48 | 5.55 | 6.47 | 6.03 | 0.28 | 4.61 | 6.20 | 6.47 | 6.10 | 0.31 | 0.67 | 0.00 | 0.74 | 0.50 | 0.68 | 0.86 | 0.99 |
| MMP9 | multiparous | 5.07 | 5.61 | 5.41 | 5.22 | 0.24 | 5.06 | 5.69 | 5.10 | 5.42 | 0.21 | 5.09 | 5.81 | 6.28 | 5.51 | 0.21 | | | | | | | |
| | primiparous | 5.40 | 5.47 | 5.90 | 5.01 | 0.27 | 5.17 | 5.56 | 6.01 | 4.95 | 0.26 | 5.39 | 5.44 | 6.00 | 5.62 | 0.30 | 0.05 | 0.00 | 0.60 | 0.62 | 0.73 | 0.09 | 0.31 |
| CXCL2 | multiparous | 4.53 | 7.58 | 7.90 | 6.93 | 0.29 | 4.80 | 7.69 | 7.74 | 6.66 | 0.26 | 5.08 | 7.91 | 7.84 | 6.99 | 0.25 | | | | | | | |
| | primiparous | 5.89 | 7.73 | 7.70 | 6.99 | 0.33 | 5.19 | 7.48 | 7.84 | 6.74 | 0.33 | 6.07 | 7.64 | 7.79 | 6.93 | 0.36 | 0.29 | 0.00 | 0.89 | 0.19 | 0.75 | 0.01 | 0.81 |
| IL10 | multiparous | 3.30 | 6.15 | 5.98 | 4.98 | 0.20 | 3.63 | 6.20 | 5.94 | 5.05 | 0.17 | 3.81 | 6.35 | 5.94 | 5.16 | 0.17 | | | | | | | |
| | primiparous | 4.17 | 6.28 | 5.68 | 5.20 | 0.22 | 3.93 | 6.03 | 5.90 | 4.97 | 0.22 | 4.18 | 6.31 | 5.98 | 5.14 | 0.25 | 0.29 | 0.00 | 0.98 | 0.26 | 0.55 | 0.05 | 0.83 |
| SIRT1 | multiparous | 6.10 | 6.19 | 6.66 | 6.11 | 0.08 | 6.05 | 6.21 | 6.56 | 6.00 | 0.07 | 6.15 | 6.35 | 6.55 | 6.20 | 0.07 | | | | | | | |
| | primiparous | 6.11 | 6.12 | 6.55 | 6.22 | 0.09 | 5.99 | 6.19 | 6.56 | 6.09 | 0.09 | 6.09 | 6.09 | 6.49 | 6.17 | 0.10 | 0.35 | 0.00 | 0.66 | 0.31 | 0.47 | 0.37 | 0.94 |
| IRF3 | multiparous | 6.41 | 6.35 | 6.20 | 6.30 | 0.10 | 6.39 | 6.41 | 6.20 | 6.23 | 0.09 | 6.44 | 6.42 | 6.22 | 6.24 | 0.09 | | | | | | | |
| | primiparous | 6.38 | 6.35 | 6.18 | 5.84 | 0.12 | 6.48 | 6.39 | 6.11 | 6.27 | 0.11 | 6.39 | 6.44 | 6.20 | 6.27 | 0.13 | 0.43 | 0.00 | 0.81 | 0.40 | 0.48 | 0.66 | 0.40 |
| IL33 | multiparous | 5.90 | 5.45 | 5.38 | 5.22 | 0.19 | 6.18 | 5.50 | 5.12 | 5.25 | 0.17 | 6.09 | 5.72 | 5.26 | 5.39 | 0.17 | | | | | | | |
| | primiparous | 6.03 | 5.52 | 5.48 | 5.53 | 0.22 | 5.96 | 5.46 | 5.62 | 5.37 | 0.22 | 6.17 | 5.85 | 5.21 | 5.63 | 0.24 | 0.77 | 0.00 | 0.15 | 0.42 | 0.98 | 0.41 | 0.29 |
| IGFBP5 | multiparous | 3.84 | 3.83 | 2.62 | 3.20 | 0.27 | 3.90 | 3.91 | 2.99 | 3.21 | 0.24 | 3.78 | 4.17 | 2.82 | 3.20 | 0.24 | | | | | | | |
| | primiparous | 3.64 | 3.94 | 2.71 | 3.40 | 0.30 | 3.98 | 4.28 | 2.87 | 3.25 | 0.30 | 4.21 | 4.39 | 3.33 | 3.64 | 0.34 | 0.17 | 0.00 | 0.98 | 0.17 | 0.46 | 0.98 | 0.96 |
| TAP | multiparous | 4.05 | 5.17 | 6.43 | 4.99 | 0.32 | 4.28 | 5.47 | 6.35 | 4.85 | 0.27 | 4.13 | 5.42 | 6.08 | 5.10 | 0.27 | | | | | | | |
| | primiparous | 5.01 | 5.72 | 6.31 | 5.29 | 0.36 | 4.39 | 4.98 | 5.47 | 4.57 | 0.35 | 4.92 | 5.31 | 5.80 | 4.84 | 0.39 | 0.33 | 0.00 | 0.96 | 0.90 | 0.18 | 0.02 | 0.98 |
| NFkB2 | multiparous | 3.33 | 5.39 | 5.25 | 4.08 | 0.43 | 4.14 | 6.11 | 4.29 | 3.70 | 0.37 | 3.51 | 5.86 | 5.32 | 3.73 | 0.37 | | | | | | | |
| | primiparous | 4.53 | 5.96 | 4.55 | 2.69 | 0.48 | 3.39 | 5.84 | 4.49 | 2.37 | 0.48 | 3.63 | 6.02 | 4.21 | 4.00 | 0.53 | 0.55 | 0.00 | 0.54 | 0.23 | 0.56 | 0.12 | 0.09 |
| DEFB5 | multiparous | 4.93 | 4.74 | 6.51 | 6.35 | 0.45 | 5.22 | 5.14 | 7.09 | 6.96 | 0.39 | 5.09 | 5.78 | 6.47 | 6.90 | 0.39 | | | | | | | |
| | primiparous | 5.53 | 5.71 | 7.15 | 7.02 | 0.51 | 5.59 | 5.65 | 6.19 | 6.83 | 0.50 | 5.42 | 4.82 | 4.84 | 5.02 | 0.56 | 0.16 | 0.00 | 0.33 | 0.65 | 0.02 | 0.10 | 0.50 |
| LBP | multiparous | 5.59 | 5.06 | 5.01 | 5.03 | 0.14 | 5.45 | 5.18 | 4.77 | 4.74 | 0.12 | 5.40 | 5.21 | 4.99 | 4.84 | 0.12 | | | | | | | |
| | primiparous | 5.58 | 5.28 | 5.19 | 4.85 | 0.16 | 5.49 | 5.48 | 4.98 | 5.13 | 0.15 | 5.24 | 5.41 | 4.99 | 4.99 | 0.17 | 0.74 | 0.00 | 0.33 | 0.13 | 0.44 | 0.34 | 0.77 |
| NLRP3 | multiparous | 4.47 | 7.43 | 6.41 | 5.97 | 0.17 | 4.49 | 7.58 | 6.30 | 5.83 | 0.15 | 4.47 | 7.60 | 6.27 | 5.95 | 0.15 | | | | | | | |
| | primiparous | 5.31 | 7.51 | 6.21 | 5.96 | 0.19 | 4.88 | 7.41 | 6.23 | 5.79 | 0.19 | 5.18 | 7.58 | 6.32 | 5.86 | 0.21 | 0.51 | 0.00 | 0.96 | 0.14 | 0.66 | 0.00 | 0.91 |
| CXCL10 | multiparous | 2.66 | 5.67 | 5.17 | 4.24 | 0.54 | 2.43 | 5.64 | 5.02 | 4.45 | 0.47 | 3.16 | 5.70 | 4.45 | 4.69 | 0.46 | | | | | | | |
| | primiparous | 2.95 | 5.96 | 5.14 | 4.28 | 0.60 | 2.35 | 5.83 | 4.91 | 3.62 | 0.60 | 2.06 | 5.60 | 4.60 | 4.61 | 0.66 | 0.77 | 0.00 | 0.84 | 0.68 | 0.78 | 0.86 | 0.91 |
| HSPA8 | multiparous | 8.00 | 7.82 | 8.09 | 8.01 | 0.06 | 8.05 | 7.85 | 8.02 | 7.93 | 0.05 | 8.03 | 7.82 | 7.98 | 8.02 | 0.05 | | | | | | | |
| | primiparous | 8.00 | 7.65 | 8.11 | 8.06 | 0.07 | 8.03 | 7.74 | 8.01 | 7.97 | 0.06 | 8.06 | 7.77 | 7.97 | 8.03 | 0.07 | 0.84 | 0.00 | 0.22 | 0.47 | 0.94 | 0.19 | 0.97 |
| CASP8 | multiparous | 6.33 | 6.15 | 6.33 | 6.25 | 0.06 | 6.35 | 6.27 | 6.31 | 6.19 | 0.05 | 6.30 | 6.25 | 6.24 | 6.26 | 0.05 | | | | | | | |
| | primiparous | 6.33 | 6.26 | 6.18 | 6.32 | 0.06 | 6.20 | 6.26 | 6.22 | 6.26 | 0.06 | 6.34 | 6.27 | 6.23 | 6.20 | 0.07 | 0.92 | 0.13 | 0.70 | 0.58 | 0.59 | 0.21 | 0.35 |
| NFkB1 | multiparous | 5.46 | 6.66 | 6.04 | 5.78 | 0.15 | 5.28 | 6.87 | 5.94 | 5.59 | 0.13 | 5.10 | 6.71 | 6.11 | 5.75 | 0.13 | | | | | | | |
| | primiparous | 5.50 | 6.74 | 5.92 | 5.73 | 0.16 | 5.34 | 6.65 | 5.89 | 5.55 | 0.16 | 5.55 | 6.78 | 5.98 | 5.59 | 0.18 | 0.64 | 0.00 | 0.77 | 0.93 | 0.82 | 0.22 | 0.66 |
| **milk and mammary epithelium integrity gene expression** | | | | | | | | | | | | | | | | | | | | | | | |
| ABCG2 | multiparous | 7.72 | 7.33 | 6.25 | 7.09 | 0.12 | 7.72 | 7.16 | 6.02 | 6.71 | 0.11 | 7.67 | 7.34 | 6.26 | 6.97 | 0.11 | | | | | | | |
| | primiparous | 7.54 | 6.91 | 6.37 | 7.04 | 0.14 | 7.70 | 7.41 | 6.27 | 7.01 | 0.14 | 7.58 | 7.36 | 6.07 | 7.04 | 0.15 | 0.87 | 0.00 | 0.13 | 0.96 | 0.10 | 0.40 | 0.37 |
| BCL2 | multiparous | 4.64 | 4.49 | 5.86 | 5.42 | 0.14 | 4.84 | 4.58 | 6.00 | 5.44 | 0.13 | 4.55 | 4.67 | 5.84 | 5.32 | 0.13 | | | | | | | |
| | primiparous | 5.05 | 4.66 | 5.91 | 5.69 | 0.16 | 4.51 | 4.58 | 5.74 | 5.38 | 0.16 | 4.90 | 4.84 | 5.94 | 5.54 | 0.18 | 0.56 | 0.00 | 0.81 | 0.21 | 0.03 | 0.65 | 0.84 |
| Bax | multiparous | 5.09 | 5.38 | 6.57 | 6.28 | 0.17 | 5.16 | 5.41 | 6.58 | 6.30 | 0.15 | 5.30 | 5.24 | 6.50 | 6.31 | 0.15 | | | | | | | |
| | primiparous | 5.48 | 5.54 | 6.52 | 6.46 | 0.19 | 5.24 | 5.42 | 5.82 | 6.29 | 0.19 | 5.45 | 4.96 | 6.54 | 6.36 | 0.21 | 0.30 | 0.00 | 0.23 | 0.97 | 0.16 | 0.15 | 0.51 |
| CASP1P3 | multiparous | 6.51 | 7.22 | 7.41 | 6.69 | 0.17 | 6.58 | 7.90 | 7.35 | 6.61 | 0.15 | 6.57 | 8.06 | 7.31 | 6.73 | 0.15 | | | | | | | |
| | primiparous | 6.71 | 7.85 | 7.15 | 6.62 | 0.19 | 6.57 | 7.77 | 6.67 | 6.60 | 0.19 | 6.72 | 7.86 | 7.31 | 6.64 | 0.21 | 0.24 | 0.00 | 0.35 | 0.62 | 0.20 | 0.14 | 0.31 |
| CDH1 | multiparous | 5.37 | 5.24 | 5.23 | 5.38 | 0.16 | 5.46 | 5.41 | 5.22 | 5.09 | 0.14 | 5.55 | 5.47 | 5.16 | 5.03 | 0.14 | | | | | | | |
| | primiparous | 5.66 | 5.47 | 5.24 | 5.30 | 0.18 | 5.28 | 5.55 | 4.88 | 5.33 | 0.17 | 5.59 | 5.58 | 5.17 | 5.14 | 0.20 | 0.66 | 0.00 | 0.44 | 0.56 | 0.71 | 0.49 | 0.61 |
| CLDN1 | multiparous | 4.39 | 5.56 | 4.79 | 4.97 | 0.19 | 4.42 | 5.79 | 4.84 | 4.77 | 0.17 | 4.63 | 5.60 | 4.19 | 4.52 | 0.17 | | | | | | | |
| | primiparous | 4.84 | 5.56 | 4.53 | 5.08 | 0.21 | 4.80 | 5.37 | 4.64 | 4.69 | 0.21 | 5.02 | 5.62 | 4.49 | 4.96 | 0.24 | 0.73 | 0.00 | 0.20 | 0.32 | 0.22 | 0.08 | 0.88 |
| CSN1S1 | multiparous | 9.65 | 9.26 | 8.50 | 9.04 | 0.11 | 9.54 | 9.17 | 8.35 | 8.75 | 0.10 | 9.51 | 9.40 | 8.56 | 9.00 | 0.09 | | | | | | | |
| | primiparous | 9.69 | 9.05 | 8.69 | 9.18 | 0.12 | 9.72 | 9.41 | 8.55 | 8.86 | 0.12 | 9.55 | 9.54 | 8.53 | 9.11 | 0.14 | 0.23 | 0.00 | 0.01 | 0.11 | 0.52 | 0.94 | 0.56 |
| CSN3 | multiparous | 9.70 | 9.20 | 8.51 | 9.11 | 0.12 | 9.67 | 9.28 | 8.37 | 8.83 | 0.10 | 9.59 | 9.52 | 8.52 | 9.03 | 0.10 | | | | | | | |
| | primiparous | 9.80 | 9.08 | 8.76 | 9.20 | 0.13 | 9.79 | 9.50 | 8.45 | 8.95 | 0.13 | 9.74 | 9.60 | 8.50 | 9.10 | 0.15 | 0.39 | 0.00 | 0.00 | 0.13 | 0.87 | 0.97 | 0.78 |
| CTSB | multiparous | 6.33 | 6.60 | 8.17 | 8.17 | 0.14 | 6.57 | 6.73 | 8.14 | 8.15 | 0.12 | 6.55 | 6.68 | 8.16 | 8.11 | 0.12 | | | | | | | |
| | primiparous | 6.91 | 6.72 | 8.18 | 8.27 | 0.16 | 6.79 | 6.72 | 8.02 | 8.12 | 0.16 | 6.88 | 6.57 | 8.10 | 8.15 | 0.18 | 0.98 | 0.00 | 0.89 | 0.29 | 0.58 | 0.03 | 0.99 |
| FABP3 | multiparous | 8.72 | 8.11 | 7.59 | 7.88 | 0.17 | 8.67 | 7.93 | 7.28 | 7.50 | 0.14 | 8.61 | 8.06 | 7.43 | 7.74 | 0.14 | | | | | | | |
| | primiparous | 8.50 | 7.82 | 7.70 | 7.41 | 0.19 | 8.75 | 8.37 | 7.35 | 7.83 | 0.18 | 8.47 | 8.17 | 7.39 | 7.83 | 0.20 | 1.00 | 0.00 | 0.19 | 0.96 | 0.13 | 0.76 | 0.48 |
| FAS | multiparous | 5.48 | 6.76 | 6.93 | 5.91 | 0.14 | 5.33 | 6.89 | 6.85 | 5.83 | 0.12 | 5.18 | 7.03 | 6.83 | 5.96 | 0.12 | | | | | | | |
| | primiparous | 5.97 | 6.95 | 6.65 | 6.04 | 0.15 | 5.73 | 6.73 | 6.77 | 5.86 | 0.15 | 5.80 | 6.93 | 6.86 | 5.97 | 0.17 | 0.55 | 0.00 | 0.58 | 0.17 | 0.82 | 0.00 | 0.70 |
| LALBA | multiparous | 9.58 | 9.17 | 8.35 | 8.93 | 0.38 | 9.58 | 9.14 | 8.21 | 8.73 | 0.33 | 9.47 | 9.26 | 8.37 | 8.92 | 0.33 | | | | | | | |
| | primiparous | 9.64 | 8.92 | 8.68 | 9.12 | 0.43 | 9.71 | 9.42 | 7.02 | 7.27 | 0.43 | 9.63 | 9.50 | 8.47 | 9.11 | 0.48 | 0.22 | 0.00 | 0.03 | 0.69 | 0.37 | 0.47 | 0.12 |
| LPL | multiparous | 6.94 | 6.73 | 6.43 | 6.81 | 0.10 | 7.05 | 6.64 | 6.36 | 6.57 | 0.09 | 6.91 | 6.67 | 6.47 | 6.66 | 0.09 | | | | | | | |
| | primiparous | 6.83 | 6.45 | 6.25 | 6.74 | 0.11 | 7.03 | 6.86 | 6.24 | 6.77 | 0.11 | 6.94 | 6.83 | 6.20 | 6.69 | 0.13 | 0.82 | 0.00 | 0.30 | 0.56 | 0.23 | 0.12 | 0.49 |
| OCLN | multiparous | 4.68 | 3.84 | 3.66 | 4.20 | 0.19 | 4.38 | 4.19 | 3.42 | 3.84 | 0.16 | 4.43 | 4.35 | 3.74 | 4.00 | 0.16 | | | | | | | |
| | primiparous | 4.45 | 3.98 | 3.84 | 3.88 | 0.21 | 4.70 | 4.41 | 3.67 | 4.22 | 0.21 | 4.19 | 4.36 | 3.79 | 4.16 | 0.23 | 0.87 | 0.00 | 0.10 | 0.45 | 0.26 | 0.76 | 0.77 |
| RIPK1 | multiparous | 5.68 | 5.51 | 5.42 | 5.43 | 0.10 | 5.57 | 5.49 | 5.43 | 5.33 | 0.09 | 5.26 | 5.58 | 5.45 | 5.39 | 0.09 | | | | | | | |
| | primiparous | 5.60 | 5.45 | 5.35 | 5.55 | 0.12 | 5.39 | 5.40 | 5.40 | 5.31 | 0.12 | 5.07 | 5.77 | 5.46 | 5.39 | 0.13 | 0.44 | 0.06 | 0.00 | 0.61 | 0.83 | 0.35 | 0.81 |
| STAT5A | multiparous | 6.47 | 6.24 | 5.87 | 5.90 | 0.07 | 6.47 | 6.28 | 5.73 | 5.65 | 0.06 | 6.45 | 6.19 | 5.89 | 5.78 | 0.06 | | | | | | | |
| | primiparous | 6.37 | 6.18 | 5.80 | 5.81 | 0.08 | 6.38 | 6.23 | 5.79 | 5.75 | 0.07 | 6.24 | 6.18 | 5.80 | 5.73 | 0.08 | 0.49 | 0.00 | 0.15 | 0.14 | 0.47 | 0.34 | 0.85 |
| SPARC | multiparous | 3.57 | 2.94 | 3.60 | 3.56 | 0.21 | 3.45 | 3.33 | 3.47 | 3.07 | 0.19 | 3.33 | 3.70 | 3.55 | 3.14 | 0.19 | | | | | | | |
| | primiparous | 3.52 | 2.69 | 3.11 | 3.31 | 0.24 | 3.75 | 3.35 | 3.48 | 3.64 | 0.24 | 3.47 | 3.77 | 3.43 | 3.67 | 0.27 | 0.20 | 0.37 | 0.03 | 0.70 | 0.11 | 0.24 | 0.96 |
| TJP1 | multiparous | 6.49 | 6.03 | 5.00 | 5.77 | 0.14 | 6.48 | 5.91 | 4.78 | 5.32 | 0.12 | 6.54 | 5.96 | 4.93 | 5.56 | 0.12 | | | | | | | |
| | primiparous | 6.30 | 5.63 | 4.92 | 5.48 | 0.15 | 6.36 | 5.91 | 5.03 | 5.55 | 0.15 | 6.16 | 5.88 | 4.59 | 5.55 | 0.17 | 0.85 | 0.00 | 0.46 | 0.17 | 0.17 | 0.41 | 0.39 |
| **Reference gene expression** | | | | | | | | | | | | | | | | | | | | | | | |
| GAPDH | multiparous | 6.77 | 6.77 | 8.04 | 7.57 | 0.07 | 6.93 | 6.91 | 8.01 | 7.65 | 0.06 | 6.92 | 6.82 | 8.13 | 7.61 | 0.06 | | | | | | | |
| | primiparous | 7.02 | 6.92 | 8.00 | 7.65 | 0.08 | 7.05 | 7.00 | 7.95 | 7.57 | 0.08 | 7.08 | 6.93 | 8.11 | 7.52 | 0.09 | 0.52 | 0.00 | 0.27 | 0.15 | 0.57 | 0.03 | 0.98 |
| RPLP0 | multiparous | 8.46 | 7.96 | 7.89 | 7.88 | 0.09 | 8.36 | 7.80 | 7.84 | 7.65 | 0.08 | 8.37 | 7.85 | 7.78 | 7.76 | 0.08 | | | | | | | |
| | primiparous | 8.40 | 7.86 | 7.96 | 7.96 | 0.10 | 8.42 | 8.08 | 7.91 | 7.86 | 0.10 | 8.28 | 8.00 | 7.86 | 7.82 | 0.11 | 0.39 | 0.00 | 0.70 | 0.21 | 0.38 | 0.37 | 0.71 |
| RPL19 | multiparous | 8.54 | 8.07 | 8.08 | 7.93 | 0.09 | 8.45 | 7.96 | 8.08 | 7.70 | 0.07 | 8.45 | 8.09 | 7.96 | 7.84 | 0.07 | | | | | | | |
| | primiparous | 8.45 | 7.94 | 8.10 | 8.02 | 0.10 | 8.50 | 8.12 | 8.11 | 7.91 | 0.09 | 8.41 | 8.10 | 8.05 | 7.88 | 0.11 | 0.71 | 0.00 | 0.38 | 0.42 | 0.40 | 0.51 | 0.90 |
| reference genes[3] | multiparous | 9.04 | 9.45 | 10.44 | 9.88 | 0.13 | 9.04 | 9.51 | 10.47 | 10.07 | 0.12 | 9.08 | 9.33 | 10.40 | 9.90 | 0.12 | | | | | | | |
| | primiparous | 8.95 | 9.87 | 10.32 | 9.95 | 0.15 | 8.96 | 9.59 | 10.27 | 10.16 | 0.15 | 8.96 | 9.30 | 10.43 | 10.04 | 0.17 | 0.64 | 0.00 | 0.21 | 0.83 | 0.84 | 0.18 | 0.73 |

1 Milk samples were collected before LPS challenge (0 h) and after LPS challenge times (4 h, 28 h and 76 h).

2 Probability: TREAT = effect of treatment (control vs vitamin E vs plant extract group); TIMES = effect of times before (-20 hours) vs after LPS challenge (vs 4 vs 28 vs 76 hours); TREAT × TIMES = the interaction between treatment and times; PAR = effect of parity (primiparous vs multiparous); PAR × TIMES = interaction between parity and times; TREAT × PAR = interaction between treatment and parity; TREAT × PAR × TIMES = interaction between treatment and parity and times.

3 Decimal logarithm of the geometric mean of ACTB (actin beta), YWHAZ (tyrosine 3-monooxygenase/tryptophan 5-monooxygenase activation protein zeta), and PPIA (cyclophilin A) genes.

**Table S4: Plasma haptoglobin and cortisol, hematological profils and functional capacities of immune cells in control unsupplemented group (n = 11), vitamin E supplemented group (n = 13), and plant extract supplemented group (n = 12) dairy cows before (- 20 h)and after LPS challenge times (4 h, 28 h, and 76 h).**

| Item [1] | Parity | control | | | | | vitamin E | | | | | Plant extracts | | | | | *P*-value [2] | | | | | | |
|---|---|---|---|---|---|---|---|---|---|---|---|---|---|---|---|---|---|---|---|---|---|---|---|
| | | -20 | 4 | 28 | 76 | SEM | -20 | 4 | 28 | 76 | SEM | -20 | 4 | 28 | 76 | SEM | TREAT | TIMES | TREAT x TIMES | PAR | TREAT x PAR | PAR x TIMES | TREAT x PAR x TIMES |
| **Plasma stress and inflammation indicators** | | | | | | | | | | | | | | | | | | | | | | | |
| haptoglobin (mg/mL) | multiparous | 0.66 | 0.81 | 1.21 | 0.94 | 0.273 | 1.21 | 0.97 | 1.59 | 1.46 | 0.237 | 0.81 | 0.73 | 1.45 | 1.45 | 0.234 | | | | | | | |
| | primiparous | 0.74 | 0.69 | 0.98 | 0.94 | 0.313 | 0.59 | 0.62 | 1.10 | 0.92 | 0.306 | 1.05 | 0.95 | 1.37 | 1.23 | 0.341 | 0.61 | **< 0.001** | 0.77 | 0.44 | 0.53 | 0.46 | 0.55 |
| cortisol (ng/mL) | multiparous | 18.35 | 43.43 | 14.00 | 20.68 | 6.114 | 14.81 | 53.22 | 20.57 | 20.55 | 4.879 | 19.14 | 59.10 | 22.90 | 23.90 | 4.852 | | | | | | | |
| | primiparous | 10.11 | 50.56 | 17.19 | 21.53 | 6.318 | 10.11 | 69.82 | 15.78 | 22.40 | 6.242 | 12.50 | 75.41 | 20.62 | 17.25 | 6.964 | 0.22 | **< 0.001** | 0.12 | 0.75 | 0.96 | **0.01** | 0.88 |
| **Hematological profils** [3] | | | | | | | | | | | | | | | | | | | | | | | |
| Red blood cells (.$10^{12}$/L) | multiparous | 5.91 | 5.88 | 5.60 | 5.89 | 0.263 | 6.17 | 6.17 | 5.97 | 5.93 | 0.228 | 6.06 | 5.69 | 5.70 | 6.00 | 0.227 | | | | | | | |
| | primiparous | 6.41 | 7.02 | 6.32 | 6.37 | 0.297 | 6.49 | 6.98 | 6.43 | 6.52 | 0.293 | 6.28 | 6.30 | 6.54 | 6.70 | 0.327 | 0.62 | 0.27 | 0.27 | **< 0.001** | 0.92 | 0.25 | 0.89 |
| Platelets (.$10^{9}$/L) | multiparous | 473 | 513 | 421 | 310 | 52.77 | 539 | 551 | 501 | 428 | 45.78 | 532 | 532 | 573 | 353 | 45.36 | | | | | | | |
| | primiparous | 557 | 489 | 543 | 340 | 60.13 | 574 | 530 | 592 | 469 | 58.97 | 515 | 393 | 516 | 345 | 65.71 | 0.30 | **< 0.001** | 0.23 | 0.77 | 0.46 | 0.03 | 0.85 |
| Haemoglobin (.$10^{9}$/L) | multiparous | 92.37 | 93.87 | 89.87 | 93.87 | 3.387 | 97.60 | 97.97 | 95.60 | 94.47 | 2.937 | 96.37 | 92.12 | 91.62 | 96.37 | 2.918 | | | | | | | |
| | primiparous | 96.58 | 99.18 | 97.78 | 99.78 | 3.818 | 98.06 | 107.46 | 97.66 | 98.66 | 3.765 | 96.05 | 98.30 | 101.05 | 102.80 | 4.199 | 0.48 | 0.37 | 0.30 | **0.02** | 0.93 | 0.34 | 0.82 |
| Hematocrit (L/L) | multiparous | 0.27 | 0.26 | 0.25 | 0.27 | 0.010 | 0.28 | 0.28 | 0.27 | 0.27 | 0.009 | 0.28 | 0.26 | 0.26 | 0.28 | 0.009 | | | | | | | |
| | primiparous | 0.28 | 0.28 | 0.28 | 0.28 | 0.011 | 0.28 | 0.30 | 0.28 | 0.29 | 0.011 | 0.28 | 0.28 | 0.29 | 0.30 | 0.012 | 0.47 | 0.26 | 0.22 | **0.03** | 0.94 | 0.50 | 0.84 |
| Mean corpuscular volume (.$10^{-15}$/L) | multiparous | 45.4 | 45.0 | 45.0 | 45.5 | 0.917 | 45.5 | 45.6 | 45.5 | 45.6 | 0.796 | 45.8 | 45.8 | 46.0 | 46.0 | 0.786 | | | | | | | |
| | primiparous | 43.9 | 44.3 | 44.1 | 44.5 | 1.060 | 43.4 | 43.4 | 43.2 | 43.4 | 1.033 | 43.9 | 43.9 | 43.9 | 44.2 | 1.149 | 0.87 | 0.22 | 0.87 | **0.05** | 0.82 | 0.87 | 0.85 |
| Mean hemoglobin concentration (.$10^{9}$/L) | multiparous | 347.4 | 352.9 | 356.3 | 350.3 | 2.529 | 348.9 | 350.4 | 353.6 | 350.4 | 2.191 | 345.9 | 354.8 | 352.6 | 349.6 | 2.184 | | | | | | | |
| | primiparous | 344.3 | 356.9 | 352.3 | 354.1 | 2.812 | 350.9 | 354.5 | 353.1 | 346.9 | 2.791 | 349.7 | 356.2 | 353.7 | 347.2 | 3.116 | 0.83 | **< 0.001** | 0.38 | 0.62 | 0.96 | 0.47 | 0.51 |
| White blood cells (.$10^{9}$/L) | multiparous | 7.31 | 7.28 | 8.78 | 7.68 | 1.028 | 7.25 | 6.96 | 8.04 | 7.66 | 0.892 | 7.01 | 5.91 | 9.36 | 7.23 | 0.883 | | | | | | | |
| | primiparous | 7.82 | 7.60 | 10.60 | 9.18 | 1.172 | 7.65 | 6.95 | 9.85 | 8.63 | 1.149 | 8.06 | 4.74 | 9.41 | 9.34 | 1.281 | 0.78 | **< 0.001** | 0.23 | 0.32 | 0.96 | 0.10 | 0.68 |
| Lymphocytes (.$10^{9}$/L) | multiparous | 3.55 | 2.58 | 3.95 | 3.37 | 0.537 | 3.76 | 2.69 | 3.77 | 3.66 | 0.466 | 3.18 | 2.61 | 4.95 | 3.75 | 0.462 | | | | | | | |
| | primiparous | 3.63 | 3.82 | 6.13 | 4.99 | 0.609 | 4.37 | 3.30 | 4.67 | 4.43 | 0.598 | 3.15 | 2.37 | 3.77 | 4.38 | 0.667 | 0.54 | **< 0.001** | 0.31 | 0.12 | 0.25 | 0.43 | 0.19 |
| Monocytes (.$10^{9}$/L) | multiparous | 0.31 | 0.23 | 0.33 | 0.35 | 0.088 | 0.37 | 0.38 | 0.39 | 0.45 | 0.077 | 0.37 | 0.34 | 0.35 | 0.32 | 0.076 | | | | | | | |
| | primiparous | 0.33 | 0.31 | 0.49 | 0.54 | 0.099 | 0.32 | 0.33 | 0.60 | 0.43 | 0.098 | 0.38 | 0.19 | 0.62 | 0.50 | 0.109 | 0.70 | **< 0.001** | 0.91 | 0.16 | 0.73 | 0.03 | 0.78 |
| Neutrophils (.$10^{9}$/L) | multiparous | 2.93 | 3.93 | 4.10 | 3.41 | 0.590 | 2.24 | 3.21 | 3.25 | 3.02 | 0.512 | 2.83 | 2.45 | 3.44 | 2.55 | 0.508 | | | | | | | |
| | primiparous | 3.24 | 2.85 | 3.52 | 2.87 | 0.666 | 2.52 | 2.93 | 4.08 | 2.76 | 0.656 | 4.36 | 1.94 | 4.52 | 3.72 | 0.732 | 0.71 | **0.01** | 0.10 | 0.67 | 0.37 | 0.11 | 0.84 |
| Eosinophils (.$10^{9}$/L) | multiparous | 0.49 | 0.52 | 0.38 | 0.53 | 0.162 | 0.85 | 0.68 | 0.63 | 0.54 | 0.141 | 0.61 | 0.50 | 0.61 | 0.59 | 0.139 | | | | | | | |
| | primiparous | 0.56 | 0.60 | 0.45 | 0.73 | 0.185 | 0.44 | 0.35 | 0.51 | 0.53 | 0.181 | 0.16 | 0.24 | 0.49 | 0.70 | 0.202 | 0.85 | 0.22 | 0.08 | 0.44 | 0.49 | **0.02** | 0.84 |
| Basophils (.$10^{7}$/L) | multiparous | 0.02 | 0.02 | 0.03 | 0.01 | 0.015 | 0.02 | 0.00 | 0.01 | 0.01 | 0.013 | 0.03 | 0.01 | 0.02 | 0.01 | 0.013 | | | | | | | |
| | primiparous | 0.05 | 0.01 | 0.00 | 0.04 | 0.016 | 0.01 | 0.04 | 0.00 | 0.05 | 0.016 | 0.02 | 0.01 | 0.02 | 0.04 | 0.018 | 0.82 | 0.26 | 0.94 | 0.19 | 0.74 | **0.05** | 0.52 |
| **functional capacities of blood neutrophil and monocyte immune cells** [4] | | | | | | | | | | | | | | | | | | | | | | | |
| classical MHCII$^{-}$ neutrophils (.$10^{9}$/L) [5] | multiparous | 2.79 | 3.74 | 3.95 | 3.28 | 0.565 | 2.13 | 3.09 | 3.13 | 2.91 | 0.490 | 2.72 | 2.37 | 3.40 | 2.46 | 0.515 | | | | | | | |
| | primiparous | 3.08 | 2.74 | 3.31 | 2.74 | 0.638 | 2.42 | 2.79 | 3.91 | 2.68 | 0.628 | 4.21 | 1.88 | 4.36 | 3.57 | 0.701 | 0.72 | **0.01** | 0.11 | 0.70 | 0.36 | 0.13 | 0.84 |
| classical MHCII$^{+}$ neutrophils (.$10^{9}$/L) | multiparous | 0.03 | 0.02 | 0.04 | 0.02 | 0.011 | 0.02 | 0.03 | 0.03 | 0.03 | 0.010 | 0.03 | 0.02 | 0.01 | 0.02 | 0.011 | | | | | | | |
| | primiparous | 0.05 | 0.03 | 0.06 | 0.02 | 0.013 | 0.03 | 0.03 | 0.04 | 0.02 | 0.014 | 0.04 | 0.01 | 0.03 | 0.02 | 0.014 | 0.45 | **< 0.001** | 0.22 | 0.43 | 0.96 | 0.03 | 0.94 |
| Monocytes (%) | multiparous | 3.61 | 1.41 | 3.75 | 3.97 | 0.560 | 3.42 | 2.14 | 3.70 | 3.62 | 0.486 | 3.67 | 1.98 | 3.84 | 3.68 | 0.483 | | | | | | | |
| | primiparous | 3.54 | 2.58 | 4.40 | 2.98 | 0.630 | 3.81 | 2.35 | 4.98 | 4.22 | 0.622 | 4.37 | 1.78 | 3.51 | 4.28 | 0.693 | 0.81 | **< 0.001** | 0.74 | 0.33 | 0.81 | 0.88 | 0.29 |
| ROS production neutrophils MHCII$^{-}$ (MFI) [6] | multiparous | 1.85 | 1.92 | 1.82 | 2.17 | 0.297 | 1.77 | 1.66 | 1.57 | 1.41 | 0.257 | 2.05 | 2.37 | 1.78 | 1.72 | 0.256 | | | | | | | |
| | primiparous | 1.57 | 1.65 | 1.98 | 1.96 | 0.331 | 1.10 | 1.55 | 1.16 | 1.04 | 0.328 | 1.14 | 1.05 | 1.27 | 1.57 | 0.366 | **0.04** | 0.90 | 0.78 | **0.01** | 0.27 | 0.56 | 0.68 |
| ROS production neutrophils MHCII$^{+}$ (MFI) | multiparous | 2.21 | 3.06 | 1.95 | 1.44 | 0.396 | 2.03 | 1.83 | 1.76 | 1.32 | 0.344 | 2.05 | 2.28 | 1.75 | 1.32 | 0.342 | | | | | | | |
| | primiparous | 1.68 | 1.74 | 2.00 | 1.93 | 0.441 | 1.23 | 1.69 | 1.11 | 0.94 | 0.438 | 1.28 | 1.45 | 1.54 | 1.19 | 0.489 | **0.07** | 0.05 | 1.00 | **0.03** | 0.91 | 0.91 | 0.65 |
| ROS production neutrophils MHCII$^{-}$ with TBHP (MFI) | multiparous | 1.91 | 3.61 | 2.86 | 1.06 | 0.654 | 1.35 | 2.26 | 1.66 | 0.88 | 0.567 | 1.36 | 3.46 | 2.79 | 0.89 | 0.564 | | | | | | | |
| | primiparous | 1.30 | 2.49 | 2.55 | 0.75 | 0.733 | 0.33 | 1.98 | 2.12 | 0.34 | 0.725 | 0.99 | 2.76 | 3.24 | 1.32 | 0.809 | 0.16 | **< 0.001** | 0.93 | 0.38 | 0.83 | 0.83 | 0.97 |
| ROS production neutrophils MHCII$^{+}$ with TBHP (MFI) | multiparous | 5.25 | 9.20 | 4.20 | 2.46 | 1.497 | 2.43 | 2.84 | 2.13 | 1.03 | 1.298 | 2.37 | 4.18 | 4.20 | 1.26 | 1.291 | | | | | | | |
| | primiparous | 2.22 | 3.65 | 5.73 | 1.22 | 1.676 | -0.07 | 3.54 | 5.91 | 0.05 | 1.658 | 0.79 | 4.07 | 4.40 | 1.69 | 1.850 | 0.12 | **< 0.001** | 0.89 | 0.41 | 0.45 | 0.45 | 0.53 |
| ROS production monocytes (MFI) | multiparous | 0.32 | 0.38 | 0.28 | 0.27 | 0.135 | 0.31 | 0.31 | 0.31 | 0.77 | 0.117 | 0.32 | 0.33 | 0.35 | 0.31 | 0.116 | | | | | | | |
| | primiparous | 0.33 | 0.56 | 0.33 | 0.36 | 0.150 | 0.37 | 0.36 | 0.40 | 0.38 | 0.149 | 0.42 | 0.46 | 0.44 | 0.41 | 0.166 | 0.81 | 0.76 | 0.52 | 0.46 | 0.53 | 0.70 | 0.79 |
| ROS production monocytes with TBHP (MFI) | multiparous | 0.47 | 0.41 | 0.55 | 0.60 | 0.051 | 0.44 | 0.41 | 0.48 | 0.49 | 0.044 | 0.55 | 0.43 | 0.50 | 0.49 | 0.044 | | | | | | | |
| | primiparous | 0.51 | 0.60 | 0.52 | 0.57 | 0.057 | 0.53 | 0.54 | 0.62 | 0.64 | 0.056 | 0.65 | 0.59 | 0.65 | 0.67 | 0.063 | 0.36 | **0.02** | 0.56 | **< 0.001** | 0.28 | 0.42 | 0.52 |
| Phagocytosis neutrophils MHCII$^{-}$ (MFI) | multiparous | 0.23 | 0.25 | 0.25 | 0.22 | 0.016 | 0.24 | 0.25 | 0.23 | 0.21 | 0.014 | 0.23 | 0.27 | 0.23 | 0.22 | 0.014 | | | | | | | |
| | primiparous | 0.23 | 0.25 | 0.20 | 0.22 | 0.020 | 0.23 | 0.25 | 0.21 | 0.22 | 0.018 | 0.22 | 0.24 | 0.22 | 0.23 | 0.020 | 0.76 | **0.01** | 0.99 | 0.29 | 0.94 | 0.94 | 0.88 |
| Phagocytosis neutrophils MHCII$^{+}$ (MFI) | multiparous | 0.39 | 0.35 | 0.57 | 0.57 | 0.057 | 0.45 | 0.44 | 0.43 | 0.53 | 0.049 | 0.36 | 0.51 | 0.50 | 0.63 | 0.049 | | | | | | | |
| | primiparous | 0.41 | 0.45 | 0.36 | 0.52 | 0.071 | 0.38 | 0.40 | 0.41 | 0.59 | 0.063 | 0.36 | 0.38 | 0.43 | 0.60 | 0.071 | 0.87 | **< 0.001** | 0.61 | 0.29 | 0.90 | 0.90 | 0.16 |
| Phagocytosis monocytes (MFI) | multiparous | 0.53 | 0.48 | 0.61 | 0.64 | 0.049 | 0.55 | 0.49 | 0.60 | 0.62 | 0.042 | 0.53 | 0.54 | 0.58 | 0.58 | 0.042 | | | | | | | |
| | primiparous | 0.61 | 0.66 | 0.60 | 0.60 | 0.061 | 0.56 | 0.58 | 0.68 | 0.67 | 0.054 | 0.54 | 0.56 | 0.52 | 0.62 | 0.060 | 0.37 | **0.04** | 0.84 | 0.12 | 0.59 | 0.42 | 0.59 |

1 Plasma and blood samples were collected before LPS challenge (-20 h) and after LPS challenge times (4 h, 28 h and 76 h).
2 Probability: TREAT = effect of treatment (control vs vitamin E vs plant extract group); TIMES = effect of times before (-20 hours) vs after LPS challenge (vs 4 vs 28 vs 76 hours);
TREAT × TIMES = the interaction between treatment and times; PAR = effect of parity (primiparous vs multiparous); PAR × TIMES = interaction between parity and times; TREAT × PAR = interaction between treatment and parity;
TREAT × PAR × TIMES = interaction between treatment and parity and times.
3 Blood hematological profils by Labocea laboratory.
4 Functional capacities of blood neutrophils and monocyte immune cells measuring by flow cytometry. ROS production was measured with or without tert butyl hydroperoxide (TBHP) stimulation. Phagocytosis was measured with beads with Escherichia coli lipopolysaccharide.
5 The count of the two MHCII- et MHCII+ neutrophil subpopulations (.109/L) was calculated from a percentage measured by flow cytometry multiplied by the neutrophil count given by hematological profiles.
6 MFI = Mean Fluorescence intensitie by flow cytometry.

Table S5: Measurement of rectal temperature, milk content and biomarkers of the mammary epithelium integrity in control unsupplemented group (n = 11), supplemented vitamin E group (n = 13), and supplemented plant extract group (n = 12) dairy cows before (0 h) and during LPS challenge times (4 h, 7 h, 9 h, 28 h, 52 h and 76 h) of the only challenged mammary quarter. The milk content measured milk somatic cell count (SCC), fat and protein milk fat and protein milk ratio, lactose milk and β-hydroxybutyrate. The biomarkers of mammary epithelial integrity measured by milk concentrations of Na+, K+, Na+:K+ ratio, plasma lactose, and other milk minerals.

| Item [1] | Parity | control | | | | | | | | vitamin E | | | | | | | | Plant extracts | | | | | | | | *P* -value [2] | | | | | | |
|---|---|---|---|---|---|---|---|---|---|---|---|---|---|---|---|---|---|---|---|---|---|---|---|---|---|---|---|---|---|---|---|---|
| | | 0 | 4 | 7 | 9 | 28 | 52 | 76 | SEM | 0 | 4 | 7 | 9 | 28 | 52 | 76 | SEM | 0 | 4 | 7 | 9 | 28 | 52 | 76 | SEM | TREAT | TIMES | TREAT x TIMES | PAR | TREAT x PAR | PAR x TIMES | TREAT x PAR x TIMES |
| Rectal temperature (°C) | multiparous | 37.93 | 39.38 | 40.30 | 39.22 | 38.32 | 38.44 | 38.30 | 0.166 | 38.30 | 39.45 | 40.00 | 39.28 | 38.48 | 38.48 | 38.46 | 0.165 | 38.14 | 39.60 | 40.20 | 39.28 | 38.32 | 38.39 | 38.36 | 0.191 | | | | | | | |
| | primiparous | 38.37 | 39.39 | 40.55 | 39.65 | 38.29 | 38.56 | 38.55 | 0.211 | 38.29 | 39.13 | 40.13 | 39.45 | 38.27 | 38.43 | 38.35 | 0.236 | 38.26 | 40.04 | 40.64 | 39.31 | 38.43 | 38.36 | 38.36 | 0.213 | 0.59 | **< 0.001** | 0.24 | 0.25 | 0.38 | 0.71 | 0.94 |
| Milk composition | | | | | | | | | | | | | | | | | | | | | | | | | | | | | | | | |
| SCC (.10$^3$/mL) | multiparous | 64.37 | 1001 | 13750 | 7737 | 16381 | 6069 | 3783 | 1804.64 | -194.03 | 1687 | 16596 | 11839 | 18948 | 7397 | 3062 | 1800.92 | 6.37 | 2892 | 14808 | 10026 | 12929 | 6143 | 4885 | 1796.78 | | | | | | | |
| | primiparous | 373.59 | 3647 | 13845 | 12655 | 10200 | 10733 | 4482 | 1999.39 | -160.74 | 2641 | 8135 | 7754 | 9694 | 6247 | 2682 | 2211.71 | -265 | 2854 | 12401 | 10238 | 5897 | 5527 | 1149 | 2220.74 | 0.60 | **< 0.001** | 0.93 | 0.18 | 0.18 | **< 0.001** | 0.89 |
| Milk fat (g/kg) | multiparous | 11.61 | 29.66 | 16.48 | 11.23 | 43.89 | 41.76 | 60.29 | 6.722 | 10.18 | 50.57 | 23.02 | 12.74 | 61.71 | 63.77 | 63.31 | 6.605 | 8.29 | 57.40 | 23.42 | 12.94 | 57.55 | 60.72 | 72.69 | 6.582 | | | | | | | |
| | primiparous | 11.23 | 37.23 | 19.03 | 13.89 | 43.49 | 50.65 | 44.13 | 7.496 | 10.45 | 41.11 | 20.96 | 17.53 | 40.89 | 50.17 | 42.59 | 8.165 | 9.71 | 29.83 | 13.69 | 9.61 | 32.66 | 38.96 | 30.86 | 8.295 | 0.55 | **< 0.001** | 0.99 | 0.11 | 0.30 | **< 0.001** | 0.65 |
| Milk protein (g/kg) | multiparous | 28.04 | 32.86 | 30.91 | 27.59 | 27.59 | 27.06 | 27.09 | 1.035 | 29.79 | 30.79 | 28.75 | 27.72 | 27.12 | 27.36 | 27.37 | 0.986 | 29.05 | 29.08 | 28.61 | 27.10 | 27.06 | 26.64 | 26.89 | 0.980 | | | | | | | |
| | primiparous | 29.69 | 30.57 | 28.91 | 28.17 | 29.43 | 28.99 | 28.91 | 1.167 | 31.39 | 28.79 | 28.27 | 27.91 | 28.43 | 28.51 | 28.37 | 1.234 | 28.88 | 27.61 | 27.08 | 26.25 | 27.15 | 26.83 | 28.15 | 1.283 | 0.46 | **< 0.001** | **0.01** | 0.81 | 0.92 | **0.01** | 0.99 |
| Milk fat:protein | multiparous | 0.39 | 0.88 | 0.52 | 0.40 | 1.64 | 1.60 | 2.36 | 0.255 | 0.34 | 1.70 | 0.79 | 0.45 | 2.26 | 2.36 | 2.32 | 0.252 | 0.28 | 2.04 | 0.82 | 0.48 | 2.13 | 2.29 | 2.71 | 0.251 | | | | | | | |
| | primiparous | 0.39 | 1.28 | 0.67 | 0.49 | 1.51 | 1.76 | 1.55 | 0.284 | 0.33 | 1.40 | 0.73 | 0.61 | 1.42 | 1.75 | 1.48 | 0.310 | 0.32 | 1.08 | 0.49 | 0.36 | 1.20 | 1.45 | 1.10 | 0.314 | 0.63 | **< 0.001** | 0.99 | **0.07** | 0.31 | **< 0.001** | 0.84 |
| Milk lactose (g/kg) | multiparous | 48.53 | 38.28 | 34.39 | 40.51 | 38.56 | 45.08 | 44.43 | 1.922 | 49.95 | 37.63 | 38.94 | 37.75 | 37.81 | 44.11 | 46.00 | 1.903 | 50.11 | 42.51 | 39.81 | 43.27 | 43.84 | 45.70 | 45.66 | 1.897 | | | | | | | |
| | primiparous | 51.86 | 44.46 | 43.02 | 42.58 | 49.04 | 46.94 | 50.38 | 2.137 | 51.46 | 48.62 | 43.31 | 43.26 | 46.94 | 46.18 | 50.22 | 2.345 | 53.63 | 51.19 | 43.80 | 44.98 | 51.53 | 50.30 | 53.05 | 2.369 | **0.06** | **< 0.001** | 0.82 | **< 0.001** | 0.94 | **< 0.001** | 0.98 |
| Milk β-hydroxybutyrate (g/kg) | multiparous | 0.11 | 0.10 | 0.25 | 0.24 | 0.14 | 0.36 | 0.22 | 0.086 | 0.09 | 0.21 | 0.18 | 0.26 | 0.46 | 0.14 | 0.15 | 0.083 | 0.07 | 0.18 | 0.24 | 0.18 | 0.21 | 0.17 | 0.22 | 0.082 | | | | | | | |
| | primiparous | 0.09 | 0.13 | 0.20 | 0.22 | 0.11 | 0.13 | 0.13 | 0.097 | 0.13 | 0.17 | 0.31 | 0.22 | 0.19 | 0.18 | 0.15 | 0.103 | 0.13 | 0.18 | 0.18 | 0.17 | 0.14 | 0.15 | 0.16 | 0.106 | 0.84 | **0.05** | 0.44 | 0.52 | 0.94 | 0.56 | 0.56 |
| Biomarkers of mammary epithelium integrity | | | | | | | | | | | | | | | | | | | | | | | | | | | | | | | | |
| Milk $Na^+$ (mg/kg) | multiparous | 307 | 1176 | 1200 | 1131 | 1112 | 611 | 592 | 138.1 | 365 | 1212 | 1337 | 1250 | 1345 | 891 | 657 | 119.8 | 286 | 801 | 868 | 867 | 724 | 547 | 459 | 119.0 | | | | | | | |
| | primiparous | 347 | 1123 | 1083 | 1091 | 879 | 674 | 612 | 155.5 | 272 | 642 | 819 | 1028 | 829 | 663 | 445 | 153.4 | 336 | 1203 | 1506 | 1145 | 955 | 773 | 594 | 171.1 | 0.80 | **< 0.001** | 0.49 | 0.69 | **0.01** | 0.77 | 0.17 |
| Milk $K^+$ (mg/kg) | multiparous | 1856 | 1239 | 1312 | 1412 | 1324 | 1607 | 1522 | 100.7 | 1811 | 1159 | 1184 | 1342 | 1182 | 1448 | 1499 | 87.32 | 1751 | 1287 | 1336 | 1419 | 1385 | 1417 | 1496 | 86.7 | | | | | | | |
| | primiparous | 1656 | 1117 | 1159 | 1228 | 1319 | 1448 | 1442 | 113.5 | 1648 | 1290 | 1177 | 1032 | 1198 | 1292 | 1425 | 111.9 | 1687 | 1160 | 992 | 1265 | 1308 | 1355 | 1484 | 124.8 | 0.63 | **< 0.001** | 0.63 | 0.10 | 0.94 | 0.30 | 0.61 |
| Milk $Na^+$:$K^+$ ratio | multiparous | 0.16 | 1.06 | 0.94 | 0.90 | 0.87 | 0.38 | 0.39 | 0.244 | 0.20 | 1.18 | 1.41 | 0.94 | 1.25 | 0.65 | 0.44 | 0.211 | 0.16 | 0.60 | 0.66 | 0.62 | 0.52 | 0.39 | 0.31 | 0.211 | | | | | | | |
| | primiparous | 0.24 | 1.93 | 0.98 | 1.00 | 0.79 | 0.54 | 0.46 | 0.271 | 0.18 | 0.50 | 0.87 | 1.10 | 0.66 | 0.52 | 0.32 | 0.269 | 0.22 | 1.04 | 1.58 | 0.99 | 0.79 | 0.59 | 0.42 | 0.300 | 0.60 | **< 0.001** | 0.44 | 0.48 | **0.05** | 0.88 | 0.36 |
| Plasma lactose (mg/L) | multiparous | 37.3 | 46.6 | NE[3] | NE | 45.0 | NE | 35.7 | 4.002 | 30.1 | 39.8 | NE | NE | 39.5 | NE | 37.0 | 3.471 | 45.1 | 52.9 | NE | NE | 40.3 | NE | 43.2 | 3.445 | | | | | | | |
| | primiparous | 40.0 | 45.0 | NE | NE | 36.1 | NE | 31.8 | 4.524 | 29.1 | 52.1 | NE | NE | 33.1 | NE | 30.6 | 4.454 | 30.9 | 45.7 | NE | NE | 38.8 | NE | 33.5 | 4.967 | 0.27 | **< 0.001** | 0.39 | 0.16 | 0.44 | 0.17 | **0.05** |
| Milk cytokines | | | | | | | | | | | | | | | | | | | | | | | | | | | | | | | | |
| IL8 (ng/mL) | multiparous | 276 | 30732 | 34906 | 19264 | 7621 | 1488 | 1357 | 8158 | 979 | 49237 | 52638 | 19655 | 6478 | 2419 | 1322 | 7070 | 4.500 | 60316 | 49618 | 26882 | 3987 | 1836 | 1109 | 7044 | | | | | | | |
| | primiparous | -1599 | 39873 | 72516 | 37270 | 3715 | 797 | -271 | 9082 | 885 | 53519 | 55795 | 46245 | 10604 | 2665 | 1471 | 9009 | 691 | 113159 | 88559 | 52755 | 10646 | 4134 | 2146 | 10059 | **0.04** | **< 0.001** | **< 0.001** | **0.01** | 0.32 | **< 0.001** | 0.28 |
| IL1β (ng/mL) | multiparous | -130 | 812 | 8376 | 5082 | 2088 | 174 | 335 | 1258 | -81 | 2588 | 6128 | 5935 | 1146 | 124 | -30 | 1090 | -1.98E-12 | 2020 | 9636 | 7600 | 1137 | 267 | 34 | 1086 | | | | | | | |
| | primiparous | 446 | 4338 | 11710 | 7997 | 1999 | 862 | 585 | 1399 | 178 | 2263 | 9238 | 2526 | 2139 | 429 | 263 | 1389 | 218 | 2428 | 9899 | 8504 | 2768 | 516 | 289 | 1550 | 0.33 | **< 0.001** | 0.60 | 0.19 | 0.56 | 0.76 | 0.64 |
| Other milk minerals | | | | | | | | | | | | | | | | | | | | | | | | | | | | | | | | |
| Calcium (mg/kg) | multiparous | 1275.5 | 905.0 | 1115 | 1194 | 1059 | 1152 | 1176 | 360.45 | 1259 | 839 | 913 | 1039 | 898 | 1039 | 1094 | 312.97 | 1786 | 1537 | 1615 | 1673 | 1586 | 1604 | 1662 | 308.79 | | | | | | | |
| | primiparous | 1400.2 | 1009.8 | 1249 | 1290 | 1216 | 1237 | 1253 | 417.50 | 2037 | 1790 | 1793 | 1824 | 1794 | 1850 | 1993 | 406.28 | 1099 | 749 | 802 | 925 | 873 | 892 | 967 | 452.097 | 0.78 | **< 0.001** | 0.23 | 0.81 | 0.11 | 0.93 | 0.85 |
| Magnesium (mg/kg) | multiparous | 17.70 | 15.26 | 16.26 | 16.24 | 17.22 | 17.45 | 17.55 | 92.51 | -2.75 | -5.25 | -4.82 | -4.11 | -3.56 | -3.02 | -3.33 | 80.32 | 10.48 | 8.41 | 9.18 | 9.39 | 10.13 | 9.49 | 9.69 | 79.284 | | | | | | | |
| | primiparous | 22.35 | 19.68 | 20.82 | 21.25 | 22.41 | 21.92 | 21.40 | 106.95 | 351 | 310 | 194.79 | 168.1 | 231 | 303 | 309 | 104.17 | -9.40 | -11.30 | -11.50 | -10.83 | -8.97 | -9.95 | -9.35 | 115.93 | 0.31 | 0.18 | 0.13 | 0.30 | 0.23 | 0.21 | 0.13 |
| Phosphorus (mg/kg) | multiparous | 810 | 593 | 702 | 724 | 701 | 772 | 790 | 58.48 | 823 | 625 | 678 | 766 | 682 | 769 | 805 | 50.69 | 806 | 709 | 783 | 760 | 755 | 750 | 786 | 50.47 | | | | | | | |
| | primiparous | 920 | 670 | 804 | 794 | 833 | 859 | 862 | 65.25 | 793 | 717 | 892 | 944 | 883 | 772 | 877 | 64.66 | 872 | 661 | 663 | 721 | 765 | 788 | 820 | 72.18 | 0.71 | **< 0.001** | 0.73 | **0.04** | 0.21 | 0.92 | 0.40 |
| Total phosphorus (mg/kg) | multiparous | 0.87 | 0.64 | 0.73 | 0.75 | 0.73 | 0.79 | 0.83 | 0.039 | 0.90 | 0.644 | 0.71 | 0.81 | 0.71 | 0.77 | 0.84 | 0.034 | 0.94 | 0.75 | 0.83 | 0.82 | 0.81 | 0.81 | 0.84 | 0.0340 | | | | | | | |
| | primiparous | 0.99 | 0.70 | 0.85 | 0.84 | 0.87 | 0.93 | 0.89 | 0.044 | 0.99 | 0.846 | 0.79 | 0.72 | 0.80 | 0.85 | 0.96 | 0.044 | 0.94 | 0.67 | 0.69 | 0.75 | 0.79 | 0.80 | 0.87 | 0.0488 | 0.97 | **< 0.001** | 0.28 | **0.04** | **0.02** | 0.13 | **0.06** |

**1 Milk samples were collected of the only challenged mammary quarter for measurement before (0 h) and during LPS challenge times (4 h, 7 h, 9 h, 28 h, 52 h and 76 h). ). The variation 7 h and 9 h after LPS challenge was linked to the fact that it was cisternal and not alveolar milk. Alveolar milk was at the kinetic point 4 h, 28 h, 52 h and 76 h after LPS challenge.**

**2 Probability: TREAT = effect of treatment control vs vitamin E vs plant extract group); TIMES = effect of times before LPS vs after LPS challenge (4 h vs 7 h vs 9 h vs 28 h vs 52 h vs 76 hours); TREAT × TIMES = the interaction between treatment and times; PAR = effect of parity (primiparous vs multiparous); PAR × TIMES = interaction between parity and times; TREAT × PAR = interaction between treatment and parity; TREAT × PAR × TIMES = interaction between treatment and parity and times.**

**3 NE =Not estimated because no collection.**

**Table S6: Functional capacities of milk neutrophil and macrophage immune cells in control unsupplemented group (n = 11), supplemented vitamin E group (n = 13), and supplemented plant extract group (n = 12) dairy cows during LPS challenge times (4 h and 76 h) of the only challenged mammary quarter.**

| Item [1] | Parity | | | | | | | | | | *P*-value [2] | | | | | | |
|---|---|---|---|---|---|---|---|---|---|---|---|---|---|---|---|---|---|
| | | 4 | 76 | SEM | 4 | 76 | SEM | 4 | 76 | SEM | TREAT | TIMES | TREAT x TIMES | PAR | TREAT x PAR | PAR x TIMES | TREAT x PAR x TIMES |
| functional capacities of milk neutrophil and monocyte immune cells [4] | | | | | | | | | | | | | | | | | |
| classical MHCII$^-$ neutrophils (%) | multiparous | 85.55 | 87.70 | 4.275 | 77.72 | 85.79 | 2.943 | 75.72 | 86.89 | 2.918 | | | | | | | |
| | primiparous | 80.12 | 83.73 | 3.845 | 83.28 | 82.06 | 3.772 | 79.70 | 80.99 | 4.915 | 0.48 | **0.06** | 0.81 | 0.50 | 0.56 | 0.17 | 0.49 |
| classical MHCII$^+$ neutrophils (%) | multiparous | 5.38 | 2.53 | 0.991 | 6.20 | 3.10 | 0.860 | 6.05 | 2.11 | 0.853 | | | | | | | |
| | primiparous | 3.59 | 1.77 | 1.241 | 3.08 | 3.46 | 1.103 | 4.03 | 3.54 | 1.437 | 0.66 | **< 0.001** | 0.76 | 0.15 | 0.76 | **0.04** | 0.66 |
| Macrophages (%) | multiparous | 0.93 | 0.91 | 0.204 | 0.83 | 0.95 | 0.189 | 1.10 | 0.98 | 0.175 | | | | | | | |
| | primiparous | 1.39 | 1.20 | 0.254 | 0.63 | 0.87 | 0.255 | 0.63 | 1.06 | 0.299 | 0.32 | 0.44 | 0.44 | 0.95 | 0.28 | 0.42 | 0.37 |
| ROS production neutrophils MHCII$^-$ (MFI) [3] | multiparous | 0.75 | 0.24 | 0.162 | 0.84 | 0.32 | 0.151 | 0.74 | 0.46 | 0.139 | | | | | | | |
| | primiparous | 0.68 | 0.42 | 0.183 | 0.86 | 0.45 | 0.180 | 1.29 | 0.45 | 0.234 | 0.28 | **< 0.001** | 0.80 | 0.22 | 0.68 | 0.76 | 0.25 |
| ROS production neutrophils MHCII$^+$ (MFI) | multiparous | 0.70 | 0.59 | 0.210 | 0.90 | 0.59 | 0.177 | 1.04 | 0.65 | 0.164 | | | | | | | |
| | primiparous | 1.06 | 0.67 | 0.216 | 1.09 | 0.63 | 0.212 | 1.36 | 0.70 | 0.338 | 0.52 | **< 0.001** | 0.67 | 0.19 | 0.92 | 0.35 | 0.97 |
| ROS production neutrophils MHCII$^-$ with TBHP (MFI) | multiparous | 0.33 | 0.33 | 0.124 | 0.81 | 0.36 | 0.092 | 0.69 | 0.43 | 0.085 | | | | | | | |
| | primiparous | 0.48 | 0.28 | 0.112 | 0.59 | 0.30 | 0.123 | 0.42 | 0.32 | 0.175 | 0.15 | **< 0.001** | 0.17 | 0.20 | 0.36 | 0.75 | 0.42 |
| ROS production neutrophils MHCII$^+$ with TBHP (MFI) | multiparous | 1.11 | 0.73 | 0.231 | 1.13 | 0.65 | 0.215 | 1.10 | 0.74 | 0.199 | | | | | | | |
| | primiparous | 1.14 | 0.61 | 0.262 | 1.24 | 0.78 | 0.257 | 0.72 | 0.75 | 0.410 | 0.80 | **0.02** | 0.68 | 0.83 | 0.72 | 0.77 | 0.77 |
| ROS production macrophages (MFI) | multiparous | 1.46 | 1.72 | 0.405 | 1.22 | 1.40 | 0.352 | 1.17 | 1.29 | 0.349 | | | | | | | |
| | primiparous | 1.27 | 1.10 | 0.458 | 1.09 | 1.57 | 0.451 | 1.13 | 1.63 | 0.586 | 0.96 | 0.37 | 0.87 | 0.76 | 0.66 | 0.86 | 0.76 |
| ROS production macrophages with TBHP (MFI) | multiparous | 0.84 | 1.30 | 0.351 | 1.49 | 1.40 | 0.304 | 1.33 | 1.45 | 0.302 | | | | | | | |
| | primiparous | 1.32 | 1.24 | 0.396 | 0.92 | 1.01 | 0.390 | 1.09 | 1.45 | 0.508 | 0.85 | 0.51 | 0.88 | 0.58 | 0.42 | 0.93 | 0.72 |
| Phagocytosis neutrophils MHCII$^-$ (MFI) | multiparous | 0.23 | 0.28 | 0.025 | 0.23 | 0.30 | 0.022 | 0.25 | 0.30 | 0.021 | | | | | | | |
| | primiparous | 0.20 | 0.23 | 0.028 | 0.23 | 0.30 | 0.028 | 0.21 | 0.30 | 0.036 | 0.36 | **< 0.001** | 0.56 | 0.21 | 0.60 | 0.86 | 0.73 |
| Phagocytosis neutrophils MHCII$^+$ (MFI) | multiparous | 0.76 | 1.33 | 0.154 | 0.66 | 0.85 | 0.146 | 0.90 | 0.98 | 0.133 | | | | | | | |
| | primiparous | 0.79 | 0.92 | 0.175 | 0.74 | 0.77 | 0.172 | 0.52 | 1.00 | 0.223 | 0.27 | **0.02** | 0.58 | 0.25 | 0.69 | 0.74 | 0.22 |
| Phagocytosis macrophages (MFI) | multiparous | 1.06 | 0.92 | 0.115 | 1.14 | 0.75 | 0.108 | 1.04 | 0.98 | 0.099 | | | | | | | |
| | primiparous | 1.10 | 1.03 | 0.131 | 1.09 | 0.98 | 0.129 | 0.90 | 0.67 | 0.168 | 0.49 | **0.03** | 0.66 | 0.83 | 0.28 | 0.68 | 0.46 |

**1 Immune cell samples of milk were collected of the only challenged mammary quarter during LPS challenge times (4 h and 76 h).**

**2 Probability: TREAT = effect of treatment (control vs vitamin E vs plant extract group); TIMES = effect of times after LPS challenge (4 vs 76 hours); TREAT × TIMES = the interaction between treatment and times; PAR = effect of parity (primiparous vs multiparous); PAR × TIMES = interaction between parity and times; TREAT × PAR = interaction between treatment and parity; TREAT × PAR × TIMES = interaction between treatment and parity and times.**

**3 MFI = Mean Fluorescence intensitie by flow cytometry.**

**Table S7: Composition on dry matter ingredients and the chemical composition of the diet during close-up, first 15 DIM (days in milk) and after 15 DIM in dairy cows.**

| Item [1] | Close-up [2] | Lactation first 15 DIM | Lactation after 15 DIM |
|---|---|---|---|
| **Dry matter ingredients** | | | |
| Corn silage (% DM) | 72 | 65 | 65 |
| Energy concentrate (% DM) | 14 | 12.5 | 12.5 |
| Soybean meal (% DM) | 14 | 12.5 | 12.5 |
| Dehydrated alfalfa (% DM) | 0 | 10 | 10 |
| Straw (kg DM/day) | 1 | 1 | 0 |
| Mineral feed dry (g DM/day) [3] | 0.15 | 0 | 0 |
| Mineral feed lactation (g DM/day) [4] | 0 | 0.2 | 0.2 |
| Magnesium chloride (g DM/day) | 0.1 | 0 | 0 |
| Magnesium oxide (g DM/day) | 0 | 0.05 | 0.05 |
| Carbonates (g/kg DM) | 0 | 0.05 | 0.05 |
| **Chemical composition of the diet** | | | |
| DMI (kg/day) | | | |
| $NE_L$ (MJ/kg DMI) [5] | | | |
| $NE_L$ (MJ/day) [5] | | | |
| DPI (g/kg DM) [6] | | | |
| DPI (g/day) [6] | | | |
| DPI/FU (g/FU) [6] | | | |
| Nitrogen efficiency (g/g) | | | |
| Calcium (g/kg DMI) | | | |
| Phosphorus (g/kg DMI) | | | |
| Magnesium (g/kg DMI) | | | |
| DCAD (mEq/kg DMI) [7] | | | |
| Selenium (g/kg DMI) | | | |
| Vitamin E (IU/day) | | | |

[1] With this diet composition, cows in the vitamin E group were supplemented with the all-rac-alpha-tocopheryl acetate

[2] Before the close-up period (far-off period), the dry matter ingredients were corn silage (6.46% DM), grass silage

[3] The mineral feed lactation composition was the same before and after calving and was composed of calcium

[4] The mineral feed lactation composition was the same before and after calving and was composed of calcium

[5] $NE_L$: Net energy for milk production in MJ/kg of DM or MJ/day.

[6] DPI: Digestible proteins in the intestine in g/kg of DM, or g/day, or g/FU (fodder unit) (INRA, 2018).

[7] DCAD: Dietary cation anion difference, calculated as the sum of the dietary contents in K and Na minus the dietary

**Table S8: Forward and reverse primers used for real-time quantitative PCR in milk and blood samples to measure gene expression of antioxidant, immune or milk production functions.**

| Item | Matrix | Forward/Reverse | Primer sequence | Ref NCBI |
|---|---|---|---|---|
| ABCG2 | blood + milk | Forward | GGAGTCATGAAACCTGGCC | NM_001037478 |
| | | Reverse | GGATCCTTCCTTGCAGCTAA | NM_001037478 |
| AHR | blood + milk | Forward | GTGCAGAAAACTGTCAAGCCA | NM_001206026 |
| | | Reverse | AACATCTGGTGGGAAAGGCAG | NM_001206026 |
| AOAH | milk | Forward | AACGACAGCAACAAGATGGC | NM_001078096.2 |
| | | Reverse | AACACCCCAAATGCCGTTAC | NM_001078096.2 |
| BAX | milk | Forward | AGAAGCTGAGCGAGTGTCTGAA | NM_173894 |
| | | Reverse | CGCTCTCGAAGGAAGTCCAA | NM_173894 |
| CASP1 | milk | Forward | CTCCCACCTGGCAGGAATAC | XM_002692921 |
| | | Reverse | AGGAGCTGGAAAAGGAGGGA | XM_002692921 |
| CASP1P3 | milk | Forward | TCCGGACATTCAACAACCGT | NM_176638.5 |
| | | Reverse | ACCCACAATTCCCCACGATT | NM_176638.5 |
| CASP8 | milk | Forward | AATATTGGGGAGCAGCTGGG | NM_001045970.2 |
| | | Reverse | AGGCATCCTTGATGGGTTCC | NM_001045970.2 |
| CAT | blood + milk | Forward | AGATGGACACAGGCACATGA | NM_001035386.2 |
| | | Reverse | ACTGCCTCTCCATTTGCATT | NM_001035386.2 |
| CCL2 | blood + milk | Forward | GCTCGCTCAGCCAGATGCAA | NM_174006 |
| | | Reverse | GGACACTTGCTGCTGGTGACTC | NM_174006 |
| CCL20 | blood + milk | Forward | TTCGACTGCTGTCTCCGATA | NM_174263 |
| | | Reverse | GCACAACTTGTTTCACCCACT | NM_174263 |
| CCL5 | blood | Forward | CTGCCTTCGCTGTCCTCCTGATG | NM_175827 |
| | | Reverse | TTCTCTGGGTTGGCGCACACCTG | NM_175827 |
| CD14 | blood + milk | Forward | TCCACAGTCCAGCCGACAAC | NM_174008 |
| | | Reverse | AACGGCGCTAGACCAGTCAG | NM_174008 |
| CD36 | milk | Forward | CTGGCTGTGTTTGGAGGGATTC | NM_001278621 |
| | | Reverse | ACTGTCACTTCATCTGGATTCTGC | NM_001278621 |
| CDH1 | milk | Forward | TGCCCGACCATCAAAGAGAT | NM_174031 |
| | | Reverse | CGGTACGAAATGGCTTCCAC | NM_174031 |
| CLDN1 | milk | Forward | GAAGAACAGCACGTACACGGC | NM_001002763 |
| | | Reverse | CCTCTGGCAGAAGTCCATGGT | NM_001002763 |
| CLEC4A | milk | Forward | AAGACGACGAGGCACAGAAG | NM_001001854 |
| | | Reverse | AGCCCAGCCAATGAAGAGA | NM_001001854 |
| CLEC7A | milk | Forward | GAAGTTACCACCGTGCTTGC | NM_001191510.1 |
| | | Reverse | CCTCTGAAGTCATGCTGCGA | NM_001191510.1 |
| CSN1S1 | milk | Forward | AGGCAAGTGTCTTCCCAGC | NM_001031852.1 |
| | | Reverse | ACAACAAGGTGGAGCCATCC | NM_001031852.1 |
| CSN3 | milk | Forward | GTTCCCAATAGTGCTGAGGAACGACT | NM_181029.2 |
| | | Reverse | GGGTAGAAGTAGGCCAGTTCCTGAT | NM_181029.2 |
| CTSB | blood + milk | Forward | TGCAATGATGAAGAGTTTTTTCCTAG | NM_174294 |
| | | Reverse | GATTGGGATATATTTGGCTATTTTGT | NM_174294 |
| CXCL10 | blood + milk | Forward | TTCAGGCAGTCTGAGCCTAC | NM_001046551 |
| | | Reverse | ACGTGGGCAGGATTGACTTG | NM_001046551 |
| CXCL2 | blood + milk | Forward | GTGTCTCAACCCCGCCGCTC | NM_174299.3 |
| | | Reverse | TCCAGATGGCCTTAGGAGGTGG | NM_174299.3 |
| CXCL8 | blood + milk | Forward | TGAAGCTGCAGTTCTGTCAAG | NM_173925.2 |
| | | Reverse | TTCTGCACCCACTTTTCCTTGG | NM_173925.2 |
| DEFB5 | blood + milk | Forward | TCGTGCTCCTCTTCCTAGTC | NM_001130761 |
| | | Reverse | GGCACGAGATCGGAATACAG | NM_001130761 |
| FABP3 | blood + milk | Forward | GCGTTCTCTGTCGTCTTTCCC | NM_174313 |
| | | Reverse | CTGTGTTCTTGAAGGTGCTTTGTG | NM_174313 |
| FAS | blood + milk | Forward | GCCCACATGGCTGGTATCAA | NM_174662.2 |
| | | Reverse | TTTTTCCGTTTGCCAGGAGG | NM_174662.2 |
| GPX1 | blood + milk | Forward | GCAAGGTGCTGCTCATTGAG | NM_174076 |
| | | Reverse | CGCTGCAGGTCATTCATCTG | NM_174076 |
| GPX3 | blood + milk | Forward | GTCAACGTGGCCAGCTACTGA | NM_174077 |
| | | Reverse | CAGAATGACCAGACCAAATGGTT | NM_174077 |
| HSPA8 | milk | Forward | CGAATCATCAATGAGCCAACTG | NM_174345 |
| | | Reverse | TGCCACCCCCTAAATCAAAG | NM_174345 |
| IFNAR1 | blood + milk | Forward | TCCTTTGCCACGTGTCAAGT | NM_174552.2 |
| | | Reverse | AGTAGCGTGAGGGAGACAGA | NM_174552.2 |
| IFNAR2 | blood + milk | Forward | GTGTGGGTAAACACGACGGA | NM_174553 |
| | | Reverse | TGGGTCCAAAGGCTTGTCTG | NM_174553 |
| IFNB1 | milk | Forward | GACAGTCGTGCAAGTGCAAA | NM_174350.1 |
| | | Reverse | CAGTCACGGACGTAACCTGT | NM_174350.1 |
| IL10 | blood + milk | Forward | GTGATGCCACAGGCTGAGAA | NM_174088.1 |
| | | Reverse | TGCTCTTGTTTTCGCAGGGCAG | NM_174088.1 |
| IL1A | milk | Forward | CTGAAGAAGAGACGGTTGAG | NM_174092 |
| | | Reverse | ATGCATTCCTGGTGGATGAC | NM_174092 |
| IL1B | milk | Forward | CTCTCACAGGAAATGAACCGAG | EU276067 |
| | | Reverse | GCTGCAGGGTGGGCGTATCACC | EU276067 |
| IL2RA | blood + milk | Forward | AACACACAGATGCGCAGAAC | NM_174358 |
| | | Reverse | TTCACGTTCGTGTTCCCATG | NM_174358 |
| IL33 | milk | Forward | GATGGTGGCAGTCATCGGAA | NM_001075297.1 |
| | | Reverse | GTAGCTCCACAGAGTGCTCC | NM_001075297.1 |
| IL6 | blood + milk | Forward | TGCTGGTCTTCTGGAGTATC | EU276071 |
| | | Reverse | GTGGCTGGAGTGGTTATTAG | EU276071 |
| IRF3 | milk | Forward | GGAAGGATAAGCCCGACCTG | NM_001029845.3 |
| | | Reverse | GAGTCCTTGCTGTGGTCCTC | NM_001029845.3 |
| ITGA4 | blood + milk | Forward | GGGTTTGTAAGTCCAGCCTCA | NM_174748.1 |
| | | Reverse | CGGTGTTGATGACGTGGAAAG | NM_174748.1 |
| LALBA | milk | Forward | ACCAGTGGTTATGACACACAAGC | NM_174378 |
| | | Reverse | AGTGCTTTATGGGCCAACCAGT | NM_174378 |
| LAP | blood + milk | Forward | TGCTCCTTGCGCTCCTCTTC | NM_203435 |
| | | Reverse | CTCCGAGACAGGTGCCAATC | NM_203435 |
| LBP | milk | Forward | CCTGATTCTAGCATTCGACAG | NM_001038674 |
| | | Reverse | GCTGAAGTTCAGGCACG | NM_001038674 |
| LCN2 | milk | Forward | TGGCGGGGAATGCAATTAAG | XM_002691670 |
| | | Reverse | TAACAGGATGGAGGTGACGTTG | XM_002691670 |
| LPL | blood + milk | Forward | AGTTTATGAACTGGATGGCGGATG | NM_001075120 |
| | | Reverse | TCAGGAGAAAGGCGACTTGGAG | NM_001075120 |
| MMP9 | blood + milk | Forward | CGTTCCGACGACATGCTCTG | NM_174744 |
| | | Reverse | CATTGCCGTCCTGGGTGTAG | NM_174744 |
| MUC1 | milk | Forward | CTCTCCAGGCCATGATAGTG | NM_174115 |
| | | Reverse | AAGTGACCATGGAGCTTGAC | NM_174115 |
| NFE2L2 | blood + milk | Forward | AGGACATGGATTTGATTGAC | NM_001011678 |
| | | Reverse | TACCTGGGAGATGTTGGCA | NM_001011678 |
| NFKB1 | blood + milk | Forward | TGAGGCCATTGACGTGATCC | NM_001076409 |
| | | Reverse | TGCGGAAGGAGGTCTCTACA | NM_001076409 |
| NLRP3 | blood + milk | Forward | CTCAGTGGCAATACCCTGGG | NM_001102219.1 |
| | | Reverse | AGCACTGTCCCAACCACAAT | NM_001102219.1 |
| PLAU | milk | Forward | GCCAGGAGTCTACACAAGGG | NM_174147 |
| | | Reverse | TGGGGTCCTTCAGAGGACAA | NM_174147 |
| RIPK1 | milk | Forward | GCGAATCTCTCGGGTTGTGT | NM_001035012 |
| | | Reverse | GCCTGCTCCAGGAAGTCTG | NM_001035012 |
| S100A7 | blood + milk | Forward | CAGCTTGAGCAGGCCATTAC | NM_174596 |
| | | Reverse | CGTGGCTGTGGTTGTGATAG | NM_174596 |
| S100A8 | blood + milk | Forward | CTCCCTGATTGACGTCTACC | NM_001113725 |
| | | Reverse | TCCAGGCCCACCTTTATCAC | NM_001113725 |

| | | | | |
|---|---|---|---|---|
| S100A9 | blood + milk | Forward | TGACACCCTGATCCAGAAAG | NM_001046328 |
| | | Reverse | GCCACCAGCATAATGAACTC | NM_001046328 |
| SCD | blood + milk | Forward | GCCTCTGCGGGTCTTCCTG | NM_173959 |
| | | Reverse | GGTGGGCACGGTGATCTCG | NM_173959 |
| SIRT1 | milk | Forward | ATACACTGGAGCAGGTT | NM_001192980 |
| | | Reverse | TTCATCAGCTGGGCATCTAG | NM_001192980 |
| SOCS3 | blood + milk | Forward | GATCCCTCTGGTGTTGAGCC | NM_174466.2 |
| | | Reverse | CAGCTGGGTGACTTTCTCGT | NM_174466.2 |
| SOD1 | blood + milk | Forward | TGTTGCCATCGTGGATATTGTAG | NM_174615 |
| | | Reverse | CCCAAGTCATCTGGTTTTTCATG | NM_174615 |
| SOD2 | blood + milk | Forward | GCAATTCCCCTTGGGGTTCT | NM_201527.2 |
| | | Reverse | CAGCCTGCGTTGAAGTTCAA | NM_201527.2 |
| SPARC | milk | Forward | GAGAATTCGATGATGGTGC | NM_174464 |
| | | Reverse | GAAGGTCTTGTTGTCGTTG | NM_174464 |
| STAT1 | blood + milk | Forward | CAAAGGAAGCCCCAGAGCCTAT | NM_001077900.1 |
| | | Reverse | GCCACTCTTCTGTGTTCACTTAC | NM_001077900.1 |
| STAT2 | blood + milk | Forward | AGCCCGTTTCAGGATCAGC | NM_001205689.1 |
| | | Reverse | CAGTGCAGCTTTCTGCCAGTTC | NM_001205689.1 |
| STAT3 | blood + milk | Forward | GACCGGTGTCCAGTTCACAA | NM_001012671 |
| | | Reverse | AAATTTCCGGGACCCTCTGA | NM_001012671 |
| STAT4 | blood + milk | Forward | ACAGCAAATCGCCTGCATTG | NM_001083692 |
| | | Reverse | CTCCAACTGCCGTCTGAGTT | NM_001083692 |
| STAT5A | blood + milk | Forward | CATGTACCCACAGAACCCTGAC | NM_001012673.1 |
| | | Reverse | GGGAGAGAGGGCTCCAGACT | NM_001012673.1 |
| TAP | blood + milk | Forward | GTAGGAAATCCTGTAAGCTGTG | AF014106 |
| | | Reverse | GTGTCTTGGCCTTCTTTTAC | AF014106 |
| TJP1 | milk | Forward | AATGCATCCTGACCACCAGG | XM_024982002 |
| | | Reverse | GATGGTGCCGGGTTTGTTTC | XM_024982002 |
| TLR2 | blood + milk | Forward | ACTGGGTGGAGAACCTCATGGTCC | NM_174197.2 |
| | | Reverse | ATCTTCCGCAGCTTACAGAAGC | NM_174197.2 |
| TLR4 | blood + milk | Forward | GCATGGAGCTGAATCTCTAC | NM_174198.6 |
| | | Reverse | CAGGCTAAACTCTGGATAGG | NM_174198.6 |
| TNFA | blood + milk | Forward | TCTTCTCAAGCCTCAAGTAACAAGC | NM_173966.3 |
| | | Reverse | CCATGAGGGCATTGGCATAC | NM_173966.3 |
| TXNRD1 | blood + milk | Forward | GACCAAACCATCGAAGGAGAGTAT | NM_174625 |
| | | Reverse | CTCGTGCAAGCATCTCTTCCT | NM_174625 |
| ACTB | blood + milk | Forward | ACGGGCAGGTCATCACCATC | BT030480 |
| | | Reverse | AGCACCGTGTTGGCGTAGAG | BT030480 |
| GAPDH | blood + milk | Forward | GGCATCGTGGAGGGACTTATG | DQ403066 |
| | | Reverse | GCCAGTGAGCTTCCCGTTGAG | DQ403066 |
| PPIA | blood + milk | Forward | TCCGGGATTTATGTGCCAGGG | BC105173 |
| | | Reverse | GCTTGCCATCCAACCACTCAG | BC105173 |
| RPLP0 | blood + milk | Forward | CAACCCTGAAGTGCTTGACAT | NM_001012682 |
| | | Reverse | AGGCAGATGGATCAGCCA | NM_001012682 |
| SLC2A1 | blood | Forward | | |
| | | Reverse | | |
| SELL | blood | Forward | | |
| | | Reverse | | |
| IL23A | blood | Forward | | |
| | | Reverse | | |
| CD80 | blood | Forward | | |
| | | Reverse | | |
| MMP14 | blood | Forward | | |
| | | Reverse | | |
| CCR6 | blood | Forward | | |
| | | Reverse | | |
| CX3CR1 | blood | Forward | | |
| | | Reverse | | |
| CD86 | blood | Forward | | |
| | | Reverse | | |
| TNFRSF18 | blood | Forward | | |
| | | Reverse | | |
| FAS | blood + milk | Forward | | |
| | | Reverse | | |
| CCR7 | blood | Forward | | |
| | | Reverse | | |
| CXCR4 | blood | Forward | | |
| | | Reverse | | |
| SPI1 | blood | Forward | | |
| | | Reverse | | |
| HMOX1 | blood | Forward | | |
| | | Reverse | | |
| C5AR1 | blood | Forward | | |
| | | Reverse | | |
| SOD3 | blood + milk | Forward | | |
| | | Reverse | | |
| CD74 | blood | Forward | | |
| | | Reverse | | |
| NQO1 | blood | Forward | | |
| | | Reverse | | |
| CCR1 | blood | Forward | | |
| | | Reverse | | |
| BCL2 | blood + milk | Forward | | |
| | | Reverse | | |
| ELANE | blood | Forward | | |
| | | Reverse | | |
| ACOX2 | blood + milk | Forward | | |
| | | Reverse | | |
| SIRPA | blood | Forward | | |
| | | Reverse | | |
| SOCS2 | blood | Forward | | |
| | | Reverse | | |
| IKZF1 | blood + milk | Forward | | |
| | | Reverse | | |
| RORa | blood | Forward | | |
| | | Reverse | | |
| ADGRE1 | blood + milk | Forward | | |
| | | Reverse | | |
| BIRC5 | blood | Forward | | |
| | | Reverse | | |
| NFkB2 | blood + milk | Forward | | |
| | | Reverse | | |
| YWHAZ | blood + milk | Forward | | |
| | | Reverse | | |
| CXCL1 | blood + milk | Forward | | |
| | | Reverse | | |
| OCLN | milk | Forward | | |
| | | Reverse | | |
| IGFBP5 | milk | Forward | | |
| | | Reverse | | |
| RPL19 | blood + milk | Forward | | |
| | | Reverse | | |
