## Supplementary figures for "Nutritional vitamin E or plant extracts affect the immune response and mammary epithelium integrity during intramammary lipopolysaccharide challenge in early lactation"

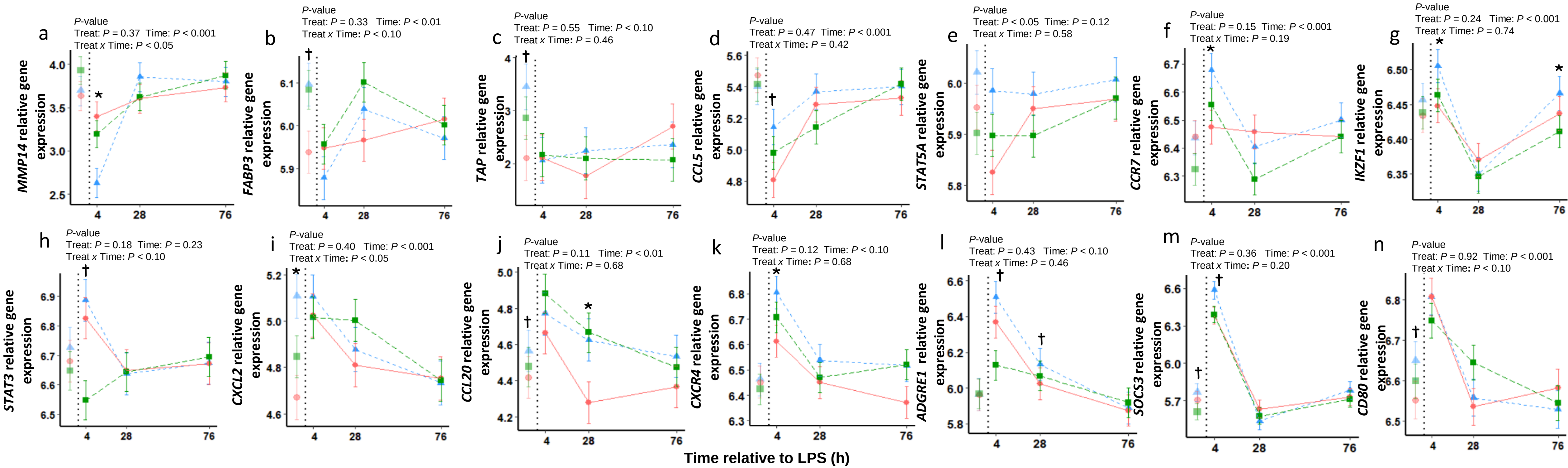


Figure S1: Abundance of blood mRNA determined by real-time quantitative PCR in control unsupplemented group (n = 11), vitamin E supplemented group (n = 13), and plant extract supplemented group (n = 12) dairy cows before and during LPS challenge times (4 h, 28 h, 76h). The expression of inflammatory response genes according to time and treatments: migration genes MMP14 (matrix metallopeptidase 14) (**a**), growth and proliferation FABP3 (fatty acid binding protein 3) (**b**), antimicrobial peptide TAP (Tracheal antimicrobial peptide) (**c**), CCL5 (C-C motif chemokine ligand 5) (**d**), STAT5A (signal transducer and activator of transcription 5A) (**e**), CCR7 (C-C motif chemokine receptor 7) (**f**), transcription factor IKZF1 (family zinc finger 1) (**g**), STAT3 (signal transducer and activator of transcription 3) (**h**), CXCL2 (C-X-C motif chemokine ligand 2) (**i**), CCL20 (C-C motif chemokine ligand 20) (**j**), CXCR4 (C-C motif chemokine receptor 4) (**k**), ADGRE1 (adhesion G protein-coupled receptor E1) (**l**), SOCS3 (suppressor of cytokine signaling 3) (**m**), CD80 (CD80 antigen) (**n**). Control group is represented in solid red line (——●——), vitamin E group in dotted green line (--■--), and plant extract group in dotted blue line (- -▲- -). Expression of these relative genes was calculated according to the decimal logarithm of the geometric mean of three reference genes ACTB (actin beta), YWHAZ (tyrosine 3-monooxygenase/tryptophan 5-monooxygenase activation protein zeta), and RPLP0 (Ribosomal Protein Lateral Stalk Subunit P0). Adjusted mean and SEM are represented and data were analyzed according to a mixed model. Significant differences between treatment (Treat) and time (Time) and their interaction (Treat x Time) were noted above the graphs. The significance thresholds were set at ∗ $P < 0.05$ and trends were noted at † $P \leq 0.10$.

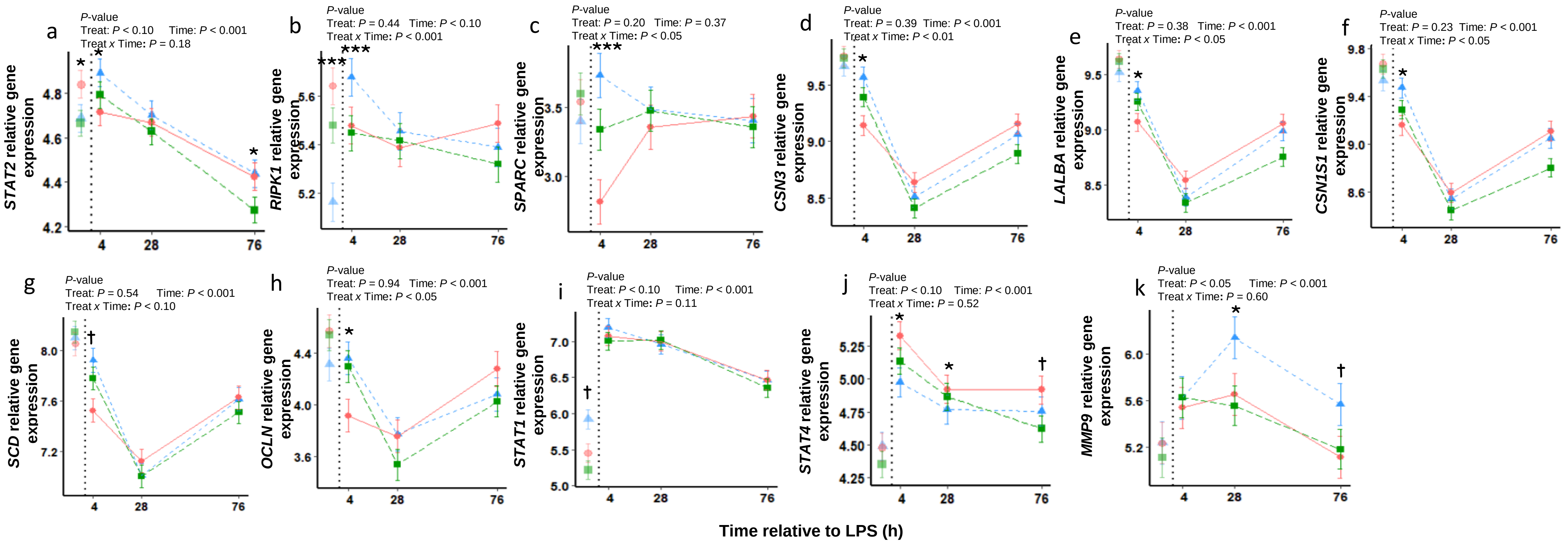


Figure S2: Abundance of milk mRNA determined by real-time quantitative PCR in control unsupplemented group (n = 11), vitamin E supplemented group (n = 13), and plant extract supplemented group (n = 12) dairy cows before and during LPS challenge times (4 h, 28 h, 76h). The expression of inflammatory response and milk synthesis genes according to time and treatments: STAT2 (signal transducer and activator of transcription 2) (**a**), necroptosis RIPK1 (receptor interacting serine/threonine kinase 1) (**b**), SPARC (secreted protein acidic and cysteine rich) (**c**), CSN3 (casein κ) (**d**), Lactose synthesis LALBA (lactalbumin α) (**e**), CSN1S1 (casein alpha s1) (**f**), SCD (stearoyl-CoA desaturase) (**g**), cell junction OCLN (occluding) (**h**), STAT1 (signal transducer and activator of transcription 1) (**i**), STAT4 (signal transducer and activator of transcription 4) (**j**), MMP9 (matrix metallopeptidase 9) (**k**). Control group is represented in solid red line (——●——), vitamin E group in dotted green line (--■--), and plant extract group in dotted blue line (- -▲- -). Expression of these relative genes was calculated according to the decimal logarithm of the geometric mean of three reference genes ACTB (actin beta), YWHAZ (tyrosine 3-monooxygenase/tryptophan 5-monooxygenase activation protein zeta), and PPIA (Cyclophilin A). Adjusted mean and SEM are represented and data were analyzed according to a mixed model. Significant differences between treatment (Treat) and time (Time) and their interaction (Treat x Time) were noted above the graphs. The significance thresholds were set at ∗ P < 0.05, ∗∗∗ P < 0.001 and trends were noted at † P ≤ 0.10.

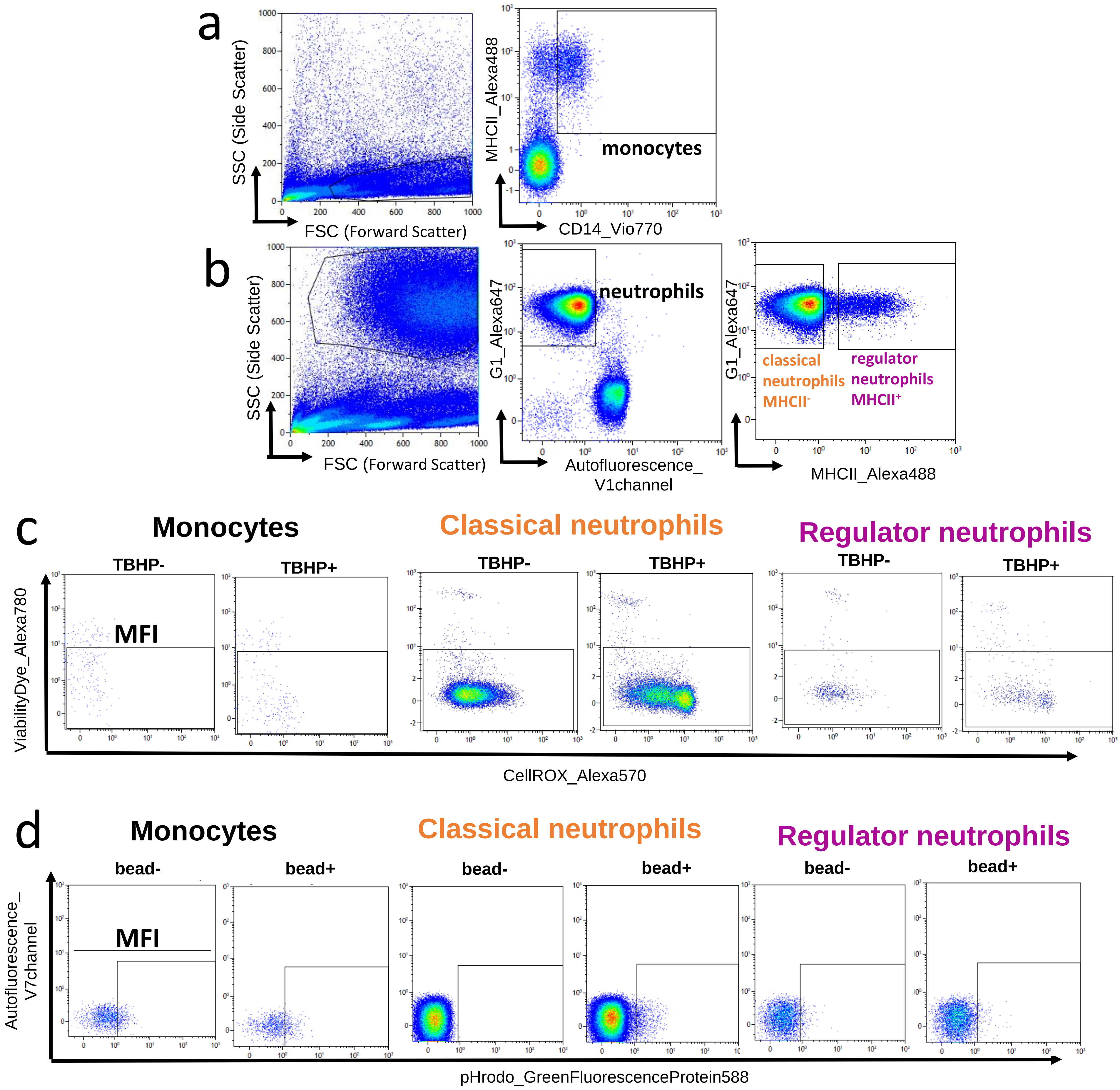


Figure S3: Flow cytometry-gating strategy for the identification of monocytes ($MCHII^+$ and $CD14^+$) (**a**) neutrophils ($G1^+MHCII^-$ or $G1^+MHCII^+$) in the blood, and measurement of reactive oxygen species (ROS) production and phagocytosis by these immune cells. The neutrophils was identified using FSC and SSC parameters, then with anti-G1 antibody and a channel free of markers to differentiate them from eosinophils using autofluorescence (Rambault et al., 2023). Two types of neutrophils were identified with the anti-MHCII antibody: classical neutrophils ($MHCII^-$) and regulatory neutrophils expressing MHCII ($MHCII^+$) in the same graph (**b**). *Ex vivo* reactive oxygen species (ROS) production was measured with CellROX Orange and dye among live cells (viability dye). Cells were stimulated with (TBHP +) or without (TBHP -) tert-butyl hydroperoxide to stimulate the levels of ROS production. Mean fluorescence intensities of CellROX_Alexa570 of the gated was used for data (**c**). *Ex vivo* phagocytosis was measured with beads of pHrodo *Escherichia coli* BioParticles in intracellular. Phagocyte cells were measured with (bead +) or without (bead -) to verify the phagocytose level signal. Total Mean fluorescence intensities of the graph with pHrodo_GreenFluorescenceProtein588 signal was used for data (**d**).

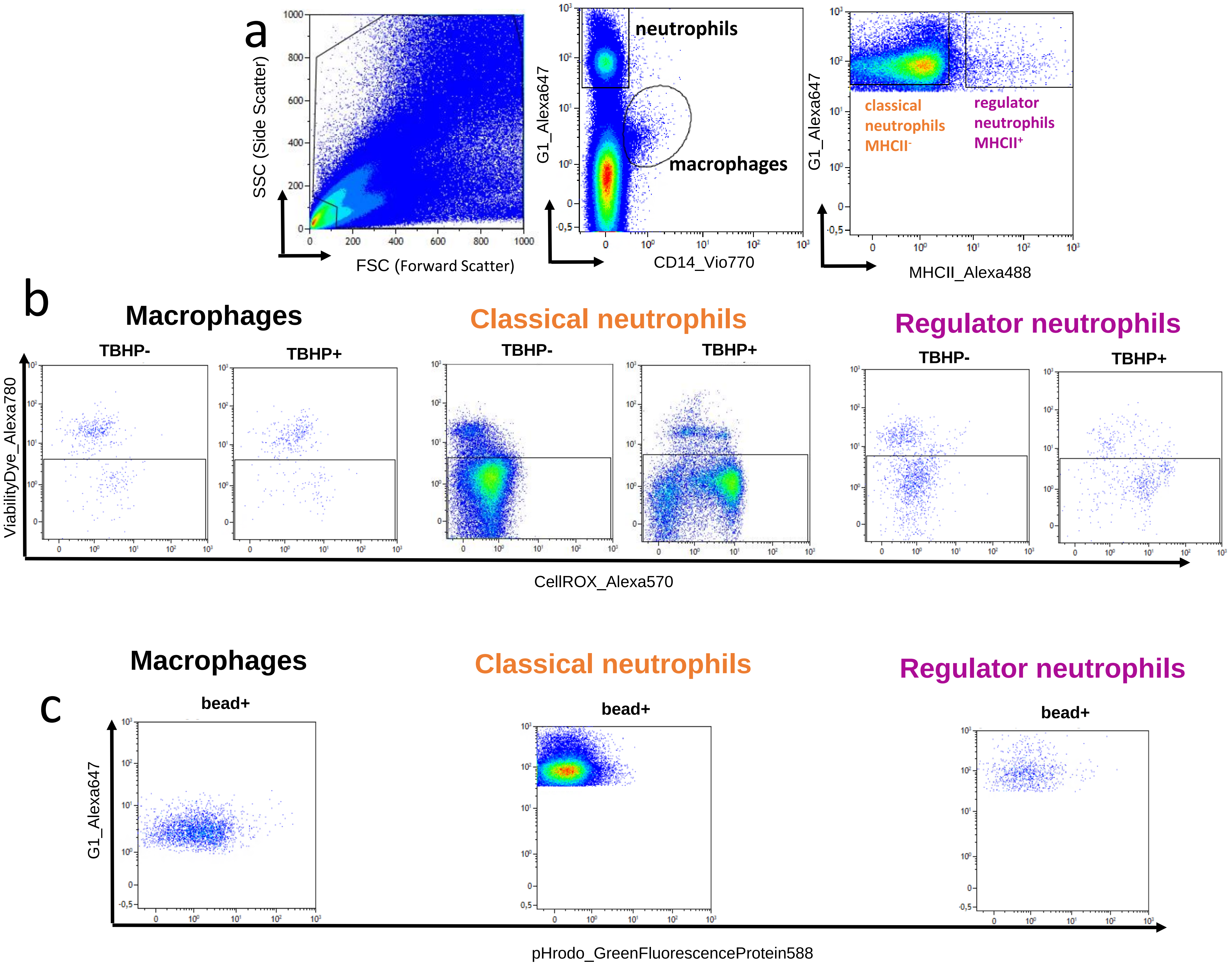


Figure S4: Flow cytometry-gating strategy for the identification of monocytes (MCHII$^+$ and CD14$^+$) neutrophils (G1$^+$MHCII$^-$ or G1$^+$MHCII$^+$) in the milk, and measurement of reactive oxygen species (ROS) production and phagocytosis by these immune cells. The neutrophils were identified with the G1 marker and macrophages with the CD14 marker on the same graph. Two types of neutrophils were identified with the anti-MHCII antibody: classical neutrophils (MHCII$^-$) and regulatory neutrophils expressing MHCII (MHCII$^+$) in the same graph (**a**). *Ex vivo* reactive oxygen species (ROS) production was measured with CellROX Orange and dye among live cells (viability dye). Cells were stimulated with (TBHP +) or without (TBHP -) tert-butyl hydroperoxide to stimulate the levels of ROS production. Mean fluorescence intensities of CellROX_Alexa570 of the gated was used for data (**d**). *Ex vivo* phagocytosis was measured with beads of pHrodo *Escherichia coli* BioParticles in intracellular. Phagocyte cells were measured with (bead +) or without (bead -) to verify the phagocytose level signal. Total Mean fluorescence intensities of the graph with pHrodo_GreenFluorescenceProtein588 signal was used for data (**c**).
